# Targeted degradation of GSPT1 and NEK7 by a molecular glue prodrug for treatment of HCC

**DOI:** 10.1101/2025.06.25.661504

**Authors:** Przemysław Glaza, Roman Pluta, Krzysztofa E. Odrzywół, Marta Klejnot, Maria Wieczorek, Sylvain Cottens, Donald Coppen, Paweł Dobrzański, Tomas Drmota, Joanna Lis-Grześniak, Agata Śnieżewska, Joanna Majkut, Martyna Mianowska, Paulina Rozborska, Marta Jarmuszkiewicz, Katarzyna Kaczanowska, Aleksandra Adamska, Toshimitsu Takagi, Anna Sawicka, Anna Serwotka-Suszczak, Olga Makowska, Daria Gajewska, Kinga Jurczak, Kinga Leszkowicz, Michał Mankiewicz, Kamil Przytulski, Janusz Wiśniewski, Anna Szlachcic, Michał J. Walczak

## Abstract

Targeted Protein Degradation (TPD) technology, in the form of CRBN-modulating molecular glues, offers numerous unprecedented therapeutic benefits as evidenced by the success of approved high-value immunomodulatory imide drugs (IMiDs) such as lenalidomide and pomalidomide. Building upon these successes, we employed a small CRBN-focused library of molecular glues in a phenotypic screen against hepatocellular carcinoma (HCC) cell lines. While the original library was primarily designed to target SALL4, we identified additional CRBN substrates, including GSPT1, NEK7, and CK1*α*, whose degradation potently induced cell death in HCC cell lines. Subsequent lead optimization efforts yielded a compound, ABS-752, which demonstrated superior *in vitro* and *in vivo* activity through the potent degradation of GSPT1. Notably, ABS-752 does not form ternary complexes with CRBN and the neosubstrates. Further investigations revealed that ABS-752 is a prodrug activated by the monoamine oxidase, VAP-1, to an aldehyde intermediate and subsequently to the active molecule, ABT-002. VAP-1, which is overexpressed in cirrhotic liver, was identified as the primary monoamine oxidase responsible for the conversion of ABS-752. ABS-752 is currently in clinical trials for the treatment of HCC.

## Introduction

Hepatocellular carcinoma (HCC) is a leading cause of cancer death, with estimated 2022 figures of 736,000 new cases and 645,000 deaths, projected to rise to 1.33 million and 1.21 million respectively by 2050.^1,2^ While traditional causes such as HCV, HBV, alcohol, and aflatoxins are declining, metabolic diseases are increasingly driving new cases.^3–5^ In the US, metabolic disorders accounted for a larger share of HCC cases (36-37%) than HCV, alcohol, and HBV combined in 2007-2011.^6,7^ This trend is mirrored in the EU.^8^

Curative HCC treatment (resection/transplant) is limited to early-stage cases. Most patients are diagnosed late and receive systemic therapies, either to downstage the disease or for palliative care.^9^ Current first-line therapies include bevacizumab plus atezolizumab, durvalumab plus tremelimumab, sorafenib, and lenvatinib; second-line options are cabozantinib, pembrolizumab, ramucirumab, and regorafenib. Despite these options, survival benefits are limited, highlighting the need for new targeted therapies.^10,11^

Promising HCC targets include GSPT1, NEK7, and SALL4. GSPT1, a translation termination factor, is often overexpressed in HCC and linked to tumor proliferation.^12–14^ NEK7 regulates the NLRP3 inflammasome and is also overexpressed in HCC, correlating with lower survival rates.^15^ SALL4 is a marker of poor prognosis in HCC.^16,17^

Molecular glue degraders using Cereblon (CRBN) have enabled the targeting of previously poorly tractable targets, including C2H2 zinc-finger transcription factors like Ikaros and Aiolos.^18–20^ SALL4, another such factor, is potently degraded by IMiDs, making it a focus for developing novel CRBN-binding molecules.^21,22^

## Results

### Identification of CRBN-based compounds that reduce the viability of Hep3B but not a multiple myeloma cell line

Glutarimide-containing compounds were screened against the Hep3B cell line to identify CRBN-based molecular glues cytotoxic to HCC cell lines. The H929 B-cell line was used as a counter-screen for general cytotoxicity. Both cell lines were incubated with compounds for 72 hours, and cell viability was analyzed. Compounds impairing Hep3B viability by >50% (relative to vehicle control) with no significant effect on H929 cells were selected (Fig. 1a). AAC-215 was chosen for further evaluation. AAC-215’s demonstrated significant potency (IC50 < 100 nM) and efficacy (mean minimum viability = 25 ± 10%) against Hep3B, and a minimal effect on H929 cells (mean minimum viability 98 ± 11%) (Fig. 1a; Supplementary Table S1).

**Figure 1.**
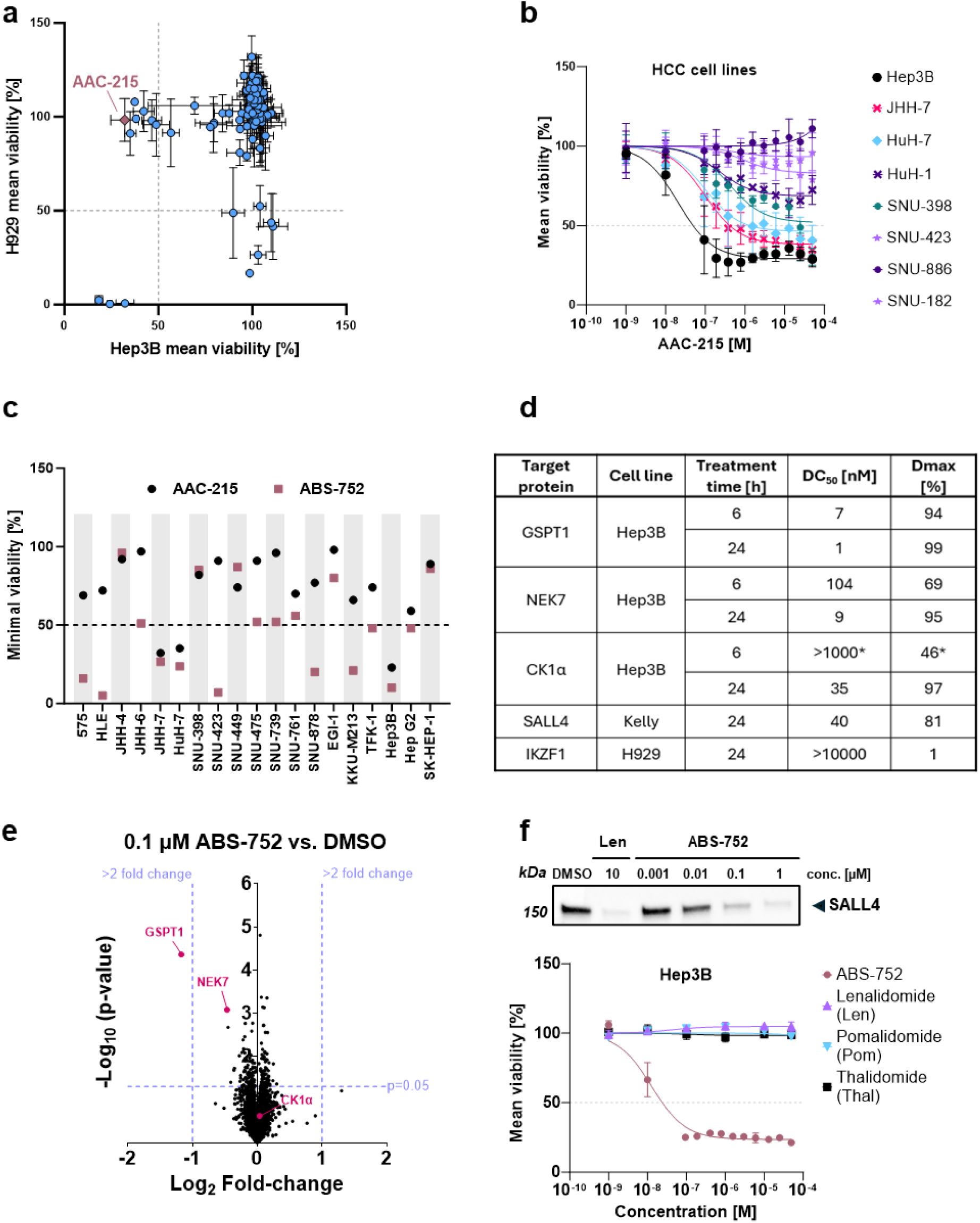
Compounds AAC-215 and ABS-752 impair the viability of HCC cell lines. **(a)** Hep3B and H929 cell line viability values determined using the CellTiter-Glo (CTG) assay in the presence of collection of molecular glues (at single concentration of 13 µM). The standard deviations shown were calculated based on at least two biological replicates **(b)** Representative CTG viability results for HCC cell lines treated with AAC-215 in a dose-dependent manner (compound concentration range 1 nM – 50 µM). Presented data are mean with standard deviation. **(c)** The AAC-215 derivative, ABS-752 shows higher cytotoxic/cytostatic activity in HCC cell lines compared to parent compound: ABS-752 affects 13 out of 19 HCC cell lines as measured by a min. of 50% of viability drop compared with 3 out 19 cell lines for AAC-215 (Charles River Laboratories cell line collection; JHH-7 and HuH-7 cell lines were tested in CTG assay in-house). **(d)** Degradation coefficients (DC_50_) determined for ABS-752-mediated protein degradation monitored by Western blot after 6 or 24-hour treatment for different cell-lines. *Values represent the mean from 3 independent experiments analyzing CK1α degradation at 1 µM ABS-752. **(e)** Proteomic analysis of ABS-752-induced changes in protein abundance in Hep3B cells. Cells were treated for 6 hours with DMSO and 0.1 µM ABS-752, total of 6218 proteins were identified. **(f)** Representative CTG viability results for Hep3B line treated with compounds in a dose-dependent manner (compound concentration range 1 nM – 50 µM). Western blot results for SALL4 degradation in Kelly cell line in the presence of Lenalidomide (Len) and various concentration of ABS-752.

### Profiling of AAC-215 and its derivatives against an expanded panel of HCC and non-HCC cell lines

AAC-215 was tested against a panel of HCC cell lines (HuH-1, HuH-7, JHH-7, SNU-182, SNU-398, SNU-423, and SNU-886), yielding varied results. JHH-7 and HuH-7 were most susceptible (minimum viabilities <50%), but less sensitive than Hep3B. Other cell lines were unresponsive (e.g., SNU-182, SNU-423, SNU-886) or showed moderate effects (HuH-1, SNU-398) (Fig. 1b; Supplementary Table S1). A chemical derivatization program led to ABS-752, which is more potent against Hep3B and also profoundly impacts viability in 575, HLE, JHH-7, HuH-7, SNU-423, SNU-878, and KKU-M213 cell lines (Fig. 1c).

To assess the selectivity of compound ABS-752 for HCC cell lines, we analyzed the GSPT1 protein degradation after 24 hours in several non-HCC cell lines: KG-1, H1155, MOLM-13, H929, and K562. Additionally, we examined NEK7 degradation at the same time point for the KG-1 and H1155 lines (Supplementary Fig. S2a-g). Strong degradation of GSPT1 was observed only in the KG-1 line. For the other tested cell lines, ABS-752 did not cause a significant change in GSPT1 levels.

Among the cell lines tested in the degradation assay, KG-1, MOLM-13, and H929 were selected for viability assays to determine if GSPT1 degradation, or its absence, correlated with cell survival under ABS-752 treatment. The panel of non-HCC cell lines in the CTG experiment was also enriched with Kasumi-1, Kelly, and MDS-L cell lines (Supplementary Fig. S2h). The observed high variability in response in both HCC and non-HCC cancer cell lines indicates that there is no correlation between cell line type and sensitivity to ABS-752 degrader treatment.

### Identification of relevant neosubstrates degraded by ABS-752

We investigated ABS-752’s interaction with CRBN neosubstrates using biochemical assays and cellular Western blots. After 24 hours of ABS-752 treatment, we analyzed the levels of SALL4, IKZF1, CK1α, NEK7, and GSPT1. We also examined the degradation of GSPT1, NEK7, and CK1α after a shorter, 6-hour treatment. Hep3B cells were used for CK1α, GSPT1, and NEK7 degradation assays. Due to low IKZF1 and SALL4 expression in Hep3B, H929 and Kelly cell lines were used for IKZF1 and SALL4 analysis^23,24,25^, respectively (Supplementary Fig. S1).

ABS-752 degraded SALL4, NEK7, CK1α, and GSPT1, but not IKZF1 (unlike Pomalidomide in H929 cells^24^). Degradation coefficients (DC_50_) determined for ABS-752-mediated neosubstrates degradation monitored by Western blot after 6 or 24-hour treatment is presented in Fig. 1d. Analysis of degradation parameters across the two time points indicates that GSPT1 is the primary target of ABS-752. This is supported by its near-complete degradation (DC_50_ < 10 nM) observed after only 6 hours of treatment. NEK7 and CK1α showed much weaker degradation after 6 hours compared to GSPT1, with DC_50_ values of approximately 100 nM and >1000 nM, respectively.

Proteomic analysis of Hep3B cells treated with 0.1 µM ABS-752 for 6 hours confirmed GSPT1 as the most preferentially degraded target. The analysis also revealed greater degradation of NEK7 compared to CK1α, for which no downregulation in the experiment was observed (Fig. 1e). The levels of SALL4 and IKZF1 proteins in Hep3B cells were below the detection limit for quantification. Taking together, Western blot and proteomics measurements of target protein degradation at the 6-hour time point show good correlation.

### Targeting GSPT1 is crucial for Hep3B proliferation inhibition

To determine the effect of ABS-752-mediated SALL4 degradation on Hep3B viability, we used known SALL4 degraders like Lenalidomide, Pomalidomide, and Thalidomide as controls.^25,26^ These IMiDs did not degrade GSPT1 in Western blot (data not shown) and did not score in our HCC cell line CTG screen (Fig. 1f). We confirmed Lenalidomide’s potent SALL4 degradation in Kelly and Hep3B cell lines. ABS-752 degraded SALL4 comparably to Lenalidomide (Supplementary Fig. S3), while only ABS-752 impaired Hep3B viability (Fig. 1f), suggesting that SALL4 degradation alone does not explain ABS-752’s activity.

To assess GSPT1’s role, we created a Hep3B cell line with a CRISPR-engineered GSPT1 degradation-resistant mutant (G575N). ABS-752 did not degrade the mutant GSPT1, nor did it affect the mutant cell line’s viability (Fig. 2a-b), strongly indicating GSPT1 degradation drives ABS-752’s activity.

**Figure 2.**
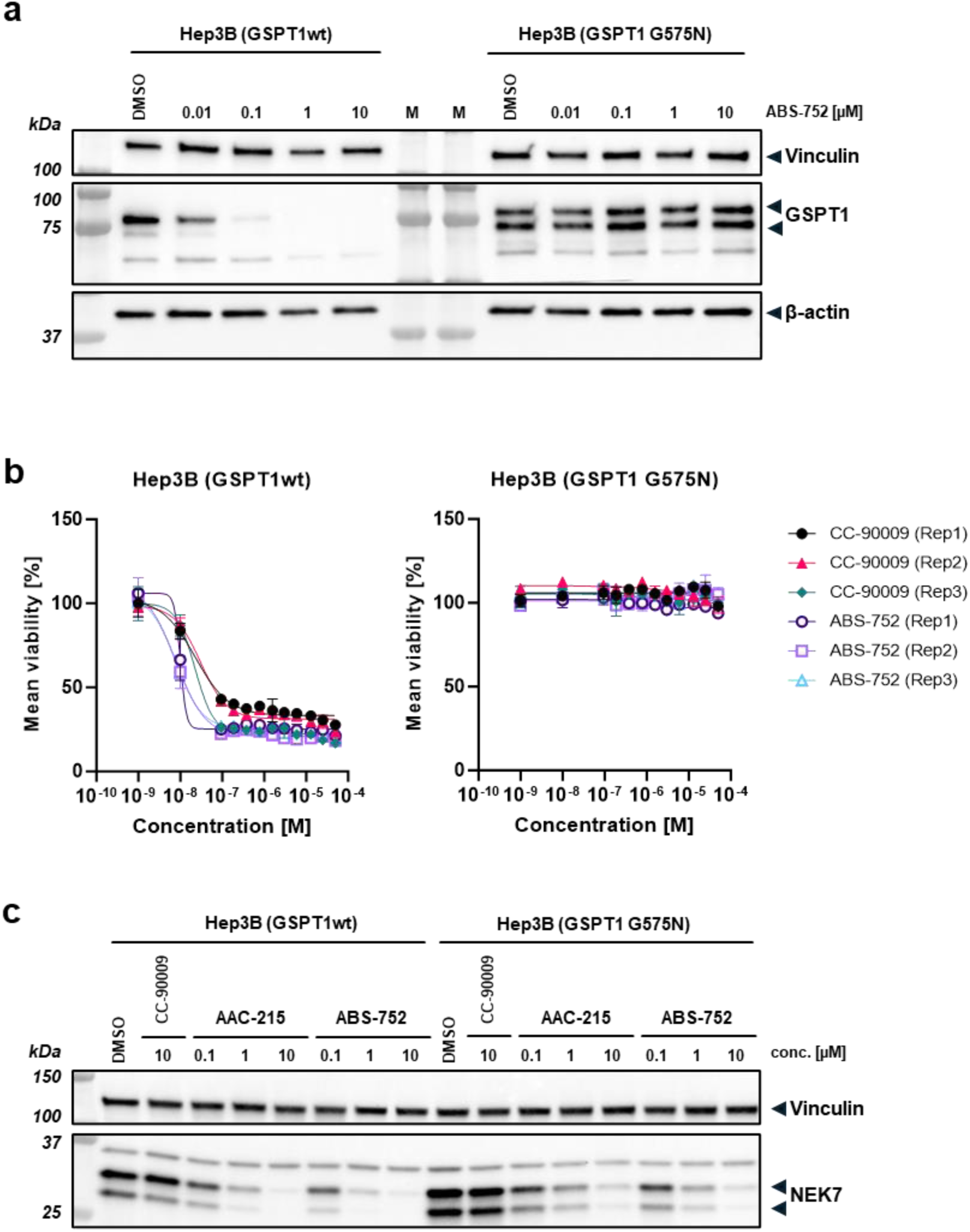
ABS-752-dependent degradation leads to the reduced viability of the Hep3B cell line. **(a)** Representative Western blot results for GSPT1 degradation in the presence of various concentrations of ABS-752 in wild type Hep3B line and Hep3B expressing GSPT1 G575N. The abundance of target proteins was analyzed after 48-hour incubation with compounds. The molecular weight of corresponding standards is indicated on the left (M wells). **(b)** CTG viability results for Hep3B cell line expressing wild type GSPT1 and GSPT1 G575N variant. The results of three independent replicates are presented for both cell lines (compound concentration range 1 nM – 50 µM). Data points are means with standard deviation (technical duplicates). **(c)** Representative Western blot results for NEK7 degradation in wild type and mutated Hep3B cell line after 24-hour treatment with ABS-752. Compounds CC-90009 and AAC-215 were used as controls.

Using the Hep3B GSPT1 G575N cell line, we confirmed ABS-752-mediated NEK7 degradation is GSPT1-independent. CC-90009, a GSPT1 degrader that doesn’t recruit NEK7, did not affect NEK7 levels in wild-type Hep3B cell line. However, both ABS-752 and AAC-215 caused dose-dependent NEK7 reduction in Hep3B cells, regardless of GSPT1 mutation status (Fig. 2c).

### Discovery of ABS-752 metabolic conversion

To determine if protein degradation results from CRBN target recruitment, we biochemically measured ABS-752-induced proximity between degron/target and CRBN. AlphaLISA assays (1µM and 10µM ABS-752) showed an unexpected selectivity profile opposite to neosubstrate degradation (Fig. 3a-b; Supplementary Table S2). Only SALL4 showed concordance between AlphaLISA recruitment and degradation in Kelly cells. While CC-90009 (a known GSPT1 degrader^27^) scored positive in our ternary complex formation assays (Fig. 3c; Supplementary Table S2), ABS-752, a potent Hep3B GSPT1 degrader, did not form a ternary complex with CRBN and GSPT1 biochemically.

**Figure 3.**
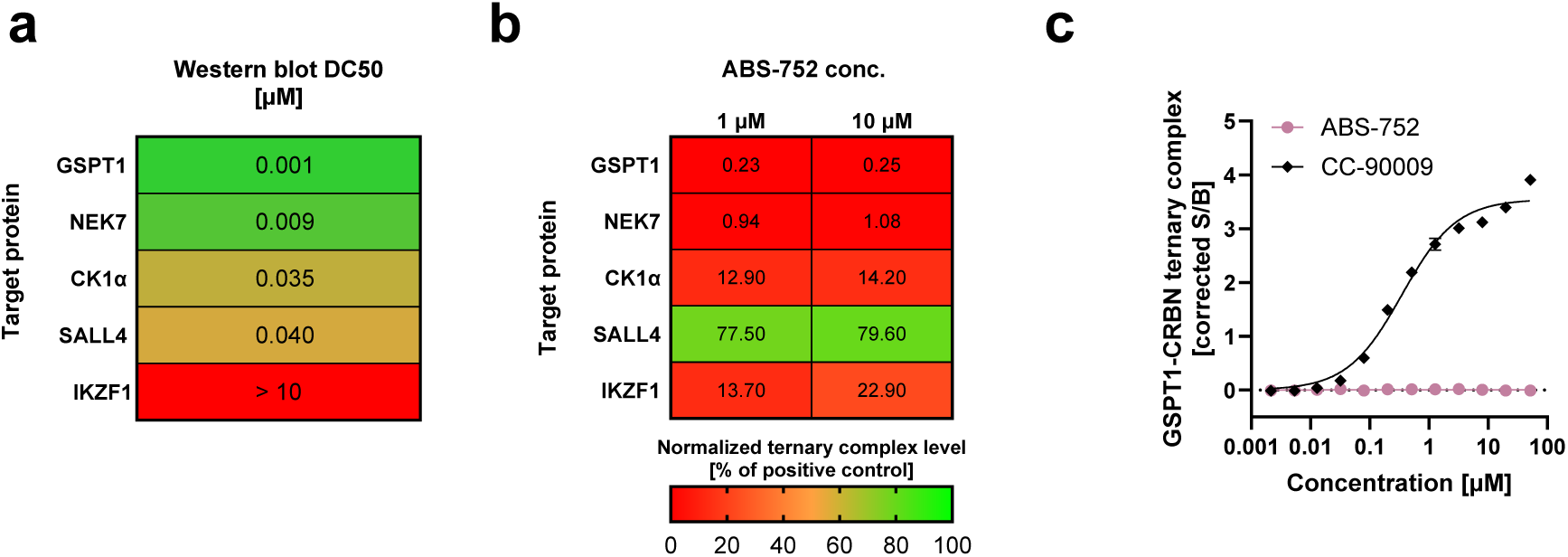
Discovery of ABS-752 metabolic conversion. **(a)** Degradation coefficients (DC_50_) determined for ABS-752-mediated protein degradation monitored by Western blot after 24-hour treatment. Dose-dependent target degradation curves are shown in Supplementary Fig. S4. **(b)** Normalized Target-glue-CRBN ternary complex response in AlphaLISA-based proximity assay. Presented values are mean from at least 3 independent experiments. **(c)** Representative results of TR-FRET-based assay for the formation of the GSPT1-CRBN ternary complex as a function of compound concentration (2 nM – 50 µM); presented data points are mean with standard deviation (technical triplicates).

### Conversion by VAP-1

ABS-752’s potent GSPT1 and NEK7 degradation, despite lacking biochemical recruitment of these targets, suggested in-cellulo conversion to an active compound. Metabolic profiling with CYP enzymes showed no significant conversions. Whole blood stability however, differed across species, with ABS-752 stable in human and rodent blood, but not others (Table 1).

**Table 1.** Stability of the compounds AAA-215 & ABS-752 in whole blood. Stability of ABS-752 in whole blood in the presence and absence of SSAO/VAP1 inhibitor PXS-4728A.

| Species | AAC-215 |  | ABS-752 |  |
| --- | --- | --- | --- | --- |
|  | % remaining @ 3 h | T1/2 [min] | % remaining @ 3 h | T1/2 [min] |
| Human (mixed gender) | >95 | >180 | >95 | >180 |
| Rat | 80 | >180 | 92.5 | >180 |
| Mouse | >95 | >180 | >95 | >180 |
| Dog | <1 | 7.3 | <1 | 21.4 |
| Monkey | <1 | 4.3 | <1 | 7.7 |
| Minipig | <1 | 3.5 | <1 | 15.5 |
| Rabbit | - | - | <1 | 11.9 |
| Blood: Human (mixed gender), Rat (male SD), Mouse (male CD-1), Dog (male Beagle), Monkey (male Cynomolgus), Minipig (male Gottingen), Rabbit (male New Zealand White), collected on sodium heparin or EDTA (Minipig). |  |  |  |  |
| Species | ABS-752 w/o inhibitor |  | ABS-752 with PXS-4728A |  |
|  | % remaining @ 3 h | T1/2 [min] | % remaining @ 3 h | T1/2 [min] |
| Dog | <1 | 19.7 | >95 | >180 |
| Monkey | <1 | 10.2 | 92.6 | >180 |
| Rabbit | <1 | 18.2 | 89.7 | >180 |
| Blood: Dog (male Beagle), Monkey (male Cynomolgus), Rabbit (male New Zealand White), collected on sodium heparin. The inhibitor was added 30 min before incubation of the compound. |  |  |  |  |

ABS-752’s primary amine suggested possible conversion to an aldehyde by amine oxidases. Potential ABS-752 metabolites were synthesized as standards (Fig. 4a). Identifying ABS-752’s conversion to an aldehyde led us to investigate the enzyme involved. Of three monoamine oxidases tested (MAO-A, MAO-B, VAP-1), only recombinant VAP-1 (mVAP-1) converted ABS-752 to ABT-971. This was confirmed by quenching aldehyde formation with the VAP-1 inhibitor PXS-4728A^28^ (Fig. 4a-b) and in whole blood (Table 1).

**Figure 4.**
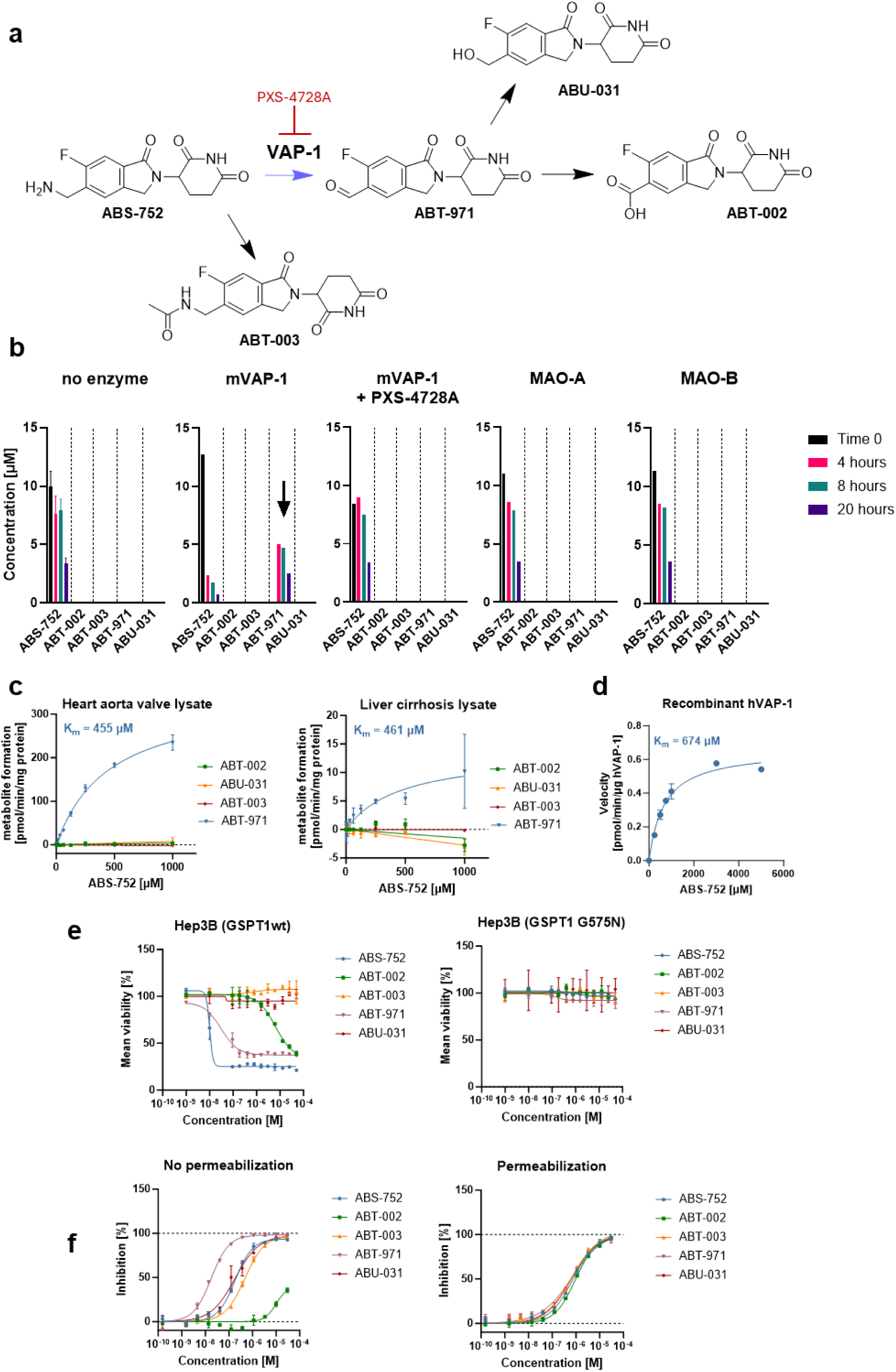
ABS-752 conversion to aldehyde by VAP-1 enzyme. **(a)** Structures and presumed mechanism of formation of ABS-752 metabolites. **(b)** Identification of the enzyme catalyzing the oxidation of compound ABS-752. Results of LC-MS/MS quantification of changes in the concentration of ABS-752 and its putative metabolites during incubation with and without recombinant monoamine oxidases (mouse VAP-1 (mVAP-1), human MAO-A and MAO-B). In the case of mVAP-1, metabolite formation was also checked in the presence of the VAP-1 inhibitor, PXS-4728A (0.25 µM). Data for the mixture with no enzyme are mean with standard deviation from two independent experiments, the remaining points are a single result. Legend indicates the time since the addition of ABS-752. **(c)** LC-MS/MS quantification results in the whole human tissue lysates for formation of presumed ABS-752 metabolites as a function of pro-drug concentration. ABS-752 was incubated with samples containing lysates of cirrhotic human liver or heart aorta for 1 hour at 37°C. Data shown are the mean with standard deviation for specific VAP-1 activity in lysates (based on 2 independently prepared samples). Presented Michaelis-Menten constants (K_m_) were determined based on formation rates of aldehyde, ABT-971. **(d)** Determination of ABS-752 deamination kinetics catalyzed by recombinant human VAP-1 monoamine oxidase. Observed ABS-752 oxidation rates as a function of pro-drug concentration. Data points are means with standard deviation from Amplex Red-based fluorogenic assay. **(e)** Representative CTG viability results for Hep3B cell line expressing wild type GSPT1 and GSPT1 G575N variant. Both cell lines were treated with ABS-752 and its presumed metabolites in a dose-dependent manner (compound concentration range 1 nM – 50 µM). In the case of Hep3B expressing GSPT1 G575N, compound ABT-002 was tested in the range of 1 nM - 25 µM. **(f)** Inhibition of intracellular Tracer/NanoLuc-CRBN complex formation by ABS-752 and its presumed metabolites. Representative inhibition dose-response curves, points are means with standard deviation calculated from technical duplicates.

VAP-1’s role in ABS-752 to ABT-971 conversion was verified in human lysates containing endogenous VAP-1. Cirrhotic liver (target tissue) and heart valve lysates were used. VAP-1 specific activity was determined by comparing metabolite levels with and without the VAP-1 inhibitor PXS-4728A. ABT-971 formation was observed in both lysates, with similar Km values for ABS-752 (461 µM and 455 µM, respectively). Recombinant human VAP-1 (hVAP-1) also oxidized ABS-752 to aldehyde (Amplex Red assay^29^), with a comparable Km (674 µM) (Fig. 4c). Aldehyde is not formed in the absence of human hVAP-1 (Supplementary Fig. S5).

### VAP-1 dependent activity of ABS-752 in cellular experiments

To confirm ABS-752’s VAP-1 dependence, Hep3B and KG-1 cell lines were treated with ABS-752 ± PXS-4728A. ABS-752 potently affected Hep3B viability (CTG viability assay) and degraded GSPT1/NEK7 (Western Blot) (Fig. 5a-b). The viability of the KG-1 line was also strongly reduced by ABS-752, with complete degradation of GSPT1 but with weak effect on NEK7 levels in this line (Supplementary Fig. S6a-b). PXS-4728A pre-treatment abolished ABS-752’s activity and target protein downregulation in Hep3B and KG-1 cell lines. CC-90009’s activity was unaffected by PXS-4728A (Fig. 5a-b).

**Figure 5.**
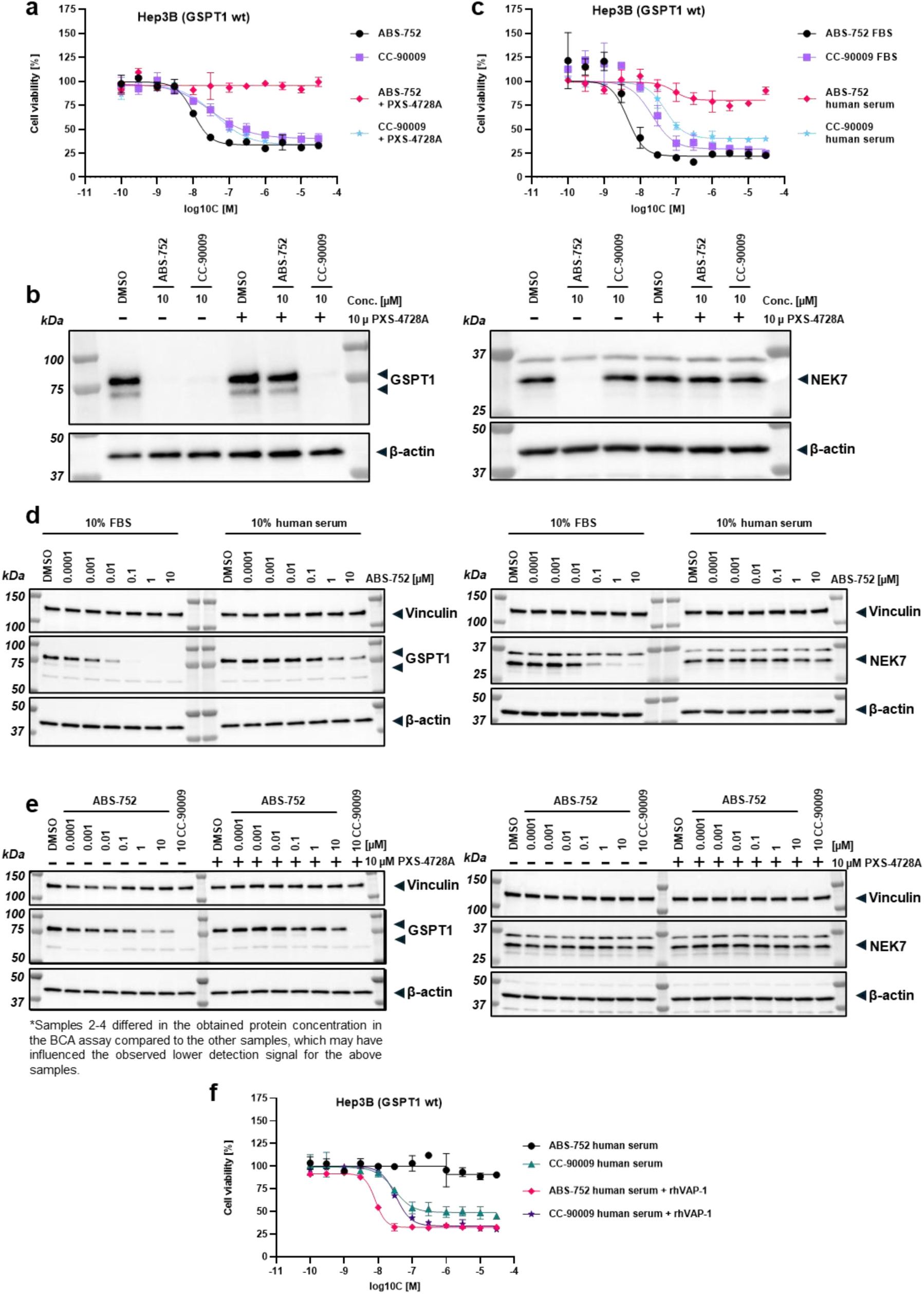
**(a)** CTG viability results for Hep3B cell line after 73-hour incubation (1 hour pre-treatment with 10 µM concentration of PXS-4728A or DMSO and 72 h co-treatment with tested compounds - ABS-752 and CC-90009 concentration range 0.1 nM – 30 µM). Data points are means with standard deviation (technical duplicates). **(b)** Western blot results for GSPT1 and NEK7 degradation in the presence or absence of 10 µM concentration of PXS-4728A in Hep3B line supplemented with 10% FBS. The molecular weight of corresponding standards is indicated on the left (M wells). **(c)** CTG viability results for Hep3B cell line supplemented with 10% FBS or 10% human serum after 72-hour treatment with tested compounds - ABS-752 and CC-90009 concentration range 0.1 nM – 30 µM. Data points are means with standard deviation (technical duplicates). **(d)** Western blot results for GSPT1 and NEK7 degradation after 24-hour post-treatment with ABS-752 and CC-90009 in Hep3B cell line supplemented with 10% FBS or 10% human serum. **(e)** Western blot results for GSPT1 and NEK7 degradation in the presence or absence of 10 µM concentration of PXS-4728A in Hep3B line supplemented with 10% human serum. **(f)** CTG viability results for Hep3B cell line supplemented with 10% human serum after 72-hour post-treatment with tested compounds (ABS-752 and CC-90009 concentration range 0.1 nM – 30 µM) in the presence or absence of 0.3 µM recombinant human VAP-1 protein. Data points are means with standard deviation (technical triplicates).

Humans express VAP-1 (AOC3), but not the SAO amine oxidase (AOC4) found in FBS.^30,31^ Hep3B and KG-1 cell lines were cultured in media with 10% FBS or 10% human serum (as a VAP-1 source) to assess cellular VAP-1 dependence. ABS-752 significantly reduced Hep3B and KG-1 viability in FBS-supplemented medium, but not in human serum-supplemented medium (pIC50 not achieved) (Fig. 5c, Supplementary Fig. S6b). In the case of Hep3B cells, ABS-752-mediated GSPT1 degradation was observed in both conditions, although lower with human serum than in FBS. ABS-752 induced prominent NEK7 downregulation in Hep3B cells with FBS, but this activity was significantly reduced with human serum (Fig. 5d). In contrast, a slight decrease in GSPT1 levels and no effect of ABS-752 on NEK7 levels were observed in KG-1 cells in the medium supplemented with human serum (Supplementary Fig. S6a). GSPT1 degradation in Hep3B cells cultured in medium supplemented with 10% human serum was confirmed to be VAP-1 dependent (Fig. 5e).

Addition of recombinant hVAP-1 (rhVAP-1) to Hep3B and KG-1 cell lines cultured in medium containing 10% human serum led to a significant reduction in the viability of both cell lines. This observation confirms VAP-1’s necessity for ABS-752 activity in cells (Fig. 5f, Supplementary Fig. S6b).

In addition, to investigate the activity of ABS-752 in normal human cells, CTG viability assay was conducted also in primary human hepatocytes. In contrast to Hep3B cells, ABS-752 showed no activity in primary human hepatocytes, even in the presence of FBS (Supplementary Fig. S7).

### Identification of ABT-002 as the active metabolite of ABS-752

To identify the active ABS-752 metabolite, Hep3B and its G575N variant were treated with ABS-752, ABT-002, ABT-003, ABT-971, and ABU-031 (Fig. 4e). Beyond the parent compound ABS-752, only ABT-002 and ABT-971 reduced viability in Hep3B cells expressing wild-type GSPT1. No inhibition was observed in the G575N line (Fig. 4e, Supplementary Table S4). ABT-002’s low CTG activity is due to limited cell membrane permeability, despite comparable CRBN affinity (Fig. 4f).

Ternary complex formation assays showed only ABT-002 formed a significant GSPT1-CRBN complex (Supplementary Fig. S8 and Table S2). ABT-002 was also the only compound to recruit NEK7.

### X-ray structure of GSPT1/ABT-002/CRBN complex

To understand ABT-002-driven GSPT1 recruitment to CRBN, we generated a crystal structure of human DDB1, CRBN, and GSPT1 complexed with ABT-002 (3.9 Å) (Fig. 6, Supplementary Table S5, PDB ID: 9HNE). The ligand binds as expected, with the glutarimide ring in the tryptophan cage and the isoindolinone in the canonical position^32^. CRBN interacts with GSPT1’s domain 3 via its thalidomide-binding domain, extending towards the N-terminal LON domain. ABT-002’s fluorine interacts with CRBN HIS353 and LYS572. The CRBN-GSPT1 interface is stabilized by GSPT1 GLN351’s interaction with ABT-002’s carboxyl group, which is positioned by hydrogen bonding to CRBN GLU377.

**Figure 6.**
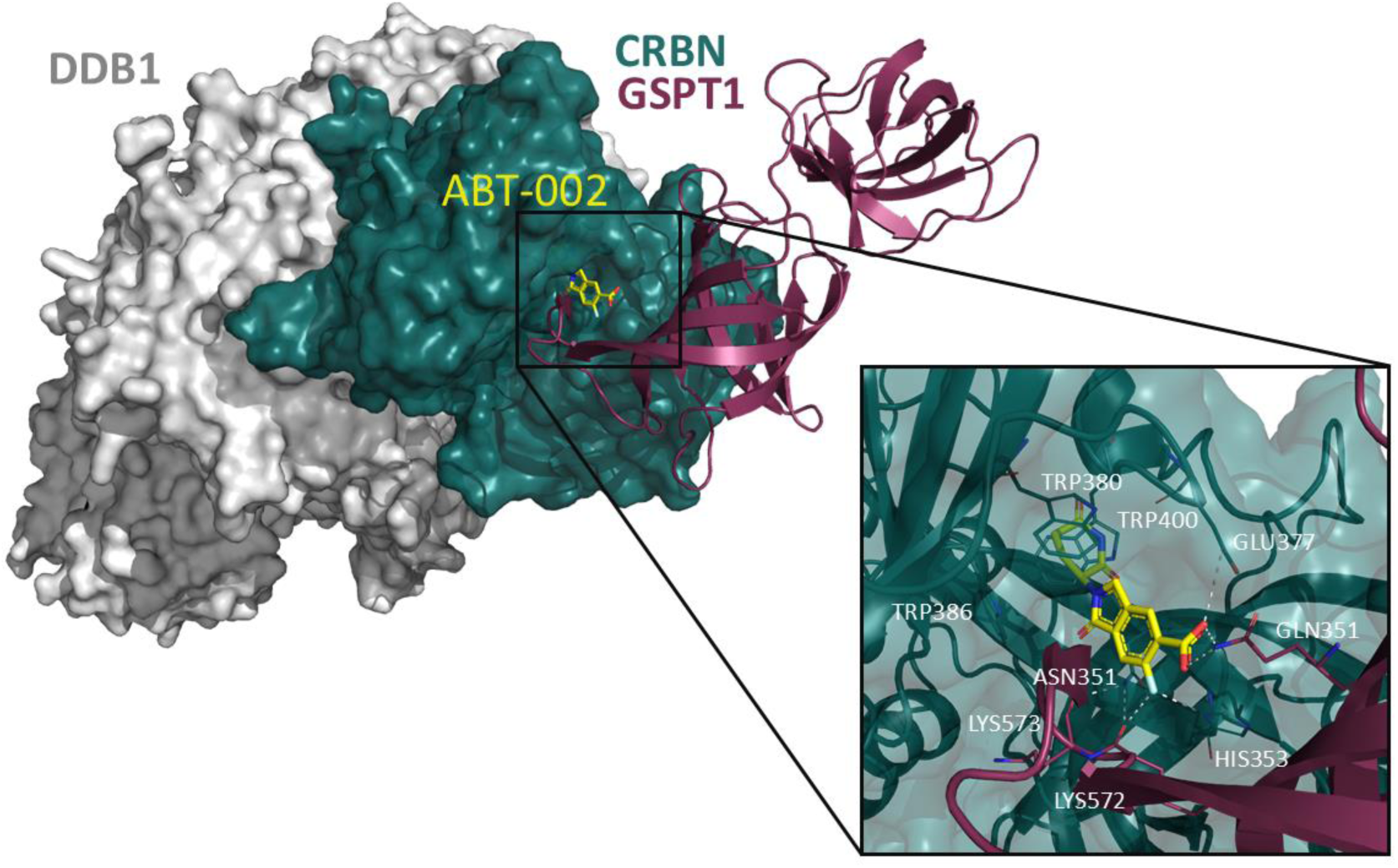
Crystal structure of the CRBN(40-442)-DDB1(FL)–GSPT1(300-496)–ABT-002 complex. Surface representation of CRBN (turquoise) and DDB1 (grey) and cartoon view of GSPT1 in violet. ABT-002 is presented as sticks (yellow). Close-up view shows CRBN/GSPT1 interface and ABT-002 binding mode in detail. Key amino acids are presented as sticks, hydrogen bonds are illustrated as white dashed lines.

### *In-vivo* effects of ABS-752

#### Pharmacokinetic (PK) studies in mice and NHPs

PK experiments showed metabolite formation (Fig. 7). The carboxylic acid ABT-002 was the main plasma compound, exceeding parent ABS-752 levels in mice and NHPs. VAP-1 presence/absence in plasma minimally impacted transformation, suggesting metabolism occurs mainly during intestinal absorption and in the liver (“first-pass” effect). Alcohol ABU-031 and aldehyde ABT-971 were detected (ABT-971 below LOQ in mice), along with acetylated amine ABT-003. Due to fast metabolism, ABS-752 shows high in vivo clearance in CD-1 mice, however ABT-002 exhibits moderate value of this parameter (Supplementary Table S6a). While drug (2.3±1.1h in mice, 2.2±0.4 in NHP) and metabolite ABT-002 (1.8±0.9h in mice, 4.9±2.2 in NHP) half-lives are moderate, PK/PD uncoupling is common for molecular glues and PROTACs, with effects persisting after drug clearance.^33,34^

**Figure 7.**
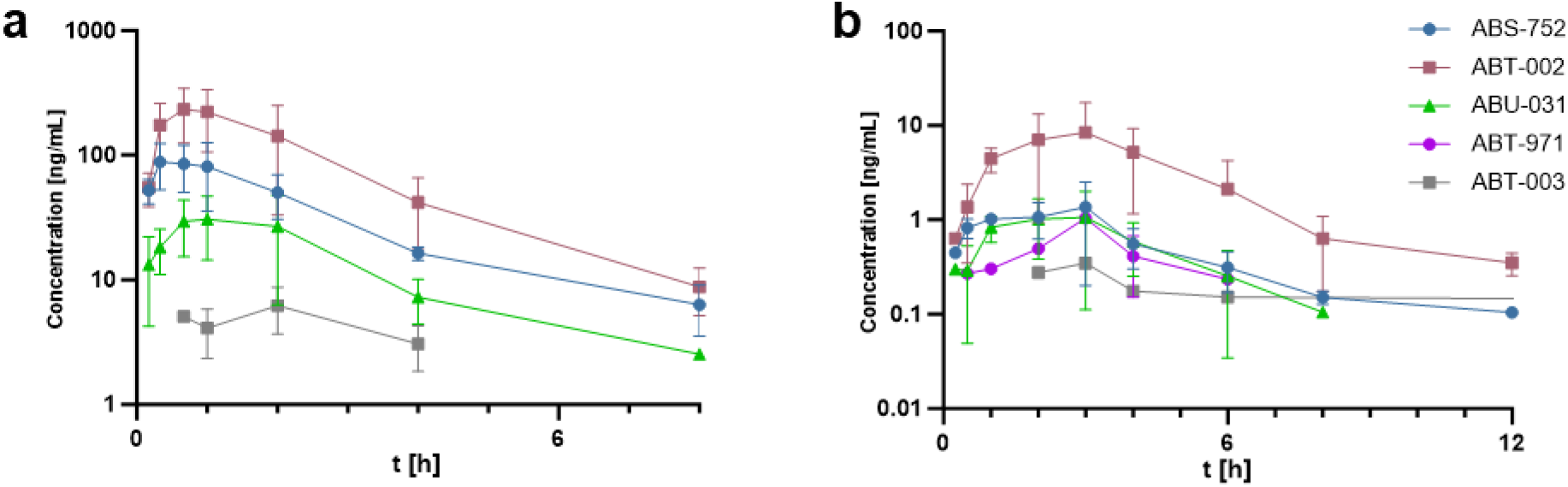
Pharmacokinetic profile of ABS-752 and known metabolites after oral administration of the ABS-752 in **(a)** male CD-1 mice (10 mg/kg, n = 3) and **(b)** male cynomolgus monkeys (0.3 mg/kg, n = 3).

#### Mouse Hep3B xenograft model

ABS-752’s *in vivo* efficacy was evaluated in HCC mouse models. Hep3B CDX xenograft (BID dosing: 1, 3, 10 mg/kg) showed strong tumor growth regression (in some animals complete eradication) at dose 10 mg/kg; 3 and 1 mg/kg induced remarkable tumor growth inhibition, demonstrating exceptional antitumor efficacy (Fig. 8a).

**Figure 8.**
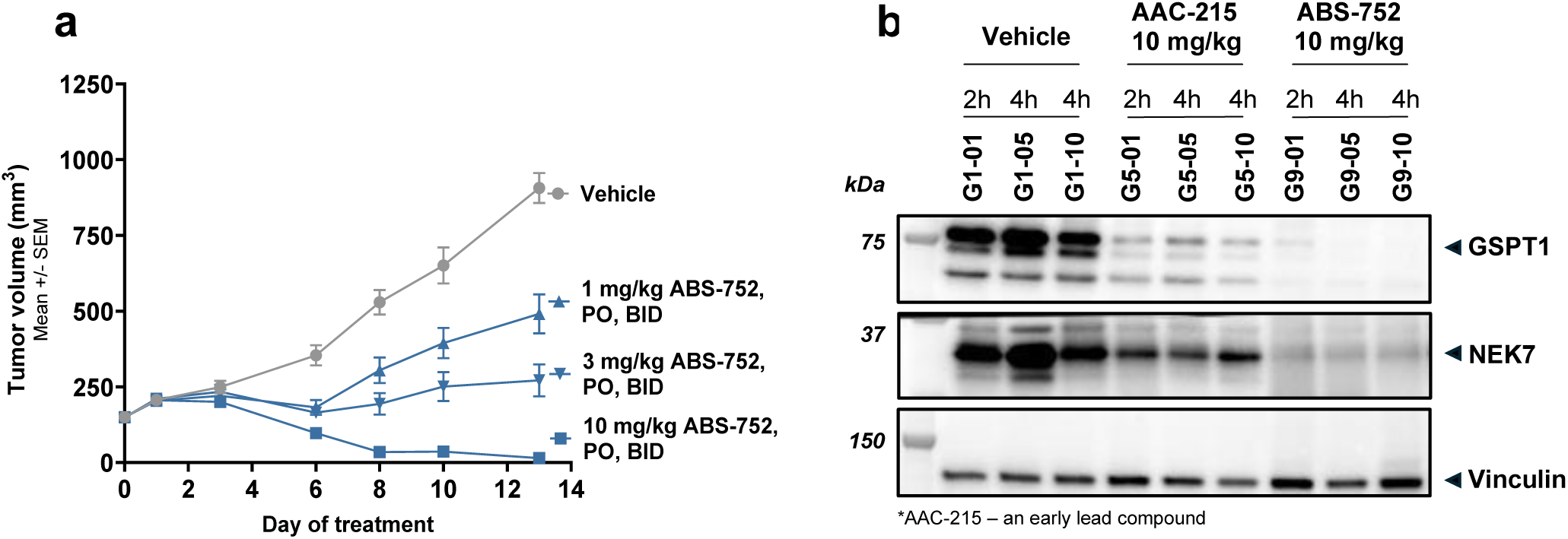
In Vivo Efficacy Study in Hep3B2.1-7 human hepatocellular carcinoma xenograft model. **(a)** Tumor growth kinetics data in the subcutaneous Hep3B2.1-7 human hepatocellular carcinoma xenograft model in female NSG mice. **(b)** Western blot results for GSPT1 and NEK7 degradation in the tumor samples obtained from animals treated with either AAC-215 or ABS-752, representative of the respective treatment groups.

Tumor excision (2, 4h post-dose) and GSPT1 Western blot analysis confirmed strong PD activity of ABS-752 (Fig. 8b).

#### Patient-derived xenograft tumor models in mice

Patient-derived xenografts (PDXs) are valuable for assessing early-stage cancer drugs. PDXs, generated by serially passaging human tumors in mice, retain human tumor heteroclonality and complex microenvironment, increasing clinical translational value.^35^ For HCC, PDX heteroclonality is especially valuable, as cell lines represent only ∼50% of HCC subtypes.^36^

ABS-752’s efficacy was evaluated in 10 randomly selected HCC PDX models. Tumor fragments were implanted subcutaneously (n=3/group), and therapy (100 mg/kg ABS-752 BID, PO) began at 100-200 mm³ tumor size, lasting 20-21 days. ABS-752 inhibited tumor growth in 8/10 models (16-100% TGI), with four models showing >50% TGI (Fig. 9a).

**Figure 9.**
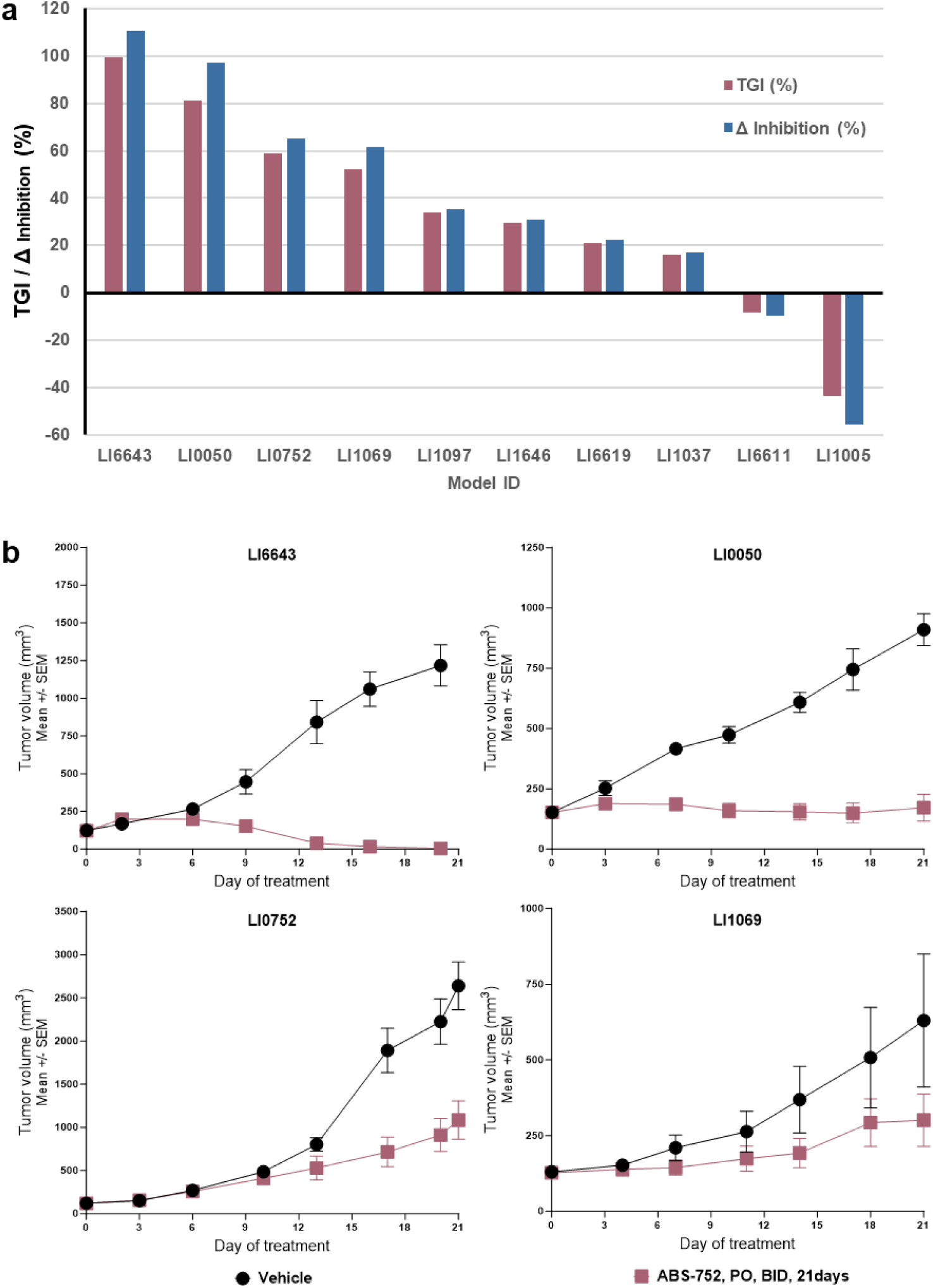
In Vivo Efficacy Study in human hepatocellular carcinoma PDX models. **(a)** Summary of ABS-752-Induced Tumor Growth Inhibition (%TGI and %Δ Inhibition at study end) for 10 randomly selected PDX models of human hepatocellular carcinoma in BALB/c nude mice. **(b)** Tumor growth kinetics data for the most sensitive models.

Tumor growth kinetics (Fig. 9b) showed complete tumor regression in the LI6643 model by day 15-16. Strong tumor growth control was observed in three other models (LI0050, LI0752, LI1069). Limited or no inhibition was seen in the remaining models, highlighting HCC heterogeneity.

## Discussion

In this study, we describe the discovery and optimization of ABS-752, a first-in-class molecular glue degrader targeting both GSPT1 and NEK7, with potential applications in the treatment of hepatocellular carcinoma (HCC). GSPT1 has been a known target of CRBN-based molecular glues since 2016^32^ and its degradation has been associated with strong cytotoxic activity in various cancer cell lines.^37–39^ However, developing GSPT1 degradation-based therapies has been challenging, likely due to a narrow therapeutic window and reported adverse events in humans.^40,41^

While several molecular glues have been reported to specifically degrade NEK7, there are no publicly disclosed molecular glues that have been shown to simultaneously co-degrade both NEK7 and GSPT1.^42–44^ NEK7 kinase is a major activator of the NLRP3 inflammasome, which leads to the production of IL-1β, an established factor involved in tumorigenic activity in various cancers.^45–47^ NLRP3 inflammasome activation by NEK7 relies on the scaffolding^48^, rather than the catalytic activity of NEK7, making it difficult to develop traditional small molecule inhibitors.

A key contribution of this work is the optimization of a prodrug activated by the VAP-1 enzyme. The clinical candidate, ABS-752, is deaminated by VAP-1 to the aldehyde, ABT-971. ABT-971, due to its good permeability across the cell membrane, should then be rapidly delivered to cells, where it would be oxidized to the carboxylic acid-containing molecule, ABT-002. The parent compound, ABS-752 and the intermediate, ABT-971 do not form ternary complex with CRBN and GSPT1 in biophysical assays, while ABT-002 does, which together with the high resolution X-ray structure confirm the active compound.

While the two-step activation mechanism by VAP-1 has been reported for emixustat^49^, its optimization and potential application have not been previously delineated. Using a battery of assays and inhibitors, we demonstrated this two-step metabolic conversion and characterized the differential cell membrane permeability of these molecules. Notably, the aldehyde ABT-971 exhibits excellent cell membrane permeability, whereas the active compound, ABT-002, is significantly less permeable (Fig. 4f), likely due to the high polarity imparted by its carboxylic acid group. This difference in cell penetration offers a potential therapeutic advantage for ABS-752. We anticipate that the rapid metabolic conversion of ABS-752 (Fig. 7) and the limited cell permeability of ABT-002 will result in higher local concentrations of ABT-002 in liver cancer cells. Given the observed short half-life of ABT-002, it will likely be rapidly eliminated from systemic circulation upon release from dying HCC cells (Supplementary Tables S6a-c). Importantly, VAP-1 expression is significantly elevated in inflamed livers, a common characteristic of HCC.^50^ We propose that these combined factors could contribute to a wider therapeutic window for ABS-752 compared to direct, systemically delivered GSPT1 degraders.

Serum VAP-1 is often upregulated in cancers, contributing to tumor progression.^51^ However, VAP-1 lower expression in tissue or serum has been observed in colorectal and thyroid cancers.^52,53^ These findings suggest that reduced VAP-1 might contribute to tumor resistance. Nevertheless, oral ABS-752 prodrug could potentially be activated to a membrane-permeable intermediate in the well-vascularized small intestine, portal vein, and non-transformed liver (high endothelial VAP-1) before reaching the tumor.^54,55^

We demonstrated that administering ABS-752 to cancer cell line cultures leads to potent and independent degradation of both GSPT1 and NEK7 (Fig. 1d-e). Further experiments using a GSPT1 G575N mutant Hep3B cell line, where GSPT1 is unable to bind to CRBN (Fig. 2), revealed that the primary cytotoxic effect of ABS-752 is driven by GSPT1 degradation.

ABS-752 exhibited remarkable activity in an aggressive Hep3B cell line-derived xenograft model. At a dose of 10 mg/kg twice daily, ABS-752 completely regressed tumors in this model. Finally, in a study using ten patient-derived xenograft (PDX) models of HCC, ABS-752 induced tumor growth inhibition greater than 50% in four cases. In four additional cases, we observed tumor inhibition ranging from 16% to 33% (Fig. 9). In an *in vivo* PKPD study in non-human primates (NHPs), we showed that 0.1 mg/kg of ABS-752 in 6h timepoint degraded ∼80% and ∼65% of GSPT1 and NEK7, respectively, (Fig. S9). Importantly, in a 28-day long toxicology study in NHPs no severe adverse events were observed (data not shown). These findings are complemented by viability study of primary human hepatocytes, where ABS-752 showed no cytotoxicity (Supplementary Fig. S3).

We speculate that the co-degradation of NEK7 will lead to the shutdown of the NLRP3 inflammasome, resulting in a significant reduction of IL-1β in the tumor microenvironment. IL-1β is a critical mediator of inflammation, fibrosis, and tumorigenesis in HCC^56–58^, contributing to an immunosuppressive tumor microenvironment, angiogenesis, and resistance to immunotherapy. Consequently, simultaneous degradation of both GSPT1 and NEK7 would offer attractive therapeutic benefits, potentially by synergistically reducing tumor growth and modulating the tumor microenvironment, which could open opportunities for combinatorial treatments.

In summary, ABS-752 is a compound in Phase I clinical trials, which selectively and preferentially degrades GSPT1 and NEK7 among other neosubstrates. Its mechanism of action depends on the activity of VAP-1, which is highly overexpressed in inflamed livers, including those of HCC patients. ABS-752 demonstrates compelling efficacy and safety *in vitro* and *in vivo*. These preclinical findings, along with favorable biopharmaceutical properties related to cell penetration and clearance of both the prodrug and the active metabolite, and the VAP-1 dependency, suggest that ABS-752 may offer a unique therapeutic opportunity for patients with liver cancer.

## Methods

### Cell culture

The cell lines used in the experiments are listed in the Supplementary Table S7. Cells were cultured at 37°C in a 5% CO_2_ humidified incubator and passaged every 2–3 days. All cell lines were regularly tested for mycoplasma contamination.

### Cellular CRBN NanoBRET Engagement Assay

NanoLuc-CRBN HEK293 cells were cultured and plated. Reference/test degraders in DMSO were arrayed in a 384-well plate and serially diluted. Cell suspensions were prepared with Opti-MEM and NanoBRET Tracer. For 100% inhibition, DMSO replaced Tracer. Cell-Tracer mixture was added to compound dilutions. After incubation, NanoBRET substrate was added. Donor/acceptor emissions were measured, and NanoBRET ratios were calculated and normalized to determine % inhibition.

The lytic assay, assessing target engagement with digitonin, followed the same procedure with modified cell suspension (Tracer and digitonin) and substrate (no inhibitor). % inhibition was calculated similarly.

### Degradation assays and Western Blot

In all experiments with adherent cell lines, media was changed prior to compound treatment. Stock solutions of compounds were prepared in DMSO (Miltenyi Biotec 170-076-303). NCI-H929 cells were plated in 6 or 12-well plates at a density of 2×10^5^ cells per mL and treated with the compounds directly after seeding. Hep 3B2.1-7 (Hep3B) cells were plated in 6-well plate or 60×15 Petri dishes at density 1.5×10^4^ - 2.1×10^4^ cells per cm^2^ depending on experimental setup and treated with the compounds 24 hours later. Kelly cells were plated in 6-well plate at 4.5×10^5^ cells per well and treated with the compounds the next day.

Protein extracts were prepared in 50 mM Tris HCl, pH 7.4,150 mM NaCl,1% NP-40, 0.25% sodium deoxycholate, 0.1% SDS and 1 mM EDTA buffer supplemented with protease (Roche 05056489001) and phosphatase inhibitors (Thermo Scientific 78426). The samples were centrifuged at 18 000 - 19 000 x g for 15 min at 4°C and the supernatants were collected. Protein concentrations in the lysates were determined using Pierce BCA kit (Thermo Scientific 23225) according to the manufacturer’s instructions. The proteins were resolved by SDS-PAGE, transferred onto nitrocellulose membranes and probed with a primary antibody, followed by a horseradish peroxidase (HRP)-conjugated secondary antibody (Supplementary Table S8).

Protein detection was performed using SuperSignal West Pico PLUS Chemiluminescent Substrate (Thermo Scientific 34578) or Westar Supernova (Cyanagen XLS3,0100) and the ChemiDoc MP Imaging System (BIO-RAD). Relative band intensities were quantified using Image Lab (BIO-RAD). The protein of interest (POI) signal values was normalized to the ß-actin loading control and relative POI levels were calculated as a percentage (%) of the DMSO control. The percentage downregulation/degradation values were calculated as 100% - % relative POI levels.

The Degradation assays and Western Blot for the VAP-1 dependent activity of ABS-752 in cellular experiments used in this work is described in the Supplementary Materials.

### CTG viability assay

Cell Titer Glo viability assay (Promega, cat. G7571, G7573) was performed in a 384-plate well format. The number of the following cells/well: Hep 3B2.1-7, JHH-7, HuH-7, HuH-1, SNU-398, SNU-886, Kelly, SNU-423, SNU-182, NCI-H929, was optimised for testing at 384 plate-well format. Cells were seeded at the indicated cell number (see Supplementary Table 9) at 384-well plate, white, solid bottom, with lid, tissue culture-treated, sterile (Greiner, cat. 781080) in a volume of 50 µl/well of a culture media (see section: cell culture) on the day before the compound treatment (adherent cells) and incubated O/N in a 5% CO_2_ incubator at 37°C or immediately before compound addition (suspension cells).

Before the compound addition the plates were spun down for 10 sec at 100 rcf. Each compound was tested in a serial dilution of 12 concentrations with 50 µM as the highest concentration in a technical duplicate. All the compounds were dissolved in a 100% DMSO.

Compound treatment was performed using Echo 555 Liquid Handler with customized protocols. As negative controls appropriate culture media (no cells, n=12) and DMSO-treated cells (n=22) were used. The final concentration of DMSO in a well was 0.25%. After the compound addition plates were spun down for 10 sec at 100 rcf. Following treatment, the cells were incubated for 72 h in a 5% CO_2_ incubator at 37°C.

CTG reagent was prepared according to the manufacturer’s protocol. Before the assay was carried out, plates were equilibrated to room temperature for 15 min. 12.5 µl of the CTG reagent was dispensed to each tested well. Subsequently, plates were shaken for 4 min at 460 rpm on PST-60HL-4 Plate Shaker-Thermostat at RT. Following shaking, plates were then left for 8 min in the dark w/o shaking to stabilize the signal. The luminescence was detected with the CLARIOstar Multimode Plate Reader (BMG Labtech), using optical Fireflymode. The Focus and Gain were adjusted on a selected well of one of negative ctrl (DMSO treated) wells. Target value was set as 80%.

Relative light unit (RLU) data were normalized and analyzed with a customized protocol. Normalization was based on the average of RLU of the negative and positive controls to calculate % inhibition. Percentage viability was calculated by the subtraction of percentage inhibition from 100%. Dose response was assessed by non-linear regression and IC_50_ calculation with the average of technical duplicate of percentage inhibition. The CTG viability procedure for the VAP-1 dependent activity of ABS-752 in cellular experiments used in this work is described in the Supplementary Materials.

Under contract with Charles River Discovery Research Services Germany GmbH (SOW P872A3), standardized screening of compounds was evaluated in 17 liver cancer cell lines (Supplementary Table S10) using a CTG based viability assay. Compounds were tested at 10 concentrations at half-logarithmic intervals (highest concentration 30 µM) under duplicate well conditions as a single series. The addition of the compound was carried out one day after seeding the cells. The incubation period with the applied compounds was set at 72 hours during a 96-hour incubation.

### TMT-based proteomics

Hep3B cells were treated with DMSO or ABS-752 for 6 hours. After washing with DPBS and detaching with Accutase, cells were washed (medium, ice-cold DPBS), and pellets were frozen for proteomic analysis (MSBioworks). Lysis was in urea buffer (QSonica). Lysate was incubated, mixed, centrifuged, and transferred to LoBind tubes. After protein quantitation (Qubit), 50µg of each lysate was reduced (DTT), alkylated (iodoacetamide), and digested (trypsin). The digest was adjusted (ammonium bicarbonate), cooled, and terminated (formic acid), then desalted (Waters Oasis HLB). Samples were lyophilized, frozen, reconstituted (HEPES/acetonitrile), and labeled (TMT-10plex). The reaction was quenched (hydroxylamine). Equal amounts of labeled peptide were combined, desalted (Waters HLB 6CC), lyophilized, and frozen. Pooled samples were fractionated (Agilent 1100 HPLC, Waters XBridge column). Samples were analyzed by nano LC-MS/MS (Waters NanoAcquity, Thermo Fisher Fusion Lumos). Peptides were loaded (trapping column) and eluted (75 µm analytical column, Luna C18 resin). The mass spectrometer used a data-dependent MS2 method (Orbitrap: 120,000/50,000 FWHM for MS/MS). Data were processed with MaxQuant (Andromeda).

### Protein purification

The purification procedure for the proteins used in this work is described in the Supplementary Materials and Methods section.

### Crystallization

CRBN(40-442)-DDB1(FL)–GSPT1(300-496)–ABT-002 complex was obtained by mixing CRBN(40-442)-DDB1(FL) (4.9 mg/ml), GSPT1(300-496) (10 mg/ml) and ABT-002 at a 1 : 1.3 : 3 molar ratio and 1 h incubation at room temperature. Crystallization trials were conducted at 19°C applying hanging-drop vapor diffusion method by mixing the complex with an equal volume of reservoir solution. Plate-shaped crystals were formed after 2 days in the 0.2 M Sodium citrate, 0.1 M Bis-Tris propane pH 7.5, 20% w/v PEG 3350 solution and flash frozen with the addition of 20% (v/v) ethylene glycol.

### Structural determination

Data for the CRBN-DDB1–GSPT1–ABT-002 complex were collected at the Deutsches Elektronen-Synchrotron DESY (Hamburg, Germany)^59,60^ to a resolution of 3.9 Å and processed automatically utilizing XDS package at the beamline.^61^ Initial phases were obtained by molecular replacement with PHASER^62^ using CRBN-DDB1–GSPT1–CC-885 structure (PDB ID: 5HXB^32^) as a search model. Two copies of CRBN-DDB1–GSPT1– ABT-002 complex were found in the asymmetric unit. The model was built using COOT^63^ and refined using REFMAC5^64^. Ligand topology was generated with AceDRG^65^. All figure models were generated using PyMOL (Schrödinger).

### Biochemical proximity assays

AlphaLISA ternary complex formation assays were performed for each target with CRBN/DDB1 under reduced light. Alpha beads were coated (neosubstrate/donor beads, CRBN/DDB1/acceptor beads). Bead mixes were dispensed into 384-well plates. Compounds (1 µM, 10 µM, triplicates) and DMSO controls were added (Echo 555). Plates were sealed, mixed, centrifuged, incubated, and read (EnSpire). Luminescence was normalized to reference compound (100%) (Supplementary Table S11).

TR-FRET ternary complex formation assays (dose response) used target protein (GSPT1/NEK7), CRBN/DDB1, Streptavidin-Eu cryptate, Anti-6His-d2, and DTT in PPI buffer. Protein mixture was dispensed into 384-well plates. Compounds (serial dilutions from 50 µM) were added (Echo 555, 0.5% DMSO final). Plates were sealed, mixed, centrifuged, and incubated. Emission (620/665 nm) was recorded (PHERAstar). Signal-to-background ratios were calculated and normalized. Data were analyzed (GraphPad Prism).

### Analysis of monoamine oxidases activity against ABS-752

Oxidation of ABS-752 by recombinant MAO-A (Sigma-Aldrich M7316-1VL), MAO-B (Sigma-Aldrich M7441-1VL) and mVAP-1 (R&D Systems 6107-AO-010) to the presumed metabolites of ABS-752 was analyzed using LC-MS/MS method. Enzyme in 1x assay buffer (5 µg/mL oxidase in 25 mM sodium phosphate buffer pH 7.4) was mixed with equal volume of 20 µM ABS-752 (20 mM stock in 100% DMSO) in 1x assay buffer. The following reaction mixtures were prepared: ABS-752 alone, ABS-752 and oxidase, ABS-752 with oxidase and a specific oxidase inhibitor PXS-4728A (Appendix_1). Samples with enzyme and enzyme supplemented with inhibitor were incubated for 30 minutes at room temperature (25°C) prior to adding substrate to the mixture (Time ‘0’). At Time ‘0’ the first sample aliquot was taken (50 µL). The samples were then transferred to a Thermoblock and incubated at 37°C. Additional samples were taken after 4, 8 and 20 hours. Reaction in the aliquot taken was immediately quenched by the addition of an equal volume of LC-MS grade methanol (ChemSolute 1203.2500). Samples were mixed, spun down, frozen in liquid nitrogen and stored at −80°C before being processed and analyzed using a LC-MS/MS method. Details are described in the Supplementary Materials and Methods section.

### Formation of ABS-752 metabolites in whole human tissue lysates

The frozen whole tissue lysates of heart aorta valve (Novus Biologicals NB820-59218) or cirrhotic liver (Novus Biologicals NB820-59328) were thawed on ice, then the suspension was mixed by pipetting. Initially, ABS-752 was dissolved in water at 100 mM. The series of 1.1x stock concentrations of ABS-752 were prepared by 2-fold serial dilutions (10 points; Supplementary Table S16) in PBS buffer (20 mM phosphate buffer pH 7.4, 2.7 mM KCl, 137 mM NaCl). Next 45 µL of each substrate dilution was transferred into a labelled 1.5 ml tube. Substrate 1.1x solutions were preincubated for 10 minutes at 37°C in the equilibrated water bath. After preheating, the sample was supplemented with 5 μL of the selected lysate stock (Time ‘0’) to give a final protein concentration from the lysate in the sample of 0.3 mg/mL. For each of the ABS-752 concentrations analyzed, 4 independent reaction mixtures were prepared, 2 without the inhibitor and 2 with the addition of 10 µM PXS-4728A (Appendix_2). The reaction was carried out for 60 minutes at 37°C and stopped by the addition of an equal volume of methanol (50 μL). Samples were mixed and immediately snap-frozen in liquid nitrogen. Solidified samples were stored at -80°C and shipped on dry ice for LC-MS/MS determination of substrate conversion products to Laboratory of Mass Spectrometry IBB PAN (Institute of Biochemistry and Biophysics, Polish Academy of Sciences in Warsaw, Poland). Details are described in the Supplementary Materials and Methods section.

### Amplex Red-based fluorogenic assay

hVAP-1 protein (Sigma-Aldrich SRP6241-10UG) was added to 2x assay buffer (25 mM sodium phosphate pH 7.4, 400 µM Amplex Red and 2 U/mL HRP enzyme (Thermo Fisher Scientific A12214)) at 50 µg/mL concentration (Oxidase master mix). 10 µL of Oxidase master mix was transferred onto a reaction plate (PerkinElmer 6008260) and 15 µL of 2x compound dilution in 1x reaction buffer was transferred to adjacent wells. The plate was sealed with polyester adhesive film and preheated at 37°C for 10 minutes. The seal was then removed and the plate inserted into a CLARIOstar plate reader chamber (BMG Labtech) adjusted at 37°C. After 60 s the plate was removed and 10 µL of 2x compound dilution was added into the wells containing Oxidase master mix using multichannel pipette and, following re-insertion, kinetic measurement was immediately started (filter settings: 545-20/600-40; gain: 600; number of cycles (total redout time): 150 (15 min)). ABS-752 was dissolved in water to create concentrations of: 250, 500, 750,1000, 3000 and 5000 µM. The final DMSO content was 1% (introduced with Amplex Red reagent). A well containing reaction buffer only was utilized as a control for hVAP-1 preparation effect on assay readout. To check compound effect on the Amplex Red non-enzymatic conversion, the same compound dilutions were mixed with Oxidase master mix without hVAP-1 enzyme and fluorescence intensity change was monitored. To determine the concentration of the product of the reaction catalyzed by hVAP-1 enzyme, a resorufin standard curve was prepared in 1x reaction buffer (0.0156-2 µM resorufin (Molecular Probes A12214)).

The initial velocity of ABS-752 oxidation was calculated based on linear regression slopes determined for each substrate concentration point (time 0-15 minute) and expressed as fluorescence intensity/min/well. The concentration of the formed reaction product was quantified using a resorufin standard curve with a linear regression slope. The velocity of ABS-752 oxidation was expressed as pmol product/min/µg of hVAP-1. Plots preparation, linear regression analysis and Michaelis-Menten constant (K_m_) determination were conducted using GraphPad Prism 10.3.1 for Windows.

### Pharmacokinetic studies

Non-GLP single dose PK studies were conducted in CD-1 mice (Aragen Life Sciences, Hyderabad/India) and cynomolgus monkeys (WuXi AppTec, Suzhou/China). The study protocols were approved by the local animal welfare and ethical review boards. The studies were carried out respecting all ethical regulations.

ABS-752 was administered as a solution in 0.5% (v/v) methylcellulose (400cP) and 0.2% (v/v) Tween 80.

Male CD-1 mice (*n* = 3) were treated with ABS-752 orally with dose 10 mg/kg. Blood samples were collected at 0.17, 0.33, 0.67, 1, 2, 4, 8 and 24 h after administration. LLOQ: 2 ng/mL (ABS-752, ABT-002, ABT-971, ABT-031, ABT-003).

Male cynomolgus monkeys (*n* = 3) were treated with ABS-752 orally with dose 0.3 mg/kg. Blood samples were collected at 0.17, 0.25, 0.5, 1, 2, 3, 4, 6, 8, 12 and 24 h. LLOQ: 0.1 ng/mL (ABS-752, ABT-031, ABT-003), 0.2 ng/mL (ABT-971, ABT-002).

Plasma was separated by centrifugation at 2500-3200 x g for 10-15 min and frozen at −80°C. Plasma samples were analyzed using liquid chromatography tandem mass spectrometry (LC-MS/MS). The pharmacokinetic parameters AUC_0-t_, C_max_, T_max_, and T_1/2_ were calculated using Phoenix WinNonlin 8.3.5.

### *In vivo* xenograft studies

The *in vivo* xenograft studies were conducted at Labcorp (Greenfield, US) - Hep3B model, and Crown Bioscience Inc. (Taicang, China) - PDX models.

Subcutaneous Hep3B2.1-7 human hepatocellular carcinoma xenograft model was conducted in female NSG mice (n=10/group). The treatment was initiated after randomization at overall mean tumor size approx. 155 mm^3^. ABS-752 was administrated per os (PO) twice a day (BID) for 14 days at 10, 3 & 1 mg/kg as a solution in 0.5% Methyl cellulose/0.2% tween-80. Following mice euthanasia on the last day of the treatment, tumors were excised and snap frozen in liquid nitrogen prior to WB analysis.

10 Subcutaneous patient-derived liver cancer xenograft (PDX) models (Crown Bioscience’s HuPrime® collection) were conducted in female BALB/c nude mice (n=3/group). The treatment was initiated when the mean tumor size reached approx. 100-200 mm^3^. ABS-752 was administrated per os (PO) twice a day (BID) for 20-21 days at 100 mg/kg as a solution in 0.5% Methyl cellulose/0.2% tween-80.

Tumor volumes were measured in two dimensions using a caliper, and the volume was expressed in mm^3^ using the formula: “V = (L x W x W)/2, where V was tumor volume, L was tumor length (the longest tumor dimension) and W was tumor width (the longest tumor dimension perpendicular to L).

Tumor Growth Inhibition (TGI) was calculated as TGI% = (1-T_i_/C_i_) × 100 where T_i_ and C_i_ are the mean tumor volumes of the treatment and control groups, respectively.

ΔInhibition was calculated as Mean % ΔInhibition = ((C_i_-C_0_) – (T_i_-T_0_)) / (C_i_-C_0_) * 100%, where Ti and Ci are the mean tumor volumes of the treatment and control groups, respectively, on a given day. T_0_ and C_0_ as the mean tumor volumes of the treatment and vehicle groups on Day 0.

## Acknowledgements

This research was co-financed by the European Regional Development Fund, European Funds for Smart Economy 2021-2027 (FENG), supported by the National Centre for Research and Development (NCBR, Poland) project grants no. POIR.01.01.01-00-0740/19; FENG.01.01-IP.01-1001/23.

The authors would like to express their gratitude to the research team at Captor Therapeutics including Piotr Biniarz and Agnieszka Gawurska for their collaborative efforts in collection and analysis of data. Their expertise has greatly assisted with the writing of this paper.

The authors gratefully acknowledge Google AI for providing access to Gemini 2.0 Flash, which was used to enhance the clarity and readability of the manuscript.

We acknowledge DESY (Hamburg, Germany), a member of the Helmholtz Association HGF, for the provision of experimental facilities. Parts of this research were carried out at PETRA III and we would like to thank Dr. Spyros D. Chatziefthymiou for assistance in using P11. Beamtime was allocated for proposal P-20010353.

## Author contributions

Conceptualization of the manuscript is performed by P.G., R.P., S.C., P.D., A.S-S., M.J.W. The original draft was written by P.G., R.P., S.C., A.S-S., M.J.W., and further revised/edited by D.C., P.D., T.D., M.Mi. Protein purification was conducted by D.G., K.J., K.L., M.Ma., K.P., J.W. under the supervision of A.Sz. Crystallization experiments and X-ray structure determination of GSPT1/ABT-002/CRBN-DDB1 complex were performed by M.K. Design and supervision of cell-based experiments were performed by K.E.O., P.D., J.M., M.J.W. with the assistance of A.Sn., M.Mi., S.Sa. Supervision of PK/PD and animal model experiments were directed by P.D. CTG assay and Western Blot were performed by J.L-G., A.Sn., J.M., M.Mi., P.R., M.J., A.A., A.Sa., O.M. NanoBRET assay was conducted by T.T. Generation of Hep3B G575N mutant cell line using the CRISPR/Cas9 system was performed by A.Sa. Supervision of the chemical synthesis and compound design were directed by S.C. Design of compounds and chemical synthesis were conducted by R.P., S.C., K.K. Supervision of project administration of pharmacokinetic studies, metabolic profiling with CYP enzymes, hepatocytes viability experiment were led by M.W. Design and performance of biochemical and biophysical experiments were conducted by P.G. Biochemical and cellular experiments on VAP-1-dependent ABS-752 activity were performed by M.Mi. Data analysis and interpretation were performed by P.G., R.P., K.E.O., M.K., M.W., S.C., P.D., T.D., J.M., M.Mi., A.Sa., A.S-S., M.J.W. Methods and supplementary information were written by P.G., R.P., M.K., M.W., J.L-G., M.Mi., P.R., A.Sa., A.S-S., A.Sz.

## Competing interests

Authors are or were employees and stockholders of Captor Therapeutics.

## Notes

### Summary of Updates

The author list for the preprint has been revised to include an experimenter who was not present in an earlier iteration of the manuscript.

