## Supplementary information for "Targeted degradation of GSPT1 and NEK7 by a molecular glue prodrug for treatment of HCC"

#### Supplementary Tables

***Table S1. Analysis of AAC-215 activity in CTG viability assay against HCC and H929 cell lines.*** *The absolute IC50 values presented are the geometric mean. Minimum viability is the arithmetic mean with standard deviation. Brackets include information on biological replications of the experiment. Abbreviations used: n.d. – not determined; absIC50 - absolute IC50; min viability – minimum viability; HCC – hepatocellular carcinoma; MM – multiple myeloma.*

| **Cell line** | **Lineage origin** | **absIC50 [nM]** | **Min viability [%]** |
|---|---|---|---|
| Hep3B | HCC | 73.3 ×/÷ 4.86 (n > 5) | 24.6 ± 10.1 (n > 5) |
| JHH-7 | HCC | 207 (n > 3) | 32.3 ± 4.0 (n > 3) |
| HuH-7 | HCC | 965 (n > 3) | 35.2 ± 5.4 (n > 3) |
| HuH-1 | HCC | n. d. | 61.7 ± 3.76 (n > 3) |
| SNU-398 | HCC | > 22800 (n > 3) | 30.7 ± 11.3 (n > 3) |
| SNU-182 | HCC | n.d. | 79.8 ± 7.57 (n > 3) |
| SNU-423 | HCC | n.d. | 92.3 ± 4.43 (n > 3) |
| SNU-886 | HCC | n.d. | 87.0 ± 5.39 (n > 3) |
| H929 | MM | n.d. | 98.4 ± 11.4 (n = 3) |

***Table S2. Results of AlphaLISA-based ternary complex formation mediated by molecular glues.*** *Data presented are means with standard deviation for the Target-CRBN ternary complex (TCF) level normalized to the reference compound. Values in parentheses indicate the number of biological repeats. The data were color coded according to the observed mean response: TCF < 25% (red), 25% ≤ TCF < 75% (orange), TCF ≥ 75% (green).*

| **Compound ID** | **Compound [µM]** | **GSPT1 (300-496 aa)** | **CK1α** | **NEK7** | **SALL4 (degron)** | **IKZF1 (degron)** |
|---|---|---|---|---|---|---|
| AAC-215 | 1 | 0.140 ± 0.162 (n=4) | 3.68 ± 7.01 (n=4) | 0.807 ± 1.40 (n=3) | 73.1 ± 3.73 (n=3) | 72.9 ± 6.98 (n=5) |
| | 10 | 0.0950 ± 0.112 (n=4) | 12.4 ± 6.21 (n=4) | 1.42 ± 2.46 (n=3) | 74.2 ± 3.50 (n=3) | 69.9 ± 9.88 (n=5) |
| ABR-522 | 1 | 93.6 ± 9.45 (n=4) | 16.1 ± 10.7 (n=5) | 176 ± 11.6 (n=4) | 92.8 ± 4.00 (n=3) | 2.11 ± 0.926 (n=3) |
| | 10 | 96.9 ± 10.2 (n=4) | 46.2 ± 16.3 (n=5) | 235 ± 24.7 (n=4) | 111 ± 3.80 (n=3) | 7.09 ± 1.83 (n=3) |
| ABR-321 | 1 | 1.42 ± 0.466 (n=4) | 5.89 ± 6.04 (n=5) | 0.197 ± 0.341 (n=3) | 91.8 ± 13.2 (n=3) | 21.4 ± 2.84 (n=3) |
| | 10 | 1.70 ± 0.336 (n=4) | 21.7 ± 11.4 (n=5) | 0.980 ± 1.61 (n=3) | 102 ± 8.94 (n=3) | 32.9 ± 2.29 (n=3) |
| ABT-394 | 1 | 11.2 ± 7.47 (n=4) | 9.17 ± 18.3 (n=4) | 0.413 ± 0.716 (n=3) | 80.5 ± 7.42 (n=3) | 11.5 ± 3.35 (n=4) |
| | 10 | 16.3 ± 10.2 (n=4) | 26.7 ± 7.10 (n=4) | 4.03 ± 2.51 (n=3) | 90.7 ± 6.49 (n=3) | 23.5 ± 13.6 (n=4) |
| ABR-958 | 1 | 19.9 ± 1.72 (n=3) | 13.2 ± 22.8 (n=5) | 1.26 ± 1.13 (n=3) | 103 ± 7.65 (n=3) | 37.5 ± 11.0 (n=4) |
| | 10 | 22.8 ± 2.08 (n=3) | 28.7 ± 13.4 (n=5) | 2.41 ± 3.56 (n=3) | 111 ± 7.35 (n=3) | 49.0 ± 15.5 (n=4) |

| | | | | | | |
|---|---|---|---|---|---|---|
| ABS-752 | 1 | 0.230 ± 0.503 (n=10) | 12.9 ± 10.4 (n=9) | 0.944 ± 0.904 (n=5) | 77.5 ± 7.53 (n=5) | 13.7 ± 9.42 (n=10) |
| | 10 | 0.252 ± 0.448 (n=10) | 14.2 ± 9.38 (n=9) | 1.08 ± 1.25 (n=5) | 79.6 ± 6.75 (n=5) | 22.9 ± 11.2 (n=10) |
| ABT-002 | 1 | 86.9 ± 11.7 (n=4) | 35.5 ± 17.3 (n=4) | 130 ± 24.4 (n=4) | 119 ± 8.70 (n=3) | 1.29 ± 1.47 (n=4) |
| | 10 | 93.6 ± 6.49 (n=4) | 53.0 ± 19.6 (n=4) | 162 ± 38.8 (n=4) | 132 ± 7.55 (n=3) | 1.65 ± 0.477 (n=4) |
| ABT-003 | 1 | 4.87 ± 1.76 (n=4) | 34.8 ± 20.0 (n=4) | 0.750 ± 1.13 (n=4) | 91.7 ± 5.45 (n=3) | 9.15 ± 5.17 (n=4) |
| | 10 | 4.41 ± 2.98 (n=4) | 41.0 ± 17.6 (n=4) | 0.640 ± 1.11 (n=4) | 95.8 ± 9.26 (n=3) | 16.5 ± 5.26 (n=4) |
| ABT-971 | 1 | 16.6 ± 7.77 (n=4) | 31.3 ± 11.7 (n=4) | 3.07 ± 5.60 (n=4) | 76.0 ± 12.3 (n=4) | 9.88 ± 3.48 (n=4) |
| | 10 | 20.1 ± 8.72 (n=4) | 30.0 ± 4.14 (n=4) | 0.975 ± 1.74 (n=4) | 79.3 ± 13.5 (n=4) | 16.3 ± 9.04 (n=4) |
| ABU-031 | 1 | 37.2 ± 4.68 (n=4) | 45.5 ± 20.9 (n=4) | 1.42 ± 1.91 (n=4) | 103 ± 6.61 (n=4) | 21.5 ± 7.24 (n=4) |
| | 10 | 40.5 ± 3.59 (n=4) | 51.3 ± 6.82 (n=4) | 3.95 ± 2.28 (n=4) | 106 ± 5.85 (n=4) | 35.1 ± 0.763 (n=4) |
| CC-90009 | 1 | 100 | 2.29 ± 4.58 (n=4) | 0.00 ± 0.00 (n=3) | 92.8 ± 4.41 (n=3) | 41.8 ± 4.39 (n=4) |
| | 10 | 100 | 5.20 ± 7.46 (n=4) | 0.543 ± 0.513 (n=3) | 90.3 ± 4.06 (n=3) | 46.3 ± 5.95 (n=4) |
| Reference compound | | CC-90009 (100%) | FPFT-2216 (100%) | ABS-674 (100%) | Lenalidomide (100%) | Lenalidomide (100%) |

***Table S3. CTG assay results for ABS-752 and its isolated metabolites in wild type Hep3B cell line.*** *The absolute IC50 values presented are geometric means. Minimum viability is the arithmetic mean with standard deviation. Brackets include information on biological replications of the experiment. Abbreviations used: n.d. – not determined; absIC50 - absolute IC50; min viability – minimum viability.*

| Compound ID | absIC50 [nM] | Min viability [%] |
|---|---|---|
| ABS-752 | 24.6 ×/÷ 2.25 (n > 5) | 30.2 ± 7.95 (n > 5) |
| ABT-002 | 20400×/÷ 1270 (n = 5) | 43.1 ± 3.28 (n = 5) |
| ABT-003 | n.d. | 95.4 ± 2.35 (n = 5) |
| ABT-971 | 103×/÷ 1.74 (n = 5) | 33.9 ± 3.92 (n = 5) |
| ABU-031 | n.d. | 89.4 ± 2.24 (n = 5) |

***Table S4. Activity of ABS-752 and its metabolites in the NanoBRET CRBN in-cell displacement assay.*** *Inhibition of intracellular Tracer/NanoLuc-CRBN complex formation by ABS-752 and its presumed metabolites. Determined geomean absolute IC50 (absIC50) with standard deviation. Values in parentheses indicate the number of biological repeats. For NanoBRET assay without added detergent, absIC50 was not determined (n.d.) for ABT-002 due to its activity not exceeding 50% inhibition.*

| Compound ID | NanoBRET absIC50 [M] | |
|---|---|---|
| | No permeabilization | Permeabilization |
| ABS-752 | 340E-09 ×/÷ 1.43 (n > 4) | 850E-09 ×/÷ 1.05 (n = 4) |
| ABT-002 | n.d. (n = 4) | 1.07E-06 ×/÷ 1.06 (n = 4) |
| ABT-003 | 570E-09 ×/÷ 1.14 (n = 4) | 634E-09 ×/÷ 1.01 (n = 2) |
| ABT-971 | 16.6E-09 ×/÷ 1.15 (n = 4) | 699E-09 ×/÷ 1.10 (n = 2) |
| ABU-031 | 191E-09 ×/÷ 1.14 (n = 2) | 685E-09 ×/÷ 1.11 (n = 2) |

***Table S5. Data collection and refinement statistics for CRBN(40-442)-DDB1(FL)–GSPT1(300-496)–ABT-002 complex.***

| | CRBN(40-442)-DDB1(FL)–GSPT1(300-496)–ABT-002 |
|---|---|
| **Data collection** | |
| Space group | *P* 1 2 1 |
| Cell dimensions | |
| a, b, c (Å) | 158.34, 112.38, 177.94 |
| α, β, γ (°) | 90.0, 95.25, 90.0 |
| Resolution (Å) | 49.49-3.9 (4.14-3.90) |
| $R_{merge}$ | 34.6 (161.4) |
| $I / \sigma I$ | 5.06 (1.11) |
| $CC_{1/2}$ | 97.8 (48.3) |
| Completeness (%) | 96.5 (96.8) |
| Redundancy | 4.44 (4.28) |
| **Refinement** | |
| Resolution (Å) | 49.49-3.90 |
| No. reflections | 244556 (37884) |
| $R_{work}$ / $R_{free}$ | 0.259 / 0.336 |
| No. atoms | |
| Protein | 25975 |
| Ligand/ion | 44/2 |
| Water | 0 |
| *B*-factors | |
| Protein | 135.73 |
| Ligand/ion | 91.55 |
| Water | - |
| R.m.s. deviations | |
| Bond lengths (Å) | 0.0038 |
| Bond angles (°) | 1.326 |
| **PDB ID** | **9HNE** |

Values in parentheses relate to highest-resolution shell.

#### Pharmacokinetic analyses of ABS-752 and its metabolites in CD-1 mice and cynomolgus macaques

Non-GLP single dose PK studies were conducted in CD-1 mice (Aragen Life Sciences, Hyderabad/India) and cynomolgus monkeys (WuXi AppTec, Suzhou/China). The study protocols were approved by the local animal welfare and ethical review boards. The studies were carried out respecting all ethical regulations.

Male CD-1 mice (n = 3) were treated with ABS-752 orally with dose 10 mg/kg in 0.5% (v/v) methylcellulose (400cP) and 0.2% (v/v) Tween 80. Blood samples were collected at 0.17, 0.33, 0.67, 1, 2, 4, 8 and 24 h after administration. LLOQ: 2 ng/mL (ABS-752, ABT-002, ABT-971, ABT-031, ABT-003).

Male CD-1 mice (n = 3, 4 groups) were treated with respectively: ABS-752, ABT-002, ABT-971 & ABU-031 intravenously with dose 1 mg/kg in 5% DMSO/5% Solutol HS-15:Ethanol (1:1 v/v)/ 90% normal saline. Blood samples were collected at 0.017, 0.083, 0.25, 0.5, 1, 2, 4, 8, and 24h. LLOQ: 1 ng/mL.

Male cynomolgus monkeys (n = 3) were treated with ABS-752 orally with dose 0.3 mg/kg in 0.5% (v/v) methylcellulose (400cP) and 0.2% (v/v) Tween 80. Blood samples were collected at 0.17, 0.25, 0.5, 1, 2, 3, 4, 6, 8, 12 and 24h. LLOQ: 0.1 ng/mL (ABS-752, ABT-031, ABT-003), 0.2 ng/mL (ABT-971, ABT-002).

Plasma was separated by centrifugation at 2500-3200 ×g for 10-15 min and frozen at −80 °C. Plasma samples were analyzed using liquid chromatography tandem mass spectrometry (LC-MS/MS). The pharmacokinetic parameters AUC0-t, $C_{max}$, $T_{max}$, and $T_{1/2}$ were calculated using Phoenix WinNonlin 8.3.5.

$V_d$ – volume of distribution; Cl – clearance; $C_0$ – extrapolated initial concentration; $T_{max}$ – time of maximum concentration; $T_{1/2}$ – elimination half-life; $AUC_{0\text{-}inf}$ – the integral of the concentration-time curve

***Table S6a. Pharmacokinetic data after dosing ABS-752, ABT-002, ABT-971 & ABU-031 (IV, 1 mg/kg) respectively in CD-1 mice.***

| Time (h) \ Concentration (ng/mL) | ABS-752 | ABT-002 | ABT-971 | ABU-031 |
|---|---|---|---|---|
| **0.017** | 1396 ± 165 | 3670 ± 460 | 17.2 ± 1.0 | 1790 ± 150 |
| **0.083** | 347 ± 111 | 1340 ± 700 | 7.1 ± 5.4 | 820 ± 70 |
| **0.25** | 56.5 ± 7.0 | 379 ± 228 | BLQ | 286 ± 14 |
| **0.5** | 33.2 ± 9.5 | 136 ± 115 | BLQ | 149 ± 50 |
| **1** | 22.2 ± 8.0 | 69 ± 68 | BLQ | 36.7 ± 23.9 |
| **2** | 6.5 ± 3.8 | 18.4 ± 25.3 | BLQ | 8.1 ± 4.9 |
| **4** | 2.8 ± ND | 28.4 ± 37.9 | BLQ | 1.7 ±0.2 |
| **8** | BLQ | BLQ | BLQ | BLQ |
| **24** | BLQ | BLQ | BLQ | BLQ |
| **$C_0$ (ng/mL)** | 2000 ± 190 | 4920 ± 960 | ND | 2190 ± 220 |
| **$V_d$ (L/kg)** | 6.5 ± 3.0 | 1.9 ± 1.5 | ND | 2.8 ± 1.2 |
| **Cl (mL/min/kg)** | 112 ± 8 | 36.2 ± 16.5 | ND | 52.9 ± 4.0 |
| **$AUC_{0\text{-}inf}$ (ng·h/mL)** | 149 ± 10 | 548 ± 300 | ND | 316 ± 25 |

*ND – not determined, BLQ – below limit of quantification*

*Table S6b. Pharmacokinetic data after dosing ABS-752 (PO, 10 mg/kg) in CD-1 mice.*

| Time (h) \ Concentration (ng/mL) | ABS-752 (parent) | ABT-002 | ABT-003 | ABT-971 | ABU-031 |
|---|---|---|---|---|---|
| **0.17** | 52.2 ± 11.6 | 55.2 ± 16.6 | BLQ | BLQ | 13.2 ± 9.0 |
| **0.33** | 88.6 ± 35.7 | 174.3 ± 86.2 | BLQ | BLQ | 18.2 ± 7.2 |
| **0.67** | 85.3 ± 35.0 | 234 ± 109 | 5.1 ± 0.3 | BLQ | 29.4 ± 14.0 |
| **1** | 81.0 ± 45.6 | 221 ± 115 | 4.1 ± 1.8 | BLQ | 30.7 ± 16.3 |
| **2** | 50.3 ± 19.6 | 142 ± 109 | 6.2 ± 2.5 | BLQ | 26.8 ± 20.6 |
| **4** | 16.3 ± 2.8 | 42.7 ± 24.2 | 3.1 ± 1.2 | BLQ | 2.5 ± ND |
| **8** | BLQ | BLQ | BLQ | BLQ | BLQ |
| **$C_{max}$ (ng/mL)** | 97.2 ± 42.7 | 242 ± 119 | 5.6 ± 2.7 | ND | 33.8 ± 17.6 |
| **$T_{max}$ (h)** | 0.8 ± 0.2 | 0.9 ± 0.2 | 1.3 ± 0.6 | ND | 1.2 ± 0.7 |
| **$T_{1/2}$ (h)** | 2.3 ± 1.1 | 1.8 ± 0.9 | ND | ND | ND |
| **$AUC_{0\text{-}inf}$ (ng·h/mL)** | 262.5 ± 42.1 | 617 ± 298 | 12.3 ± 9.8 | ND | 85.1 ± 39.4 |

*ND – not determined, BLQ – below limit of quantification*

*Table S6c. Pharmacokinetic data after dosing ABS-752 (PO, 0.3 mg/kg) in cynomolgus macaque.*

| Time (h) \ Concentration (ng/mL) | ABS-752 (parent) | ABT-002 | ABT-003 | ABT-971 | ABU-031 |
|---|---|---|---|---|---|
| **0.117** | BLQ | BLQ | BLQ | BLQ | BLQ |
| **0.25** | 0.45 ± ND | 0.63 ± ND | BLQ | BLQ | 0.30 ± ND |
| **0.5** | 0.82 ± 0.19 | 1.4 ± 1.0 | BLQ | 0.27 ± ND | 0.29 ± 0.24 |
| **1** | 1.02 ± 0.05 | 4.5 ± 1.3 | BLQ | 0.30 ± ND | 0.83 ± 0.26 |
| **2** | 1.07 ± 0.45 | 7.1 ± 6.1 | 0.28 ± ND | 0.49 ± ND | 1.02 ± 0.64 |
| **3** | 1.4 ± 1.2 | 8.4 ± 9.1 | 0.35 ± ND | 1.0 ± ND | 1.06 ± 0.94 |
| **4** | 0.55 ± 0.25 | 5.2 ± 4.0 | 0.18 ± ND | 0.41 ± 0.26 | 0.59 ± 0.34 |
| **6** | 0.31 ± 0.15 | 2.1 ± 2.1 | 0.15 ± ND | 0.24 ± ND | 0.25 ± 0.22 |
| **8** | 0.15 ± 0.02 | 0.63 ± 0.45 | BLQ | BLQ | 0.10 ± ND |
| **12** | BLQ | 0.35 ± 0.10 | BLQ | BLQ | BLQ |
| **24** | BLQ | 0.34 ± 0.16 | BLQ | BLQ | BLQ |
| **$C_{max}$ (ng/mL)** | 1.57 ± 0.96 | 9.3 ± 8.3 | 0.35 ± ND | 1.0 ± ND | 1.3 ± 0.8 |
| **$T_{max}$ (h)** | 2.0 ± 1.0 | 2.3 ± 1.1 | 3.0 ± ND | 3.0 ± ND | 2.3 ± 1.1 |
| **$T_{1/2}$ (h)** | 2.2 ± 0.4 | 4.9 ± 2.2 | ND | 1.4 ± ND | 1.4 ± 0.1 |
| **$AUC_{0\text{-}inf}$ (ng·h/mL)** | 5.5 ± 2.2 | 38.0 ± 29.2 | 3.8 ± ND | 3.4 ± ND | 4.3 ± 2.4 |

*ND – not determined, BLQ – below limit of quantification*

### Supplementary Figures

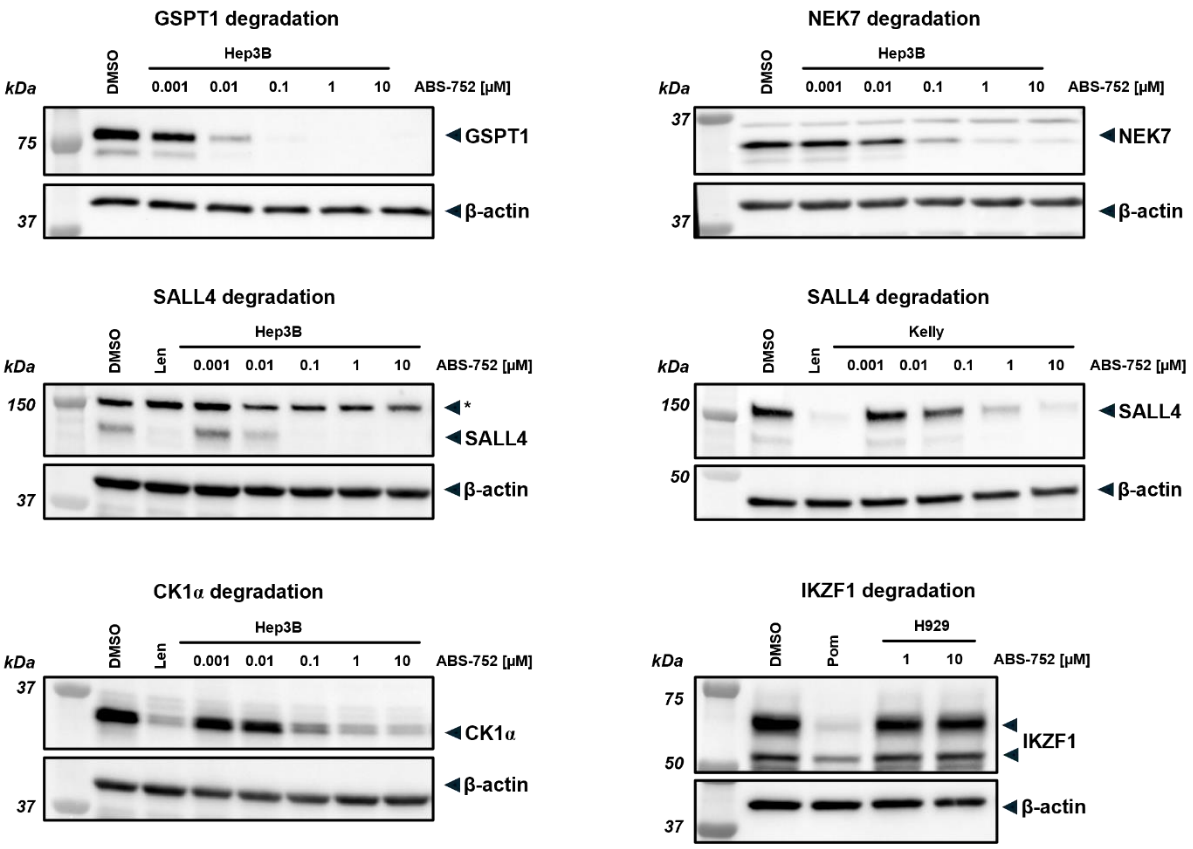

***Figure S1. Representative Western blot results for GSPT1, NEK7, SALL4, CK1α and IKZF1 degradation after 24-hour treatment with various concentrations of ABS-752 in Hep3B, Kelly or H929 cell lines.** The molecular weight of corresponding standards is indicated on the left.*

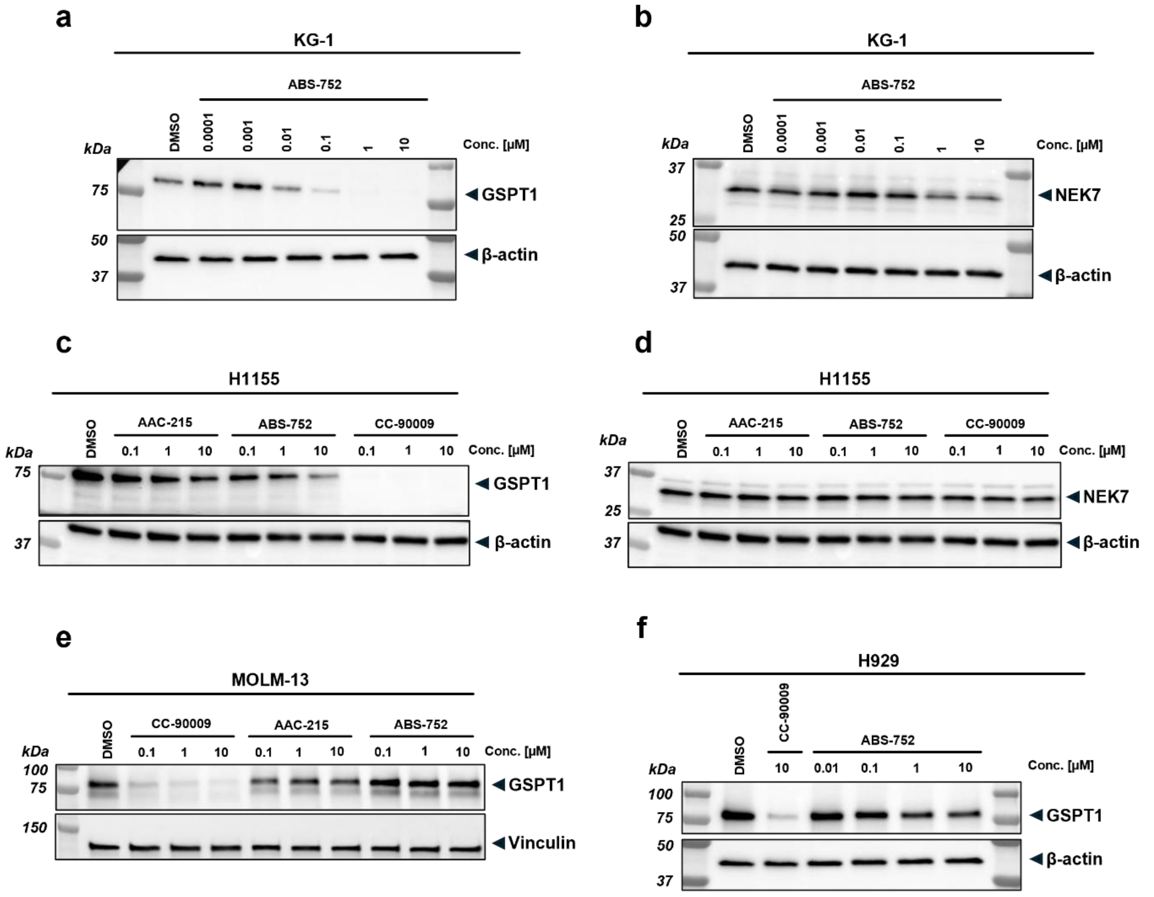

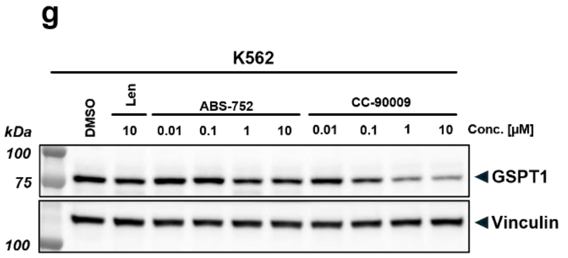

**h**

| Cell line | Lineage origin | absIC50 [nM] | Min viability [%] |
|---|---|---|---|
| H929 | mutiple myeloma, MM | n. d. | 84.0 ± 5.14 |
| KG-1 | leukemia, AML | 21.6 ×/÷ 1.70 | 2.70 ± 1.52 |
| Kasumi-1 | leukemia, AML | 109 ×/÷ 1.59 | 6.24 ± 1.51 |
| Kelly | neuroblastoma | 52.5 ×/÷ 1.53 | 2.76 ± 1.19 |
| MOLM-13 | leukemia, AML | n. d. | 94.2 ± 7.61 |
| MDS-L | human myelodysplatic syndrome | 832 ×/÷ 1.16 | 42.6 ± 5.46 |

***Figure S2. Activity of ABS-752 in selected non-HCC cell lines.*** ***(a) - (g)*** *Representative Western blot results for GSPT1 and NEK7 degradation after 24-hour treatment with various concentrations of ABS-752 in KG-1, H1155, MOLM-13, H929 and K562 cell lines. The molecular weight of corresponding standards is indicated on the left.* ***(h)*** *Analysis of ABS-752 activity in CTG viability assay against selected non-HCC cell lines. The absolute IC50 values presented are the geometric mean. Minimum viability is the arithmetic mean with standard deviation. The results are of at least three independent replicates. Abbreviations used: n.d. – not determined; absIC50 - absolute IC50; min viability – minimum viability.*

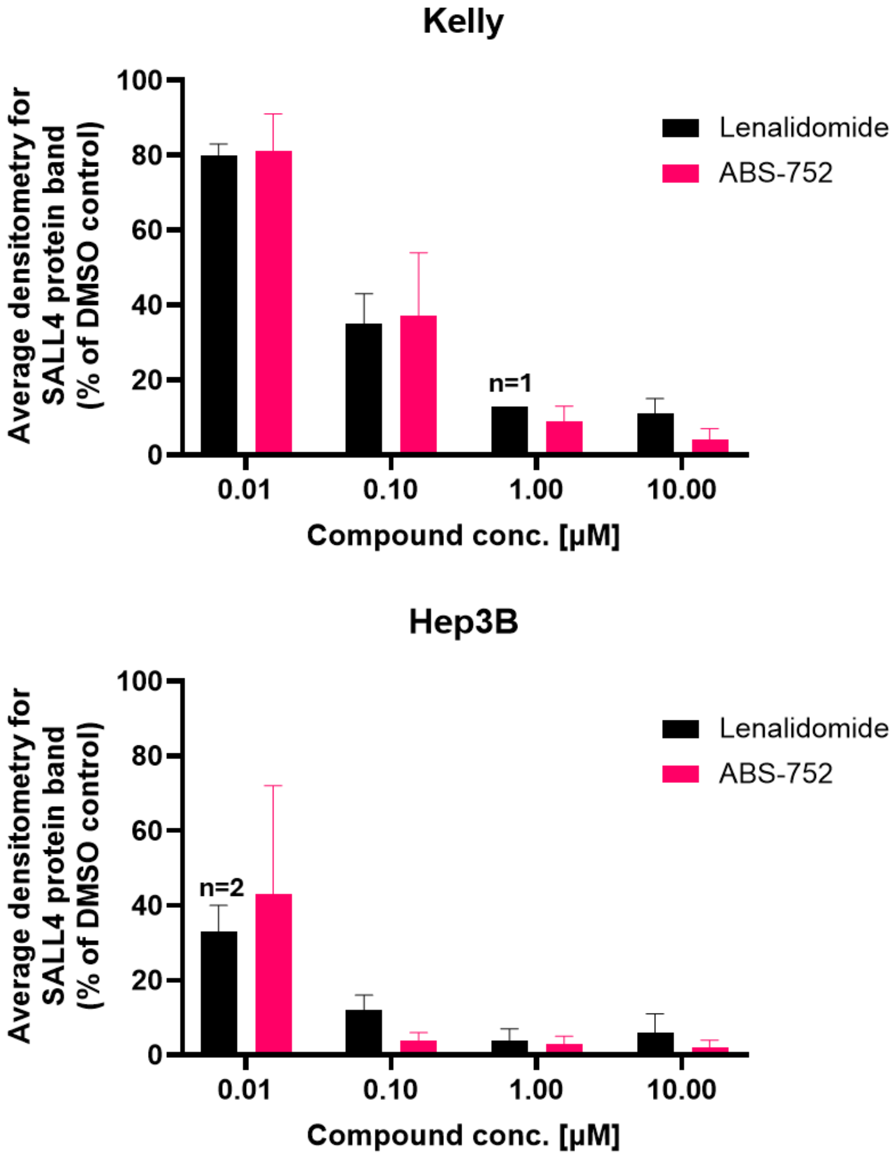

***Figure S3*. *Lenalidomide causes a comparable reduction in SALL4 protein level in the Kelly and Hep3B cell lines compared to compound ABS-752.*** *Western blot densitometry results for SALL4 protein degradation in Kelly and Hep3B cell lines after 24-hour treatment with lenalidomide and ABS-752. Data points are means with standard deviation from at least 3 independent experiments (For exceptions, the number of repetitions is given above the bar).*

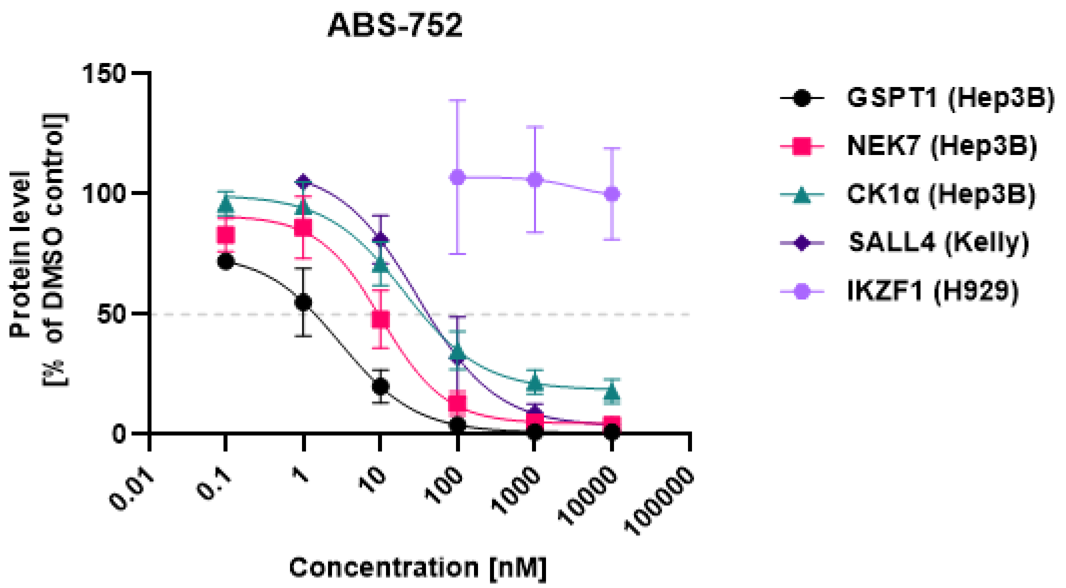

***Figure S4. Effect of ABS-752 on GSPT1, NEK7, CK1α and SALL4 protein levels after 24 hours of treatment based on Western blot.*** *Presented data are means with standard deviation which shown were calculated based on at least two biological replicates. Names in parentheses next to the target name indicate the cell line in which protein degradation was tested.*

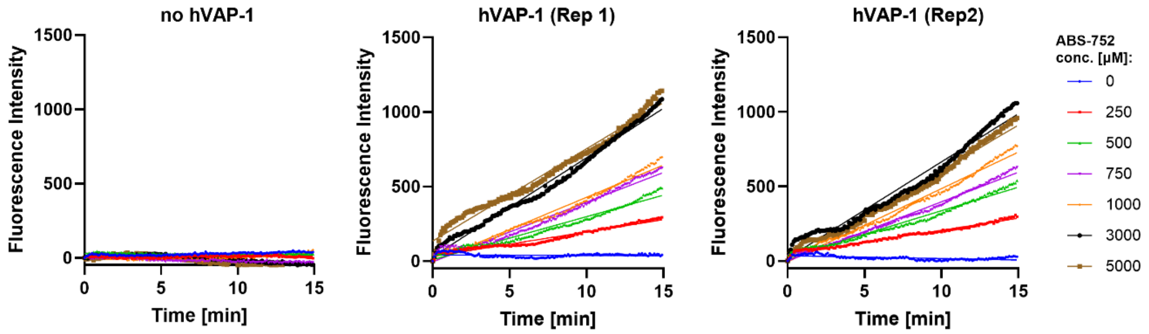

***Figure S5. Rate of change in fluorescence intensity during incubation of ABS-752 without and with recombinant hVAP-1 enzyme****. Results were obtained using Amplex Red monoamine oxidase kit (Thermo Fischer Scientific A12214).*

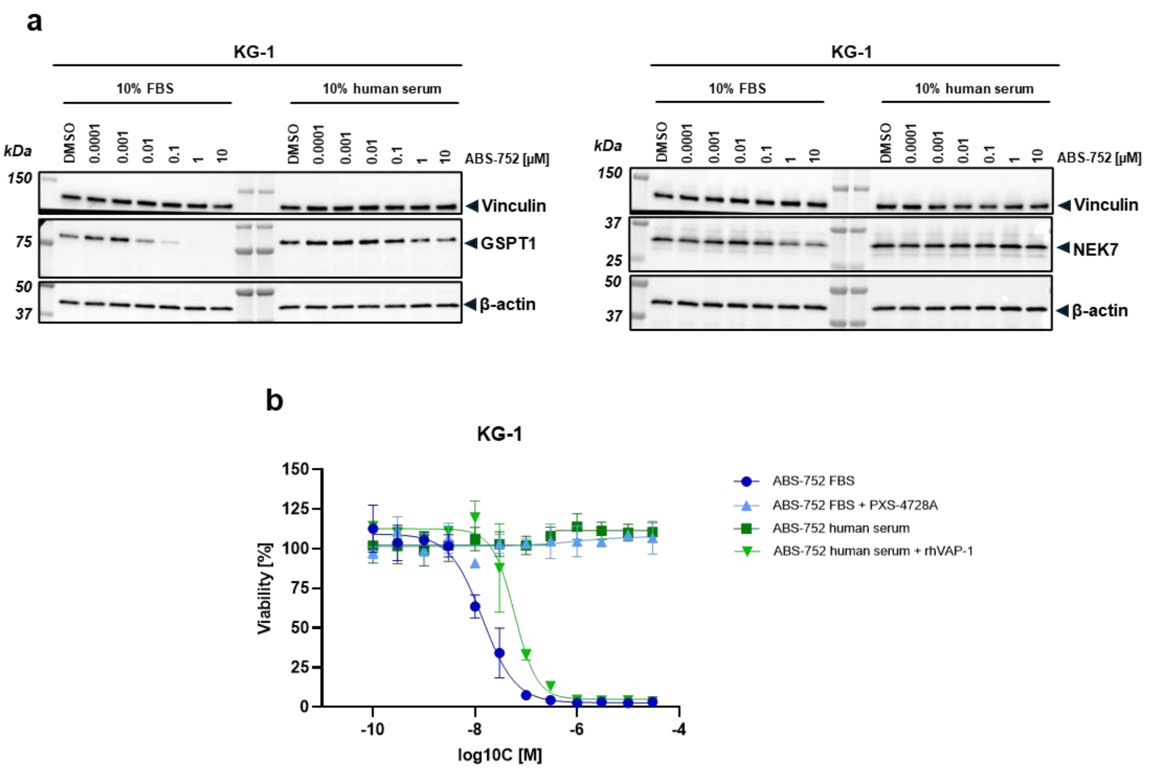

***Figure S6. (a)** Western blot results for GSPT1 and NEK7 degradation after 24-hour post-treatment with ABS-752 in KG-1 cell line supplemented with 10% FBS or 10% human serum. **(b)** CTG viability results for KG-1 cell line supplemented with 10% human serum after 72-hour post-treatment with ABS-752 in a concentration range of 0.1 nM - 30 µM in the presence or absence of 0.3 µM recombinant human VAP-1 protein. Data points are means with standard deviation (the results are of at least two independent replicates).*

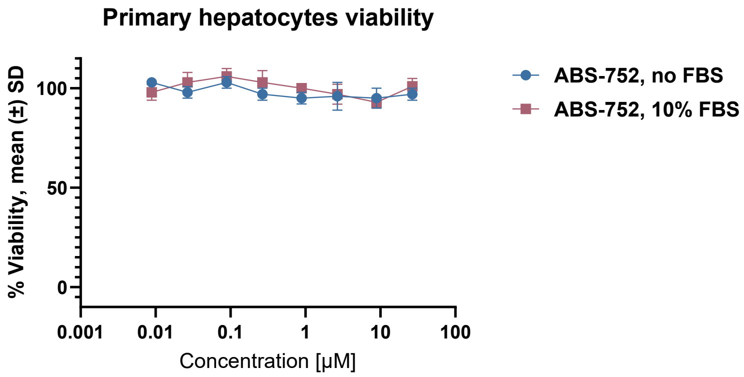

***Figure S7. Cell viability study of primary human hepatocytes treated with ABS-752 in the absence or presence of FBS (CTG assay).***

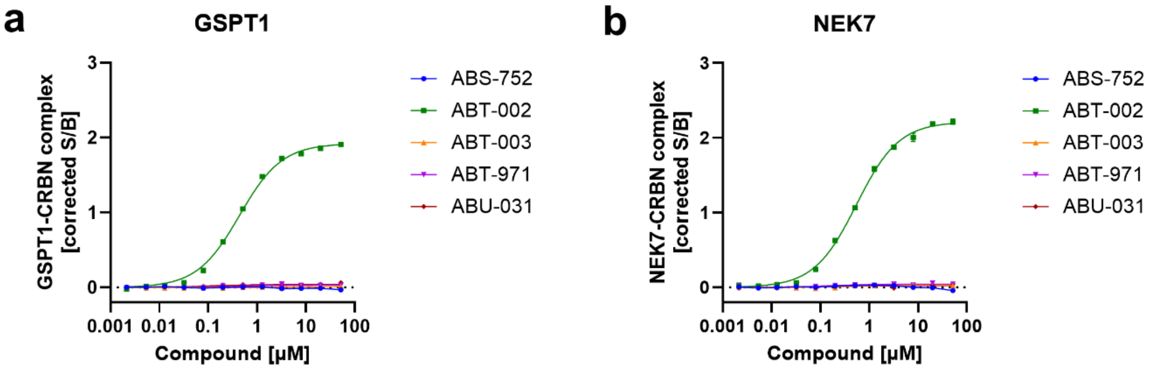

***Figure S8. Analysis of molecular glue activity of ABS-752 and its presumed metabolites in biochemical ternary complex formation assay.*** *Representative results of TR-FRET-based assay for the formation of the GSPT1/CRBN **(a)** and NEK7/CRBN **(b)** ternary complexes as a function of compound concentration (2 nM - 50 µM), data presented are means with standard deviation.*

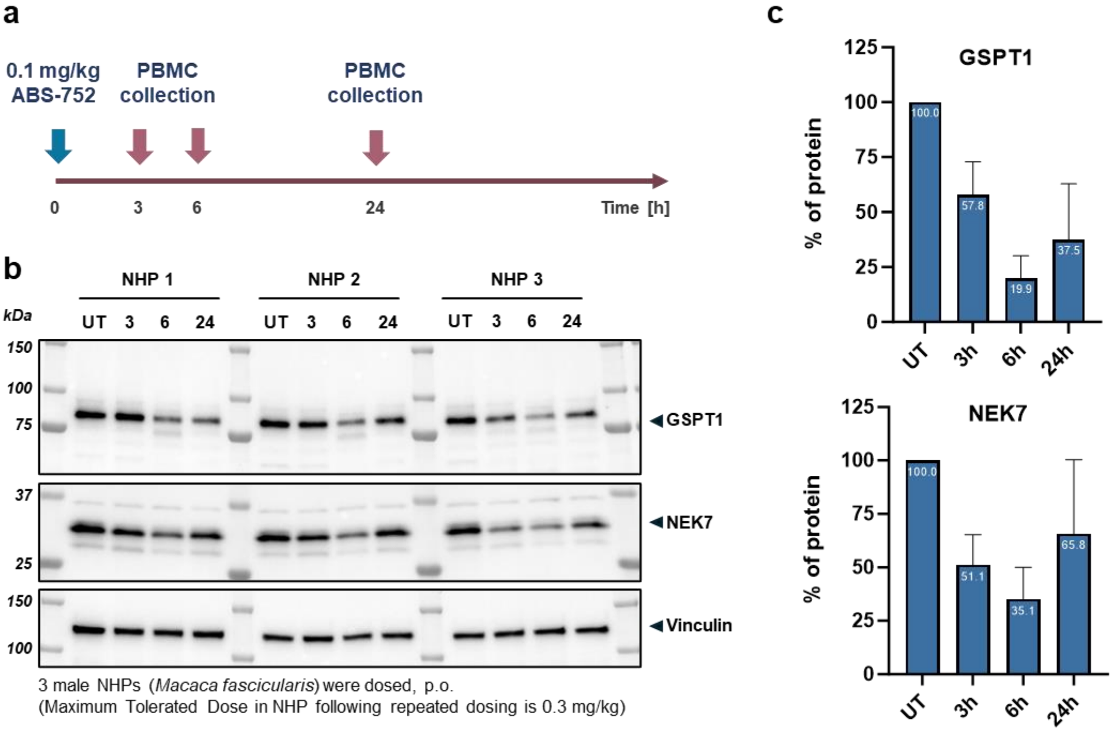

***Figure S9. In vivo PKPD study in non-human primates (NHPs). (a)*** *Design of the experiment: ABS-752 was dosed at 0.1 mg/kg per os to 3 male cynomolgus macaques. Blood samples were collected directly before compound administration (UT), and at 3, 6 & 24h post dose. PBMCs were separated, snap frozen and analyzed by WB. **(b)** Western blot analysis of PBMC lysates. **(c)** GSPT1 and NEK7 protein levels relative to pre-dose, normalized to vinculin loading control.*

### Supplementary Materials and Methods

#### Synthesis of the compounds

The reagents and solvents were used as received from the commercial sources. Proton nuclear magnetic resonance (NMR) spectra were recorded on 500 MHz Bruker Avance spectrometers. The spectra are reported in terms of chemical shift (δ [ppm]), multiplicity (s = singlet, d = doublet, t = triplet, q = quartet, p = quintet, m= multiplet), coupling constant (*J* [Hz]), and integration. Chemical shifts are reported in ppm relative to dimethyl sulfoxide-d6 (δ 2.50) or chloroform-d (δ 7.26) as indicated in NMR spectra data. The samples were prepared by dissolving a dry sample (0.2-2 mg) in an appropriate deuterated solvent (0.7-1 mL).

LCMS measurements were collected using Shimadzu Nexera X2/MS-2020 coupled to liquid chromatograph. All masses reported are the *m/z* of the protonated parent ions unless otherwise stated. The sample was dissolved in an appropriate solvent (e.g. DMSO, ACN, water) and was injected directly into the column using an automated sample handler.

Column: Shim-pack Scepter C18 3.0 µm 300 Å, 150 x 3.0 mm, Mobile Phases: A: water+0.1% FA, B: ACN+0.1% FA, flowrate: 0.5 mL/min

Program 1: 5→95% B (15 min) – 95% B (3 min) – 95→5% B (1 min) – 5% B (6 min)

Program 2: 1% B (2 min) – 1→50% B (5 min) – 50% B (2 min) – 50→1% B (1 min) – 1% B (5 min)

The chemical names were generated using ChemDraw Professional v. 18.2.0.48 from PerkinElmer Informatics, Inc.

Abbreviations used in the following examples are presented below in the alphabetical order:

ACN acetonitrile
DMF *N*,*N*-dimethylformamide
DCM dichloromethane
DMAP 4-(dimethylamino)pyridine
DIPEA *N*-ethyldiisopropylamine
DMSO dimethylsulfoxide
ESI electrospray ionization
FA formic acid
HPLC high performance liquid chromatography
IPA 2-propanol
LCMS Liquid chromatography mass spectrometry
RT Room temperature (temperature of between 20°C and 30°C)
TFA trifluoroacetic acid
THF tetrahydrofuran

3-(5-(Aminomethyl)-1-oxoisoindolin-2-yl)piperidine-2,6-dione hydrochloride (AAC-215; Angene Chemical, AG01JXT0, Lot: AGN21-447835) and 2-(2,6-dioxopiperidin-3-yl)-1-oxoisoindoline-5-carboxylic acid (ABR-522; Enamine, EN300-1653366) were purchased from the commercial sources.

2-(2,6-Dioxopiperidin-3-yl)-1-oxoisoindoline-5-carbaldehyde (ABT-394; Ref: WO2022/29573, 2022), 3-(5-(hydroxymethyl)-1-oxoisoindolin-2-yl)piperidine-2,6-dione (ABR-958; Ref: WO2022/29138, 2022), *N*-[2-(2,6-dioxo-piperidin-3-yl)-1-oxo-2,3-dihydro-1H-isoindole-5-ylmethyl]acetamide (ABR-321; Ref: WO2022/29138, 2022) and 2-(2,6-dioxopiperidin-3-yl)-6-fluoro-1-oxoisoindoline-5-carbonitrile (WO2023/116835, 2023) were prepared according to the previously published procedures.

1. Synthesis of 3-(5-(aminomethyl)-6-fluoro-1-oxoisoindolin-2-yl)piperidine-2,6-dione (ABS-752)

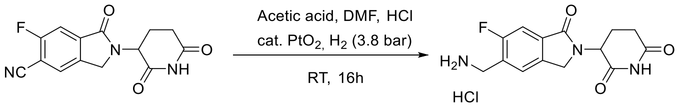

To a solution of 2-(2,6-dioxopiperidin-3-yl)-6-fluoro-1-oxoisoindoline-5-carbonitrile (50 g, 0.17 mol) in acetic acid (300 mL) and DMF (50 mL), and 35% $HCl_{aq}$ (100 mL), and Adams' catalyst (6.0 g, 0.026 mol) were added, and the reaction mixture was stirred under hydrogen atmosphere (3.8 bar) at RT for 16 h. After completion, the reaction mixture was filtered through celite bed, the solids were washed with water (6.20 L) and the filtrates were concentrated under reduced pressure. The crude product was suspended in IPA/$H_2O$ (7:3), stirred at 70-75°C for 1.5 h and cooled to RT. The solids were filtered, washed with IPA and dried under reduced pressure to give 3-(5-(aminomethyl)-6-fluoro-1-oxoisoindolin-2-yl)piperidine-2,6-dione hydrochloride salt (35 g, 61% yield).

$^1$H NMR (500 MHz, DMSO) δ 11.00 (s, 1H), 8.59 (s, 3H), 7.84 (d, *J* = 6.2 Hz, 1H), 7.63 (d, *J* = 8.7 Hz, 1H), 5.13 (dd, *J* = 13.3, 5.1 Hz, 1H), 4.48 (d, *J* = 17.4 Hz, 1H), 4.34 (d, *J* = 17.3 Hz, 1H), 4.16 (s, 3H), 2.92 (ddd, *J* = 17.5, 13.7, 5.4 Hz, 1H), 2.68 – 2.57 (m, 1H), 2.42 (qd, *J* = 13.3, 4.5 Hz, 1H), 2.03 (ddt, *J* = 10.3, 5.2, 3.1 Hz, 2H). $^{13}$C NMR (126 MHz, DMSO) δ 172.81, 170.76, 166.77 (d, *J* = 2.7 Hz), 160.25 (d, *J* = 246.9 Hz), 137.56 (d, *J* = 2.2 Hz), 133.85 (d, *J* = 8.6 Hz), 126.34 (d, *J* = 3.6 Hz), 125.21 (d, *J* = 16.8 Hz), 109.80 (d, *J* = 23.8 Hz), 51.86, 46.99, 36.00 (d, *J* = 4.7 Hz), 31.14, 22.34. LCMS (ESI+) *m/z* 292.2 $[M+1]^+$ (Program 2).

1. Synthesis of 2-(2,6-dioxopiperidin-3-yl)-6-fluoro-1-oxoisoindoline-5-carboxylic acid (ABT-002)

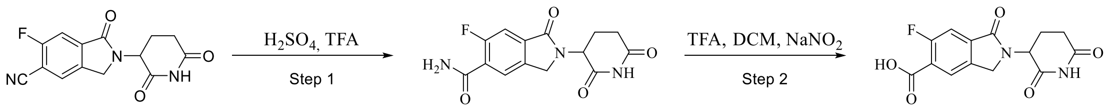

Step 1. 2-(2,6-dioxopiperidin-3-yl)-6-fluoro-1-oxoisoindoline-5-carbonitrile (100 g, 0.348 mol), 98% sulfuric acid (260 mL), TFA (60 mL) were placed in a flask and the reaction mixture was stirred at 80-85°C for 16 h. After completion, the reaction mixture was cooled in an ice bath to 5°C and cold water (1.30 L) was added slowly. The solids were filtered, washed with water and dried in air to give 2-(2,6-dioxopiperidin-3-yl)-6-fluoro-1-oxoisoindoline-5-carboxamide (85 g, 80% yield).

LCMS (ESI+) *m/z* 306.2 $[M+1]^+$ (Program 1).

Step 2. To a solution of 2-(2,6-dioxopiperidin-3-yl)-6-fluoro-1-oxoisoindoline-5-carboxamide (60 g, 0.19 mol) in DCM (480 mL) and TFA (300 mL) at 5°C portionwise sodium nitrite (67.7 g, 0.98 mol) was added and the reaction mixture was stirred for 2 h. Water (120 mL) was added, and DCM and TFA were removed under reduced pressure. Additional water (600 mL) was added, and the solids were filtered. The crude product was suspended in IPA/$H_2O$ (1:1) (600 mL) and stirred at 90°C for 2 h. After cooling, the product was filtered and dried in air to give 2-(2,6-dioxopiperidin-3-yl)-6-fluoro-1-oxoisoindoline-5-carboxylic acid (44 g, 73% yield, purity by HPLC: 99.4%).

$^1$H NMR (500 MHz, DMSO) δ 12.96 (br s, 1H), 11.03 (s, 1H), 8.11 (d, *J* = 6.1 Hz, 1H), 7.64 (d, *J* = 9.4 Hz, 1H), 5.16 (dd, *J* = 13.3, 5.1 Hz, 1H), 4.52 (d, *J* = 17.5 Hz, 1H), 4.40 (d, *J* = 17.4 Hz, 1H), 2.93 (ddd, *J* = 17.5, 13.7, 5.4 Hz, 1H), 2.67 – 2.59 (m, 1H), 2.42 (ddd, *J* = 25.7, 12.9, 4.1 Hz, 1H), 2.09 – 2.01 (m, 1H). $^{13}$C NMR (176 MHz, DMSO) δ 172.82, 170.73, 166.40 (d, *J* = 2.8 Hz), 164.78 (d, *J* = 2.6 Hz), 160.98 (d, *J* = 257.4 Hz), 137.24 (d, *J* = 2.7 Hz), 136.69 (d, *J* = 8.8 Hz), 127.12, 122.71 (d, *J* = 12.4 Hz), 111.17 (d, *J* = 24.5 Hz), 51.99, 47.14, 31.16, 22.37. LCMS (ESI-) *m/z* 305.2 $[M-1]^-$ (Program 1).

2. Synthesis of 2-(2,6-dioxopiperidin-3-yl)-6-fluoro-1-oxoisoindoline-5-carbaldehyde (ABT-971)

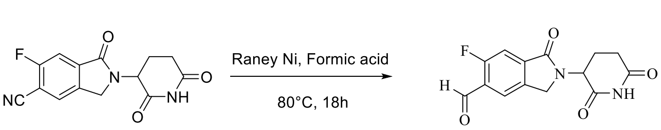

To a solution of 2-(2,6-dioxopiperidin-3-yl)-6-fluoro-1-oxoisoindoline-5-carbonitrile (20 g, 69 mmol) in formic acid (200 mL) Raney Nickel (20.0 g, 0.24 mol) at RT was added. The reaction mixture was then stirred at 80°C for 18 h. After cooling, the reaction mixture was filtered through celite bed and the solids were washed with DCM (2x100 mL). The combined filtrates were washed with water (2x100 mL), the organic layer was concentrated under reduced pressure and the crude product was purified by preparative

HPLC to give 2-(2,6-dioxopiperidin-3-yl)-6-fluoro-1-oxoisoindoline-5-carbaldehyde (12.1 g, 60% yield, purity by HPLC: 98.8%).

$^1$H NMR (500 MHz, DMSO) δ 11.02 (s, 1H), 10.32 (s, 1H), 8.09 (d, *J* = 5.8 Hz, 1H), 7.74 (d, *J* = 9.3 Hz, 1H), 5.15 (dd, *J* = 13.3, 5.1 Hz, 1H), 4.53 (d, *J* = 17.5 Hz, 1H), 4.42 (d, *J* = 17.5 Hz, 1H), 2.91 (ddd, *J* = 17.5, 13.7, 5.4 Hz, 1H), 2.66 – 2.56 (m, 1H), 2.41 (qd, *J* = 13.2, 4.4 Hz, 1H), 2.04 (dtd, *J* = 12.6, 5.3, 2.2 Hz, 1H). $^{13}$C NMR (176 MHz, DMSO) δ 187.81 (d, *J* = 5.9 Hz), 172.80, 170.64, 166.22 (d, *J* = 2.8 Hz), 163.46 (d, *J* = 257.7 Hz), 138.28 (d, *J* = 9.2 Hz), 137.68 (d, *J* = 2.5 Hz), 126.19 (d, *J* = 9.8 Hz), 124.45, 111.16 (d, *J* = 22.8 Hz), 52.06, 47.27, 31.13, 22.31. LCMS (ESI-) *m/z* 289.0 $[M-1]^-$ (Program 1).

3. Synthesis of 3-(6-fluoro-5-(hydroxymethyl)-1-oxoisoindolin-2-yl) piperidine-2,6-dione (ABU-031)

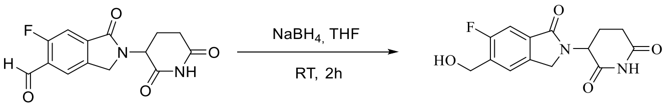

To a solution of 2-(2,6-dioxopiperidin-3-yl)-6-fluoro-1-oxoisoindoline-5-carbaldehyde (5.0 g, 17.2 mmol) in THF (50 mL) portionwise sodium borohydride (6.54 g, 0.172 mol) was added and the resulting mixture was stirred at RT for 2 h. 1M $HCl_{aq}$ was added slowly to quench the excess of the reagent. The mixture was filtered through Celite bed, the solids were washed with DCM (2 x 10 mL) and the combined filtrates were concentrated under reduced pressure. The crude product was purified by preparative HPLC to give 3-(6-fluoro-5-(hydroxymethyl)-1-oxoisoindolin-2-yl)piperidine-2,6-dione (2.50 g, 50% yield, purity by HPLC: 99.6%).

$^1$H NMR (500 MHz, DMSO) δ 10.98 (s, 1H), 7.71 (d, *J* = 6.2 Hz, 1H), 7.46 (d, *J* = 8.9 Hz, 1H), 5.47 (t, *J* = 5.6 Hz, 1H), 5.11 (dd, *J* = 13.3, 5.1 Hz, 1H), 4.65 (d, *J* = 5.6 Hz, 2H), 4.45 (d, *J* = 17.1 Hz, 1H), 4.32 (d, *J* = 17.1 Hz, 1H), 2.91 (ddd, *J* = 17.4, 13.7, 5.4 Hz, 1H), 2.63 – 2.57 (m, 1H), 2.39 (qd, *J* = 13.3, 4.5 Hz, 1H), 2.02 (dtd, *J* = 12.6, 5.3, 2.2 Hz, 1H). $^{13}$C NMR (176 MHz, DMSO) δ 172.87, 170.92, 167.32 (d, *J* = 3.1 Hz), 159.42 (d, *J* = 244.9 Hz), 137.78 (d, *J* = 2.3 Hz), 133.88 (d, *J* = 16.6 Hz), 131.85 (d, *J* = 8.6 Hz), 123.51 (d, *J* = 4.9 Hz), 108.98 (d, *J* = 23.5 Hz), 56.92 (d, *J* = 5.1 Hz), 51.80, 47.06, 31.20, 22.48. LCMS (ESI+) *m/z* 293.0 $[M+1]^+$ (Program 1).

4. Synthesis of *N*-((2-(2,6-dioxopiperidin-3-yl)-6-fluoro-1-oxoisoindolin-5-yl)methyl)acetamide (ABT-003)

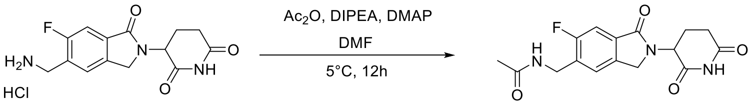

To a solution of 3-(5-(aminomethyl)-6-fluoro-1-oxoisoindolin-2-yl)piperidine-2,6-dione hydrochloride (5.0 g, 15.2 mmol) in DMF (50 mL) DIPEA (5.3 mL, 30.4 mmol) and DMAP (0.18 g, 1.5 mmol) were added, the solution was cooled to 0-5°C and acetic anhydride (1.44 mL, 15.2 mmol) was added dropwise. The reaction mixture was stirred for 12 h at RT and concentrated under reduced pressure. The crude product was purified by preparative HPLC to give *N*-((2-(2,6-dioxopiperidin-3-yl)-6-fluoro-1-oxoisoindolin-5-yl)methyl)acetamide (2.50 g, 50% yield, purity by HPLC: 98.8%).

$^{1}$H NMR (500 MHz, DMSO) δ 10.98 (s, 1H), 8.43 (t, *J* = 5.8 Hz, 1H), 7.54 (d, *J* = 6.3 Hz, 1H), 7.49 (d, *J* = 8.8 Hz, 1H), 5.10 (dd, *J* = 13.3, 5.1 Hz, 1H), 4.43 (d, *J* = 17.2 Hz, 1H), 4.36 (d, *J* = 5.8 Hz, 2H), 4.30 (d, *J* = 17.2 Hz, 1H), 2.91 (ddd, *J* = 17.4, 13.7, 5.5 Hz, 1H), 2.60 (dt, *J* = 17.2, 1.7 Hz, 1H), 2.38 (qd, *J* = 13.5, 4.6 Hz, 1H), 2.01 (ddq, *J* = 10.4, 5.3, 3.1, 2.6 Hz, 1H), 1.89 (s, 3H). $^{13}$C NMR (176 MHz, DMSO) δ 172.86, 170.88, 169.50, 167.17 (d, *J* = 3.2 Hz), 160.00 (d, *J* = 245.8 Hz), 137.70 (d, *J* = 2.2 Hz), 132.13 (d, *J* = 8.8 Hz), 130.78 (d, *J* = 16.6 Hz), 124.45 (d, *J* = 4.6 Hz), 109.29 (d, *J* = 23.8 Hz), 51.82, 47.01, 36.37 (d, *J* = 4.9 Hz), 31.18, 22.49, 22.44. LCMS (ESI+) *m/z* 334.0 $[M+1]^+$ (Program 1).

#### NMR spectra of the compounds

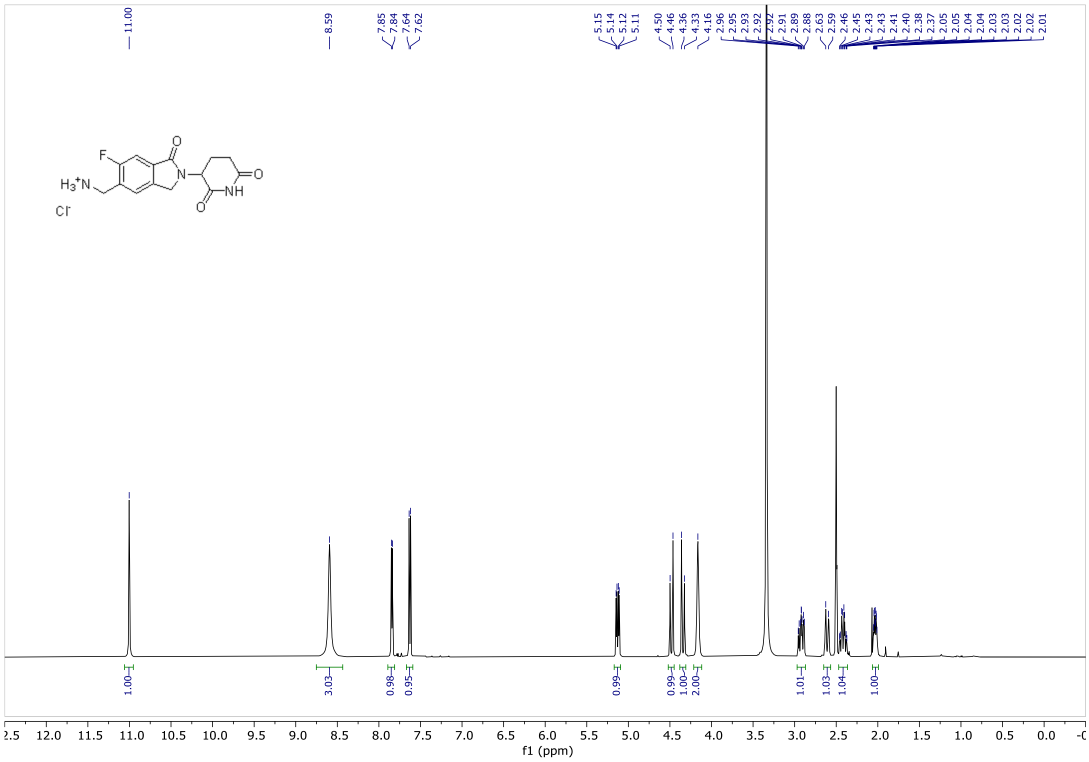

***Figure S10.** $^{1}$H NMR of 3-(5-(aminomethyl)-6-fluoro-1-oxoisoindolin-2-yl)piperidine-2,6-dione.*

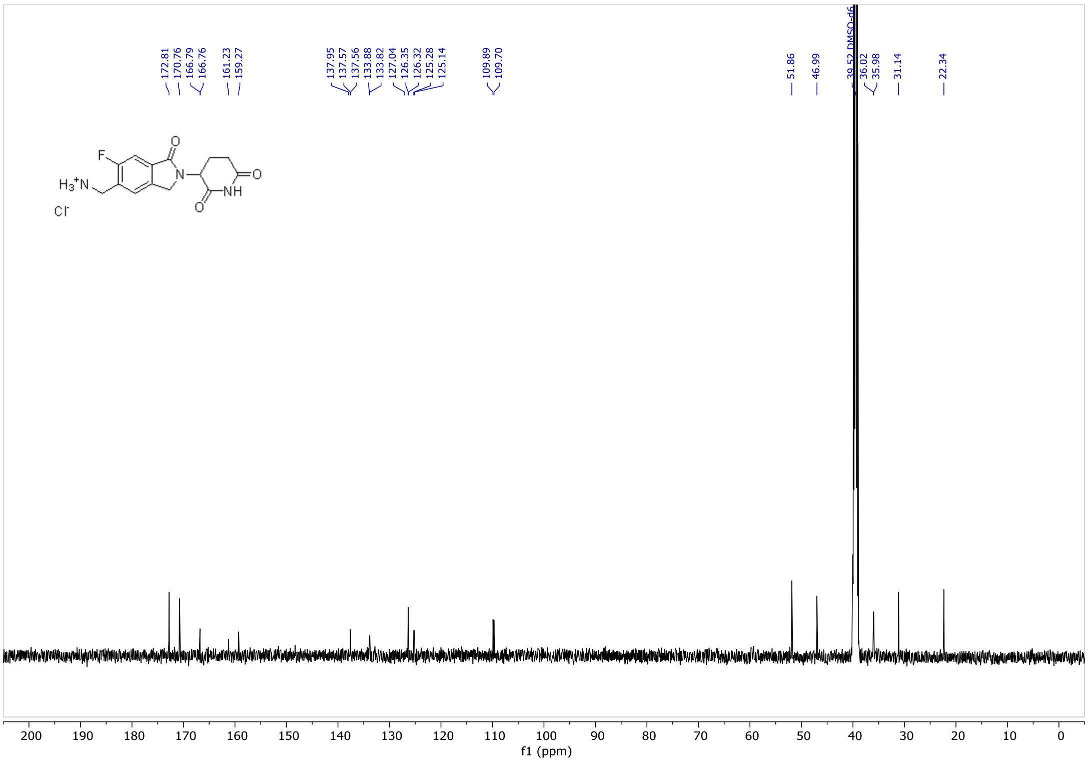

***Figure S11. $^{13}$C NMR of 3-(5-(aminomethyl)-6-fluoro-1-oxoisoindolin-2-yl)piperidine-2,6-dione.***

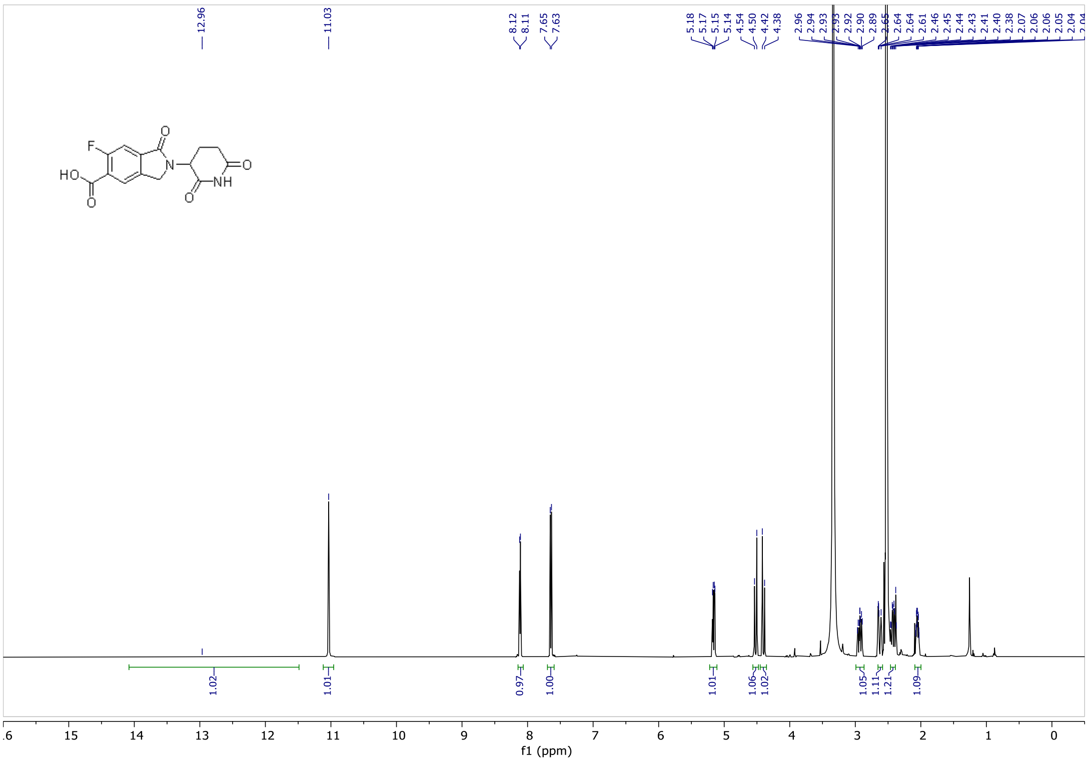

***Figure S12. $^{1}$H NMR of 2-(2,6-dioxopiperidin-3-yl)-6-fluoro-1-oxoisoindoline-5-carboxylic acid.***

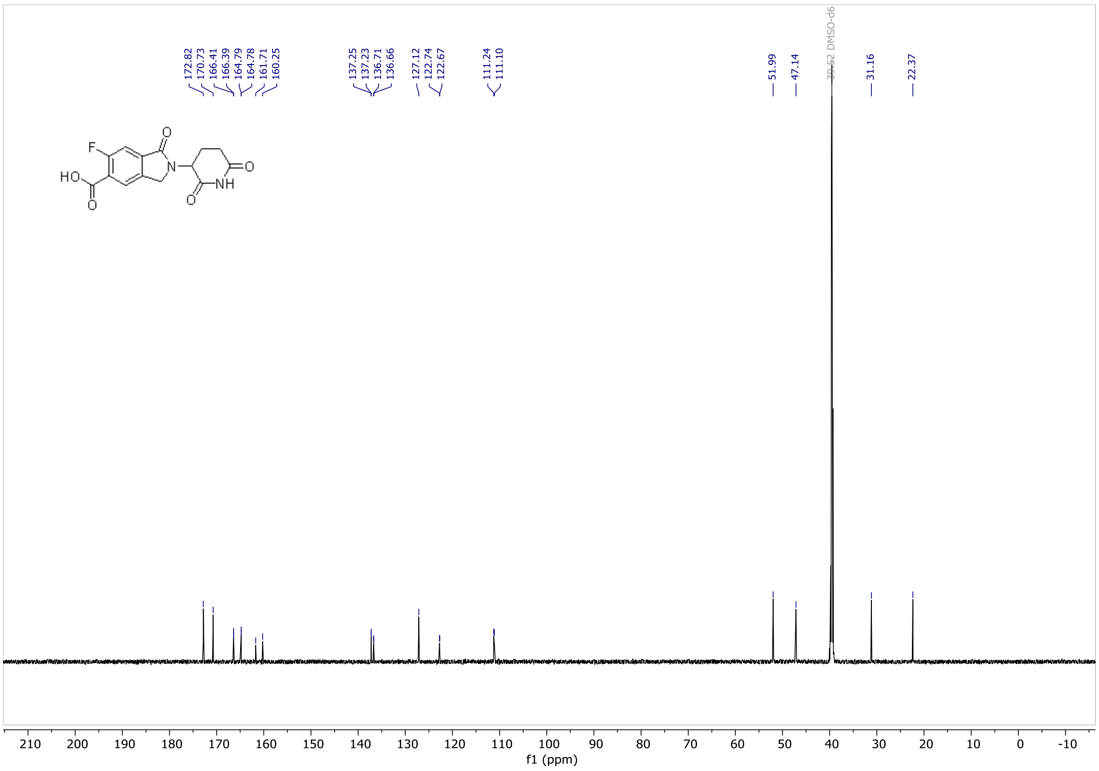

***Figure S13. 1H NMR of 2-(2,6-dioxopiperidin-3-yl)-6-fluoro-1-oxoisoindoline-5-carboxylic acid.***

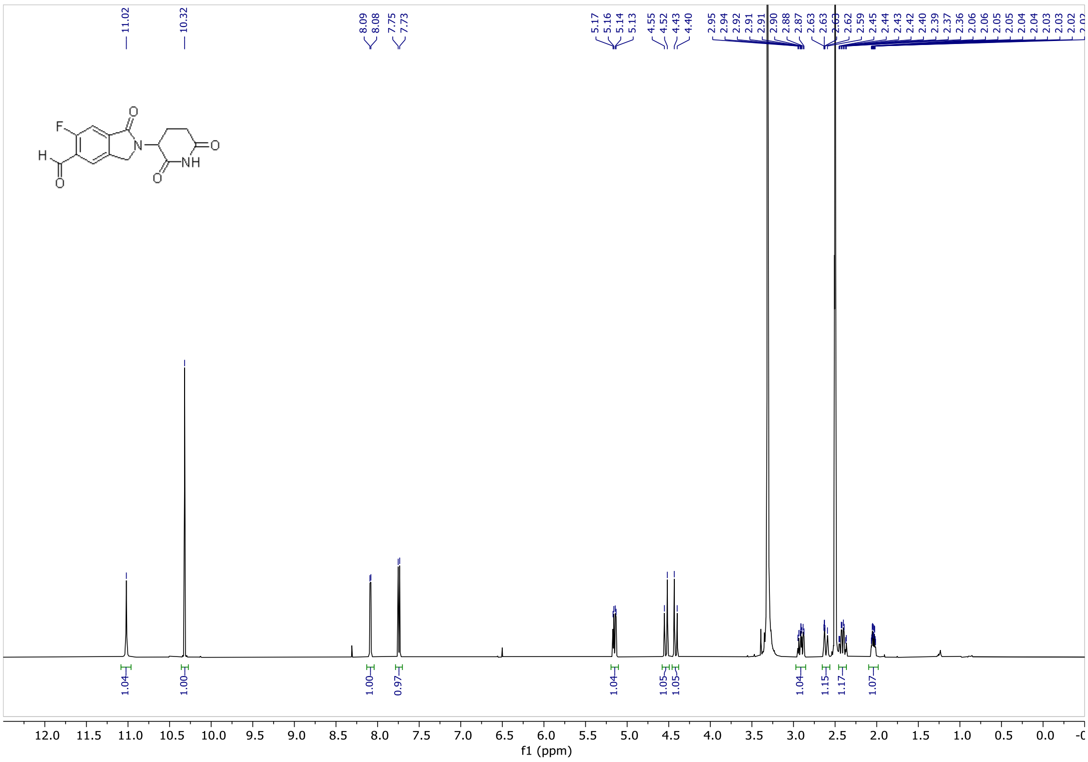

***Figure S14. 1H NMR of 2-(2,6-dioxopiperidin-3-yl)-6-fluoro-1-oxoisoindoline-5-carbaldehyde.***

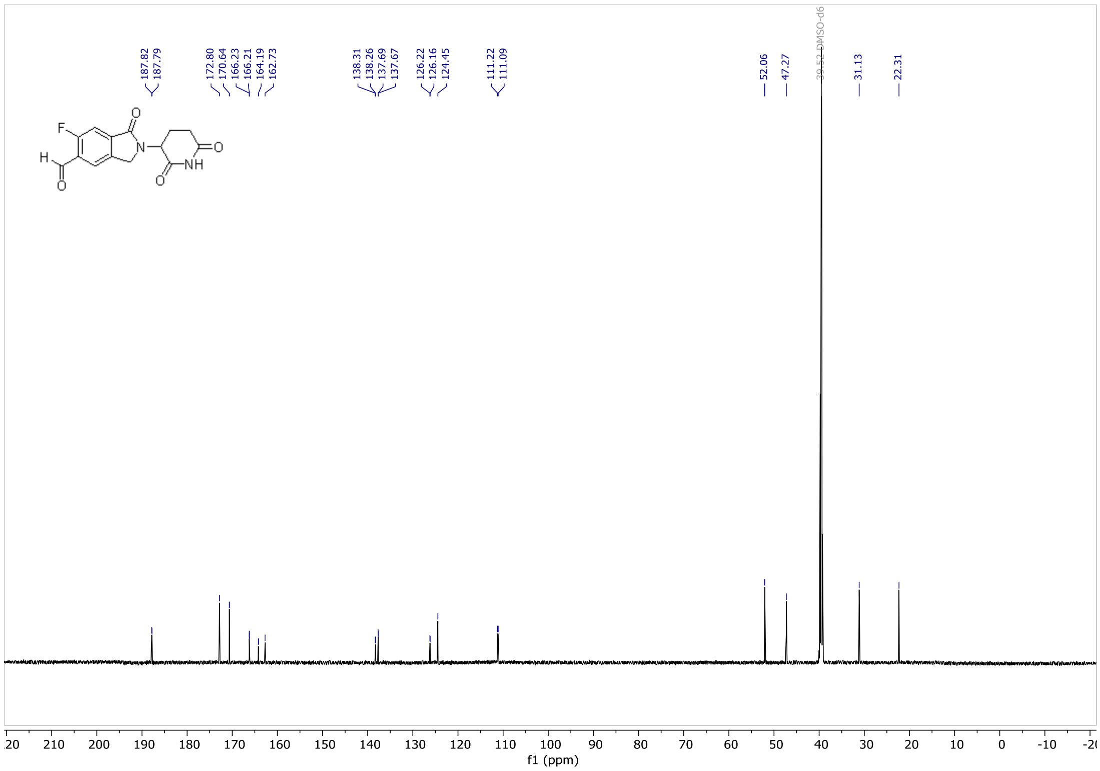

***Figure S15.** *$^{13}$*C NMR of 2-(2,6-dioxopiperidin-3-yl)-6-fluoro-1-oxoisoindoline-5-carbaldehyde.**

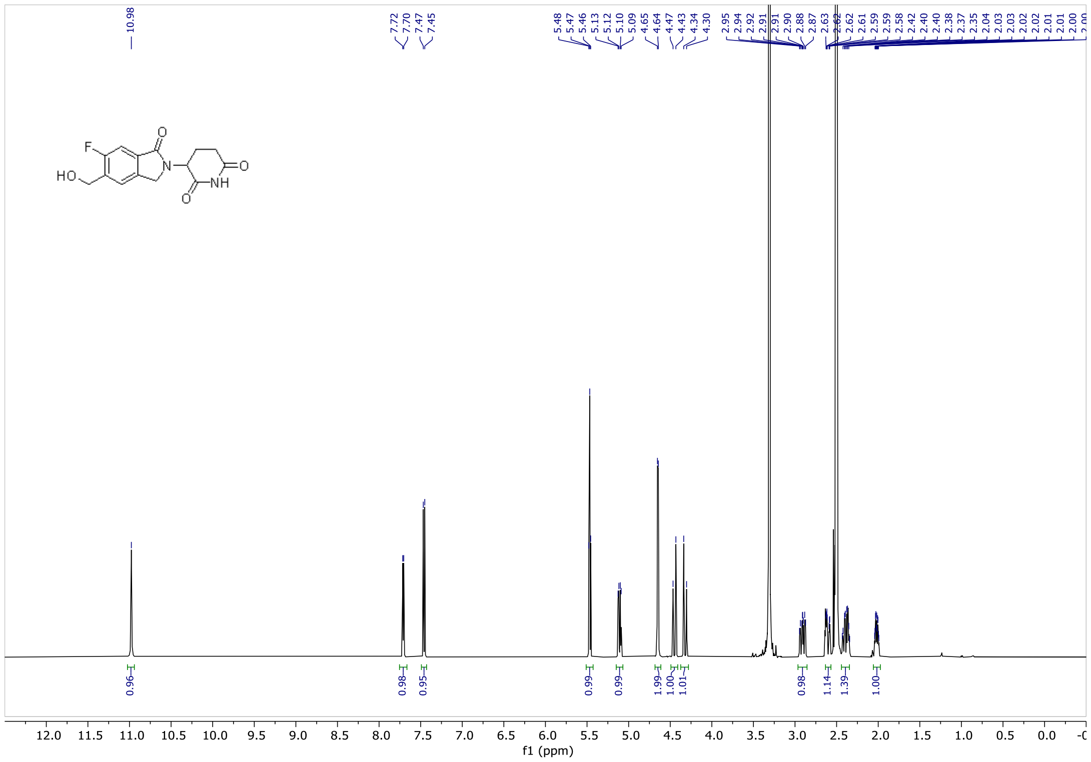

***Figure S16.** *$^{1}$*H NMR of 3-(6-fluoro-5-(hydroxymethyl)-1-oxoisoindolin-2-yl) piperidine-2,6-dione.**

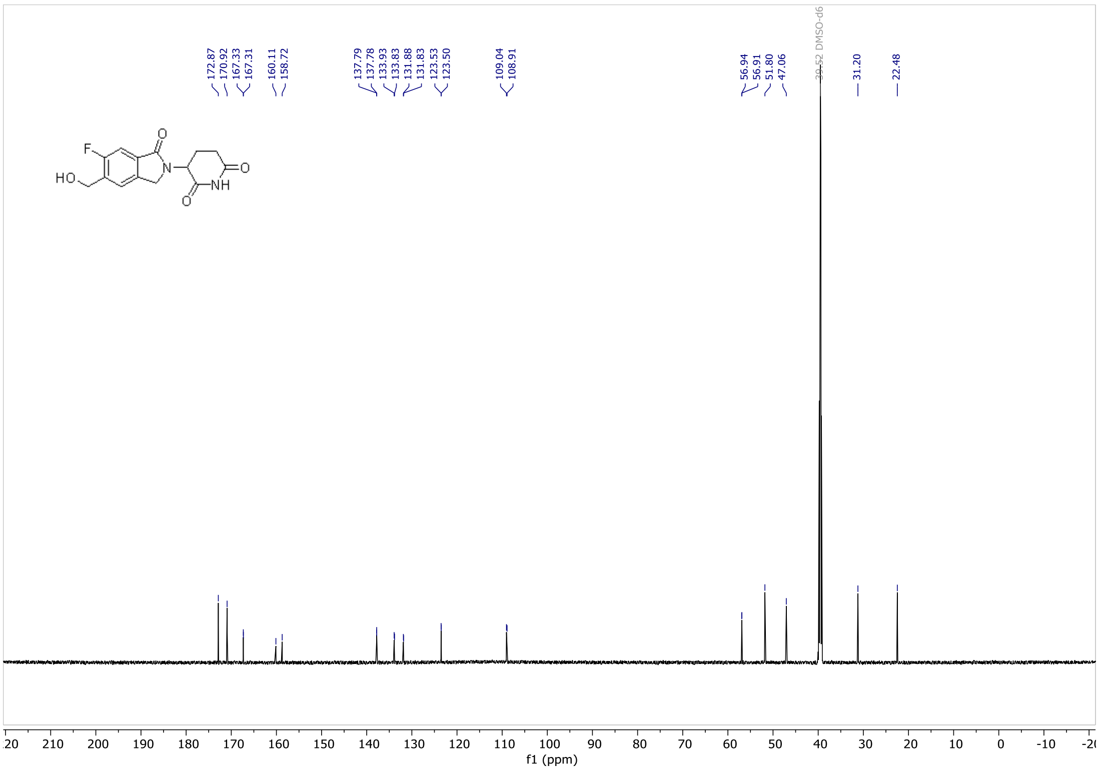

***Figure S17. $^{13}$C NMR of 3-(6-fluoro-5-(hydroxymethyl)-1-oxoisoindolin-2-yl) piperidine-2,6-dione.***

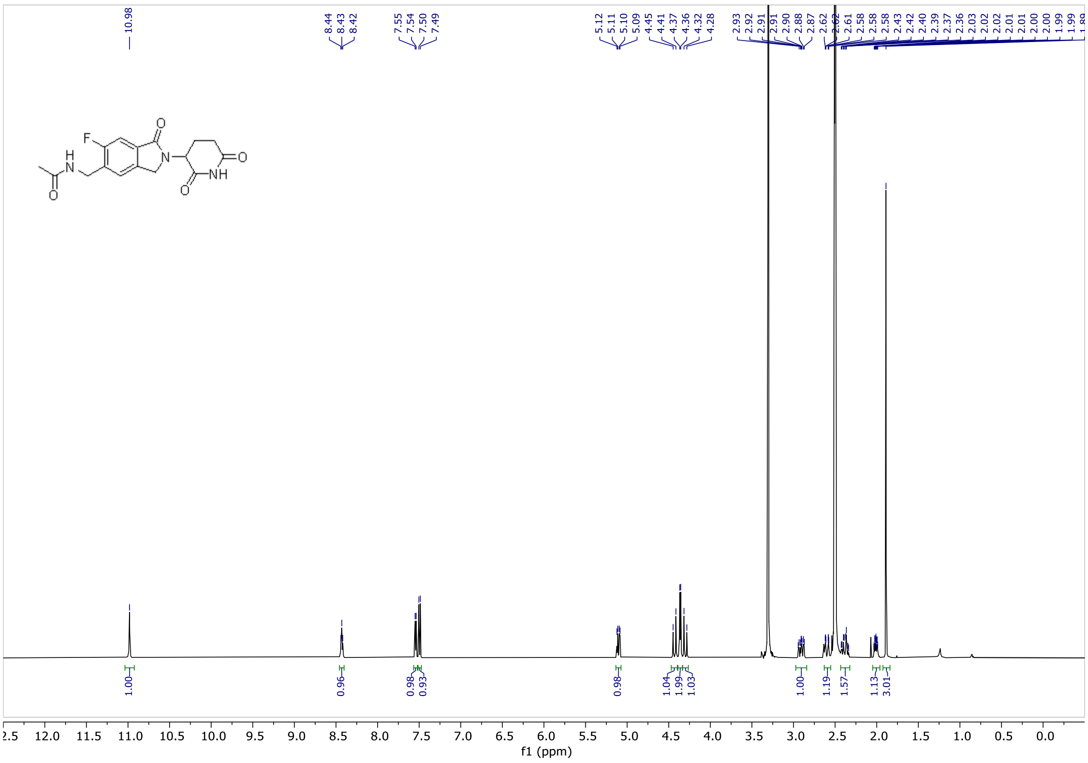

***Figure S18. $^{1}$H NMR of N-((2-(2,6-dioxopiperidin-3-yl)-6-fluoro-1-oxoisoindolin-5-yl)methyl)acetamide.***

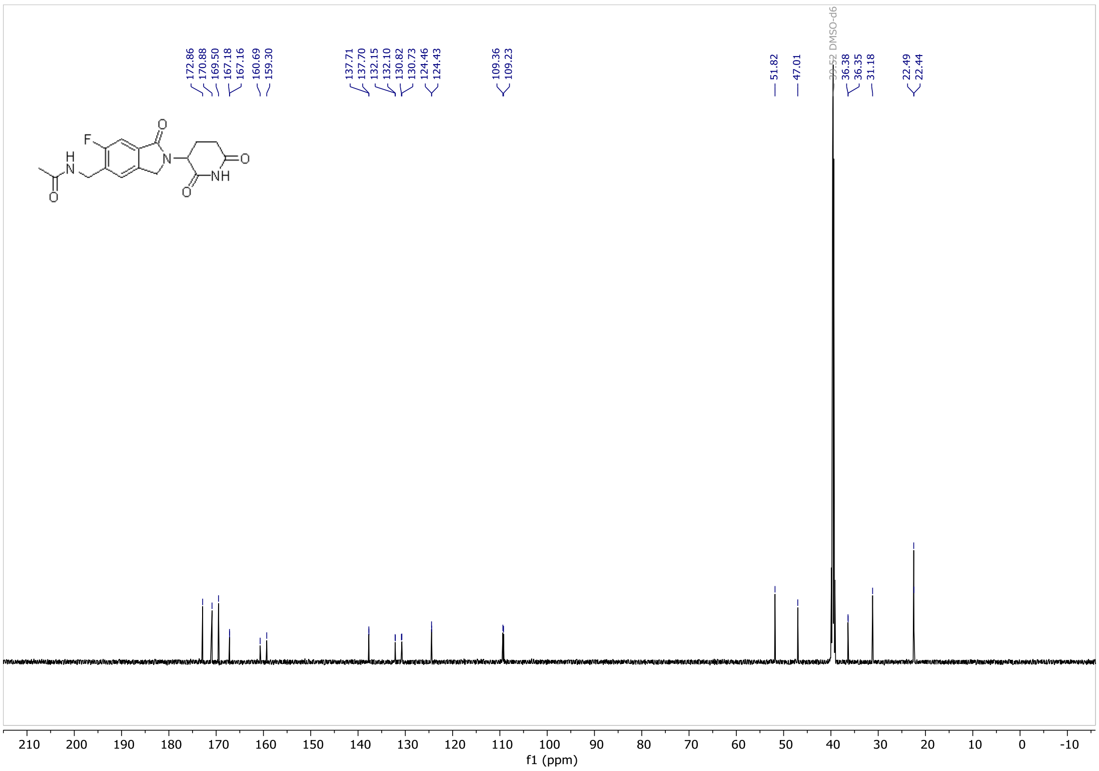

***Figure S19.*** ***$^{13}$C NMR of N-((2-(2,6-dioxopiperidin-3-yl)-6-fluoro-1-oxoisoindolin-5-yl)methyl)acetamide.***

#### Purification of recombinant proteins

SALL4, IKFZ1, NEK7 and GSPT1 were produced in *E. coli* based expression system, whereas all CRBN-DDB1 constructs and CK1α were expressed using the Bac-to-Bac baculovirus system in *Spodoptera frugiperda* Sf9 cells (Expression Systems) or *Trichoplusia ni* High Five cells (Thermo Fischer Scientific), as indicated below.

**SALL4 ZF2** (406-432) and **IKFZ1 ZF2** (141-174) tagged N-terminally with StrepTagII, AviTag and SUMO were cloned into pET28 vector and expressed in *E. coli* BL21(DE3) cells in LB medium (3 hours post induction, at 37°C). Cells were disintegrated via pressure drops using EmulsiFlex microfluidizer (Avestin), in 50 mM Tris/HCl (pH 8.0, 8°C), 300 mM NaCl, 5% glycerol, cOmplete EDTA-free inhibitor cocktail (Roche), 0.05% Triton X-100, 5 U/mL Viscolase (A&A Biotechnology), supplemented with 1 mM DTT. Recombinant proteins were captured from lysate with Streptactin XT 4flow high capacity resin (IBA Lifesciences) and eluted with buffer A with 5 mM biotin and 1mM DTT. POI was diluted to lower the NaCl concentration to ± 30 mM and purified further with Anion-Exchange Chromatography (HiTrap Q HP, Cytiva) and eluted in 30-500 mM NaCl gradient. In case of SALL4 ZF2 additional Size-Exclusion Chromatography polishing step was performed using HiLoad 16/600 Superdex 75 pg (Cytiva) in 50 mM Tris-HCl pH 8.0, 300 mM NaCl, 1 mM DTT.

**NEK7** (full-length protein, 1-302) tagged N-terminally with HisTag and AviTag enabling site-specific biotinylation was cloned into pET28 vector and expressed in *E. coli* BL21(DE3) cells in LB medium supplemented with 0.5% glucose (16 hours post induction, at 16°C). Cells were disintegrated using EmulsiFlex microfluidizer (Avestin), in 50 mM Tris/HCl (pH 8.0, 8°C), 300 mM NaCl, 5% glycerol, cOmplete EDTA-free inhibitor cocktail, 0.05% Triton X-100, 5 U/mL Viscolase with 10 mM βME and 20 mM imidazole. POI was captured from lysate Ni Sepharose 6 Fast Flow (Cytiva) and eluted with buffer A with 250 mM imidazole. Next, the POI was cleaved with HRV 3C Protease and tags were removed using HisTrap HP (Cytiva). The cleaved protein was diluted to lower the NaCl concentration to ± 30 mM and purified with Anion-Exchange Chromatography (HiTrap Q HP, Cytiva) with 300 mM NaCl step elution. Recombinant protein was enzymatically biotinylated using GST-tagged BirA ligase and Size-Exclusion Chromatography polishing step was performed using HiLoad 26/600 Superdex 75 pg (Cytiva). The POI was formulated in 20 mM HEPES pH 7.5, 200 mM NaCl, 5% glycerol, 1 mM DTT.

**GSPT1** fragment (300-496) containing the degron sequence was cloned into pET28 vector, with N-terminal HisTag and MBP as a solubility tag, followed by the TEV protease cleavage site and AviTag for biotinylation. GSPT1 was expressed in *E. coli* BL21(DE3) cells in Dynamite, a modified Studier's auto-induction medium (16 hours post induction, at 16°C). Cells were disintegrated using EmulsiFlex microfluidizer (Avestin) in 50 mM Tris/HCl (pH 8.0, 8°C), 300 mM NaCl, 5% glycerol, cOmplete EDTA-free inhibitor cocktail, 0.05% Triton X-100, 5 U/mL Viscolase, 1 mM DTT, centrifuged (30 000 rcf, 4°C, 30 min) and PEI solution was added to the supernatant fraction to a final concentration of 0.1% (v/v) and re-centrifuged (50 000 rcf, 4°C, 30 min). Recombinant protein was captured from lysate with IMAC (Immobilized Metal Affinity Chromatography) using Ni-NTA resin (Cytiva) and eluted with 500 mM imidazole in buffer A. To remove imidazole, the sample was subjected to buffer exchange using HiPrep 26/10 Desalting column (Cytiva). Protein was cleaved overnight with TEV protease at 4°C (1:50 mass ratio) and cleaved POI was separated from tags and protease using reverse IMAC on HisTrap HP column (Cytiva). For biotinylated protein, the sample was enzymatically biotinylated using GST-tagged BirA ligase. For non-biotinylated protein batch, to remove any endogenously biotinylated fraction of the POI, sample was subjected to reverse StrepAC using Strep-Tactin XT 4Flow high capacity (IBA Lifesciences) and the flow-through sample containing non-biotinylated POI was collected. Both biotinylated and non-biotinylated protein samples were subjected to SEC using HiLoad 16/600 Superdex 75 pg (Cytiva) in PBS pH = 7.4 (Sigma-Aldrich), 1 mM DTT.

For **CRBN-DDB1-CaptorBait**, wild-type and full-length human CRBN and human full-length DNA damage-binding protein 1 (DDB1) genes were subcloned into the pFastBac Dual vector, followed by the preparation of baculovirus in *E. coli* DH10Bac cells. CRBN was tagged with an N-terminal HisTag (8xHis) sequence, followed by an HRV 3C protease cleavage site. DDB1 was fused to an N-terminal StrepTagII, also followed by an HRV 3C protease cleavage site and the CaptorBait peptide[1]. Sf9 cells were infected with the baculovirus, harvested 48 hours post-infection and lysed via pressure drops using EmulsiFlex microfluidizer (Avestin) in 50 mM Tris/HCl (pH 8.0, 8°C), 300 mM NaCl, 5% glycerol, cOmplete EDTA-free inhibitor cocktail, 0.05% Triton X-100, 5 U/mL Viscolase with addition of 1 mM $MgCl_2$, 10 µM $ZnCl_2$, 5 mM sodium citrate, 20 mM imidazole and 5 mM β-mercaptoethanol (β-ME).

The CRBN-DDB1 complex was isolated from the lysate using Ni-INDIGO resin (Cube Biotech). After washing the resin with lysis buffer supplemented with 20 mM imidazole protein was eluted with the same buffer containing 0.3 M imidazole and the eluate was diluted 10x with 50 mM Tris/HCl (pH 8.0, RT), 1 mM DTT followed by direct load on HiTrap Q HP 5 mL column (Cytiva). Resin was washed with 50 mM Tris/HCl (pH 8.0, RT), 290 mM NaCl, 1 mM DTT to remove impurities and buffer containing ~365 mM NaCl was used to elute CRBN-DDB1 CaptorBait complex. The salt concentration corresponded to conductivity equal 31 and 37 mS/cm, respectively. Final polishing was performed using size-exclusion chromatography with a HiLoad Superdex 16/600 200 pg column (Cytiva). The complex was stored in a buffer composed of 50 mM HEPES/NaOH (pH 8.0; RT), 200 mM NaCl, 5% glycerol, 1 mM TCEP.

**CRBN-DDB1ΔB-CaptorBait** protein was prepared with the use of DDB1 sequence lacking amino acids 396-705 (corresponding to DDB1 β-propeller B), which were substituted with a GNGNSG linker. CRBN was tagged with an an N-terminal StrepTagII with HRV 3C protease cleavage site. DDB1 contained StrepTagII, HRV 3C protease cleavage site and CaptorBait sequence introduced at its N terminus. Similarly to CRBN-DDB1-CaptorBait, pFastBac Dual vector was used and the proteins were expressed in High Five cells, for 72 hours post-infection. Afterwards cells were harvested and submitted to lysis in 50 mM Tris/HCl (pH 8.0, 8°C), 300 mM NaCl, 5% glycerol, cOmplete EDTA-free inhibitor cocktail, 0.05% Triton X-100, 5 U/mL Viscolase, 1 mM DTT and POI was captured using Strep-Tactin affinity chromatography, eluted in lysis buffer supplemented with 10 mM biotin, diluted to final concentration of 20mM NaCl and further purified by anion-exchange chromatography (AIEX) using HiTrap HP Q column (Cytiva). POI were eluted using linear gradient of 2-60% of 50 mM Tris-HCl (pH 8.0, RT), 1M NaCl, 1 mM DTT, 1 mM TCEP. CRBN-DDB1 was finally purified by size-exclusion chromatography on HiLoad Superdex 16/600 200 pg column (Cytiva) and stored in buffer composed of 50 mM Tris-HCl (pH 8.0, RT), 200 mM NaCl, 1 mM TCEP, 5% glycerol.

For crystallographic studies, a CRBN construct with truncated N-terminus was used (**CRBN(40-442)** together with full-length **DDB1**). CRBN was N-terminally tagged with StrepTagII, HisTag and GST-like protein from *S. frugiperda*, followed with HRV 3C protease cleavage site, HisTag, AviTag and TEV protease cleavage site, so that all solubility and affinity tags were removed during recombinant protein preparation, with only glycine residue remaining at the N-terminus of CRBN. POI was cloned into pFastBac Dual vector and after baculovirus preparation expressed in High Five cells for 72 hours post-infection. Cells were harvested and lysed in 50 mM Tris/HCl (pH 8.0, 8°C), 300 mM NaCl, 5% glycerol, cOmplete EDTA-free inhibitor cocktail, 0.05% Triton X-100, 5 U/mL Viscolase, 1 mM DTT. POI was captured using Strep-Tactin affinity chromatography and eluted in lysis buffer supplemented with 10 mM biotin. Eluate was bound to HisTrap column (Cytiva), washed with lysis buffer with 1M NaCl and eluted with 500 mM imidazole. Protein was dialyzed and cleaved overnight with TEV protease at 4°C (1:50 mass ratio) and cleaved POI was separated from tags and protease using AIEX on HiTrap HP Q column (Cytiva).

The DNA sequence encoding human **CK1α** (residues 1–323) was inserted into the pFastBac Dual vector, and CK1α was tagged with N-terminal StrepII-AviTag. Protein expression was carried out in *Trichoplusia ni* High-Five insect cells using the baculovirus expression system (Invitrogen). The cells were lysed using an EmulsiFlex homogenizer in 50 mM Tris/HCl (pH 8.0, 8°C), 300 mM NaCl, 5% glycerol, cOmplete EDTA-free inhibitor cocktail, 0.05% Triton X-100, 5 U/mL Viscolase, supplemented with 10 mM β-ME. After centrifugation, the soluble fraction was incubated with Strep-Tactin XT Sepharose (IBA), and the eluate was diluted and further purified using cation-exchange chromatography with phosphate-based buffers at pH 7.0. The protein was then biotinylated using GST-BirA, by adding 1 mM ATP, 5 mM $MgCl_2$, and 50 μM biotin. Following overnight biotinylation, the sample was purified by Size-Exclusion Chromatography on HiLoad 16/600 Superdex 75 pg (Cytiva) in 50 mM HEPES (pH 7.4), 250 mM NaCl, and 0.5 mM TCEP.

#### Degradation assays and Western Blot

***Table S7. List of the cell lines used in experiments including culture conditions.***

| Cell line | Source (ref#) | Growth property | Culture medium and centrifuge protocol |
|---|---|---|---|
| **Hep 3B2.1-7 [Hep3B]** | ATCC (HB-8064) | adherent | Eagle's Minimum Essential Medium (ATCC)<br>10% Fetal Bovine Serum, qualified, heat inactivated (Gibco)<br>1% Penicillin/Streptomycin (Biowest) |
| **JHH-7** | JCRB BioBank (JCRB1031) | adherent | William's E Medium, GlutaMAX™ (Gibco)<br>10% Fetal Bovine Serum, qualified, heat inactivated (Gibco)<br>1%Penicillin/Streptomycin (Biowest) |
| **HuH-7** | JCRB BioBank (JCRB0403) | adherent | Dulbecco's modified Eagle's medium (low glucose) (Gibco)<br>10% Fetal Bovine Serum, qualified, heat inactivated (Gibco)<br>1% Penicillin/Streptomycin (Biowest) |
| **HuH-1** | JCRB BioBank (JCRB0199) | adherent | |
| **KG-1** | DSMZ (ACC14) | non-adherent | RPMI 1640 medium (Gibco)<br>10% Fetal Bovine Serum, qualified, heat inactivated (Gibco)<br>1% Penicillin/Streptomycin (Biowest) |
| **Kelly** | DSMZ (ACC355) | adherent | |
| **SNU-886** | Korean Cell Line Bank (KCLB00886) | adherent | |
| **SNU-398** | ATCC (CRL-2233) | Mixed (adherent and non-adherent cells) | |
| **SNU-182** | ATCC (CRL2235) | adherent | RPMI 1640 (ATCC modified) medium (Gibco)<br>10% Fetal Bovine Serum, qualified, heat inactivated (Gibco)<br>1% Penicillin/Streptomycin (Biowest) |
| **SNU-423** | ATCC (CRL-2238) | adherent | |
| **NCI-H929 [H929]** | ATCC (CRL-9068) | non-adherent | RPMI 1640 (ATCC modification) medium (Gibco)<br>10% Fetal Bovine Serum, qualified, heat inactivated (Gibco)<br>1% Penicillin/Streptomycin (Biowest)<br>0.05 mM β-Mercaptoethanol (Gibco) |

***Table S8. List of primary and secondary antibodies used in Western Blot experiments.***

| Primary antibody | Manufacturer | Catalog number | Dilution |
|---|---|---|---|
| **GSPT1** | Invitrogen | PA5-28256 | 1:1000 |

| | | | |
|---|---|---|---|
| **NEK7** | Abcam | ab133514 | 1:10000 |
| **CK1α** | Abcam | ab108296 | 1:2000 |
| **SALL4 (Kelly)** | Abcam | ab57577 | 1:1000 |
| **SALL4 (Hep3B)** | Santa Cruz Bio. | sc-101147 | 1:500 |
| **Ikaros** | Cell Signaling | 14859S | 1:2000 |
| **β-actin (HRP -conjugated)** | Abcam | ab20272 | 1:5000 (H929, Kelly)<br>1:10000 (Hep3B) |
| **Secondary antibody** | **Manufacturer** | **Catalog number** | **Dilution** |
| **anti-Rabbit IgG (HRP -conjugated)** | Invitrogen | 31466 | 1:2500 (For GSPT1)<br>1:10000 (For other Ab) |
| **anti-Mouse IgG (HRP -conjugated)** | Invitrogen | 31430 | 1:10000 |

*Degradation assays and Western Blot for VAP-1 dependent activity of ABS-752*

**Figure 5b:** During passage, after centrifugation Hep 3B2.1-7 cells with trypsin (Biowest, cat. L0931-500) and growth medium containing 10% FBS (Gibco) for 5 minutes 150 rcf, supernatant was discarded and cells were resuspended in the fresh growth medium containing 10% FBS, counted and at the appropriate density of cells were plated in 60x15 Petri dishes at density $0.44 \times 10^6$ cells/dish in a volume of 4.5 mL/dish of a complete growth medium containing 10% FBS, on the day before the compound treatment and incubated O/N in a 5% $CO_2$ incubator at 37ºC. Next day, the medium was discarded, cells were washed with DPBS (Biowest, cat. L0615-500) and fresh complete growth medium containing 10% FBS was added followed by 2 hours incubation in a 5% $CO_2$ incubator at 37ºC. After incubation, 1 hour pre-treatment with 10 µM concentration of PXS-4728A or DMSO (Miltenyi Biotec 170-076-303) was performed followed by 24 hours of compounds treatment. Stock solutions of compounds were prepared in 100% DMSO. The abundance of target proteins was analyzed after 25-hour incubation with compounds (1 h pre-incubation and 24 h co-treatment with tested compounds). Protein extracts as well as the other Western Blot steps were prepared as described in the Methods section. The POI signal values were normalized to the β-actin and Vinculin loading controls and relative POI levels were calculated as a percentage (%) of the DMSO control. The percentage downregulation/degradation values were calculated as 100% - % relative POI levels.

**Figure 5d:** During passage, after centrifugation Hep 3B2.1-7 cells with trypsin and growth medium containing 10% FBS for 5 minutes 150 rcf, supernatant was discarded and cells were resuspended in the sterile DPBS. Cells were centrifuged 150 rcf for 5 minutes, the supernatant was discarded and cells were resuspended in the sterile DPBS, counted and at the appropriate density of cells were centrifuged. After centrifugation, cells were resuspended in growth medium containing 10% FBS or 10% human serum AB male (Biowest, cat. S4190-100) and plated in 60x15 Petri dishes at density $0.44x10^6$ cells/dish and incubated O/N in a 5% $CO_2$ incubator at 37ºC. Next day, prior to compound treatment, media were changed for the fresh complete growth media followed by 24 hours of compounds treatment. Protein extracts as well as the other Western Blot steps were prepared as described in the Methods section. Secondary antibody (anti-Rabbit IgG HRP -conjugated, Invitrogen, cat. 31466) was used in 1: 10 000 dilution for both primary antibodies (GSPT1 and NEK7). The protein of interest (POI) signal values were normalized to the β-actin and Vinculin loading controls and relative POI levels were calculated as a percentage (%) of the DMSO control. The percentage downregulation/degradation values were calculated as 100% - % relative POI levels.

**Figure 5e:** Cell seeding was performed as described in Figure 5d. Next day, the medium was discarded and fresh complete growth medium containing 10% human serum was added followed by 1 hour pre-treatment with 10 μM concentration of PXS-4728A or DMSO. After pre-treatment, cells were treated with the compounds for 24 hours in a 5% $CO_2$ incubator at 37ºC. Protein extracts as well as the other Western Blot steps were prepared as described in the Methods section. Secondary antibody (anti-Rabbit IgG HRP -conjugated, Invitrogen, cat. 31466) was used in 1: 10 000 dilution for both primary antibodies (GSPT1 and NEK7). The protein of interest (POI) signal values were normalized to the β-actin and Vinculin loading controls and relative POI levels were calculated as a percentage (%) of the DMSO control. The percentage downregulation/degradation values were calculated as 100% - % relative POI levels.

##### *Tumors preparation for Western Blot analysis*

**Figure 8b:** Tumor samples were placed on dry ice and each was weighed on an analytical balance. Each tumor sample was dissected using a scalpel and transferred to homogenization tubes (MP Biomedicals, cat no. 115076400) containing ceramic spheres (MP Biomedicals, cat no. 116540422) which were placed on dry ice. Prepared samples were put on ice and to each sample appropriate volume of RIPA buffer prepared as described in the Methods section (Degradation assays and Western Blot) was added. After that, tubes were tightly closed with caps (MP Biomedicals, cat no. 115065005). The samples were kept on ice till the end of the homogenization and lysis.

The crushed dry ice was put to the adapter before use (MP Biomedicals, cat no. 6002528). The tubes containing tumor fragments prepared as described above were transferred to the cooled adapter and homogenized using FastPrep-24 Classic Homogenizer (MP Biomedicals, cat no. 6004500) with the following parameters: CY:24x2, 6.5 m/s; 45s; (5 minutes resting time for the homogenizer to cool down, samples stored on ice then). If some of the samples were non-homogenized completely, additional homogenization cycle was performed. Tumors fragments were maximally homogenized 6 times.

After homogenization, tumors were lysed on ice for 30 minutes (2-3 times vortexed during the incubation on ice). Lysates in homogenization tubes were centrifuged at 4°C, 15 000 x g, 10 minutes. After that, the supernatants were transferred to new microtubes (1.5 mL). To clarify the samples, supernatants were centrifuged at 4°C, 18 213 x g (maximum centrifuge speed), for 15 minutes. After centrifugation, the supernatants were transferred to 1.5 mL new microtubes and centrifuged once again at 4°C, 18 213 x g for 15 minutes. After centrifugation, supernatants were transferred to new 1.5 mL microtubes and tumor lysates were briefly placed into liquid nitrogen for snap freezing. Frozen lysates were thawed on ice and then centrifuged once more at 4°C, 18 213 x g for 15 minutes. Obtained supernatants were transferred to new 1.5 mL microtubes.

The protein concentrations in the tumors lysates were evaluated by BCA assay (according to the manufacturer's protocol). BCA assay was performed twice - in the first step samples were diluted to the 'working dilution' 10x and/or 20x. After the initial samples concentration measurement, samples were diluted to prepare stock concentration of approximately 5 mg/mL (5 µg/µL) of each sample. Then the samples were diluted 5x and measured in BCA assay again to evaluate the accurate concentration. Next steps were performed as described in the Methods section (Degradation assays and Western Blot). Primary antibody for GSPT1 (Invitrogen, cat. PA5-62621) was used in the dilution described in the Supplementary Section – Table S8.

#### CTG assay

***Table S9. The list of cell lines tested in CTG assay and the number of the cells seeded/well at 384-plate.***

| Cell line | Cell number/well at 384 well-plate |
|---|---|
| Hep 3B2. 1-7 | 1250 |
| JHH-7 | 1000 |
| HuH-1 | 1000 |
| HuH-7 | 750 |
| SNU-398 | 1500 |
| SNU-886 | 750 |
| Kelly | 2500 |
| SNU-423 | 375 |
| SNU-182 | 1250 |
| NCI-H929 | 3000 |

***Table S10. Cancer cell line information (Charles River Laboratories collection).***

| # | Cancer Type | Cell line name | Comment |
|---|---|---|---|
| 1 | Liver (Asian) | HLE | |

| | | | |
|---|---|---|---|
| 2 | Liver (Asian) | JHH-4 | |
| 3 | Liver (Asian) | JHH-6 | |
| 4 | Liver (Asian) | SNU-398 | |
| 5 | Liver (Asian) | SNU-423 | |
| 6 | Liver (Asian) | SNU-449 | |
| 7 | Liver (Asian) | SNU-475 | |
| 8 | Liver (Asian) | SNU-739 | |
| 9 | Liver (Asian) | SNU-761 | |
| 10 | Liver (Asian) | SNU-878 | |
| 11 | Liver | 575L (575) | Proprietary PDX derived cell line |
| 12 | Liver (Cholangiocellular) | EGI-1 | |
| 13 | Liver (Cholangiocellular) | KKU-M213 | |
| 14 | Liver | Hep3B | |
| 15 | Liver | HEP-G2 | |
| 16 | Liver | SK-HEP-1 | |
| 17 | Liver (Cholangiocellular) | TFK-1 | |

##### *CTG viability assay for VAP-1 dependent activity of ABS-752*

**Figure 5a:** Hep3B cells seeding was performed as described in Methods section. Next day, plate was spun down for 10 seconds at 100 rcf prior to 1 hour pre-treatment with 10 µM PXS-4728A, which was performed using Echo 555 Liquid Handler. After 10 µM PXS-4728A addition, plate was spun down for 10 seconds at 100 rcf and incubated in a 5% $CO_2$ incubator at 37°C. After pre-treatment, plate was spun down for 10 seconds at 100 rcf followed by the compounds addition performed using Echo 555 Liquid Handler. Each compound was tested in a serial dilution of 12 concentrations (0.1 nM - 30 µM) in a technical duplicate. All the compounds were dissolved in a 100% DMSO. For background subtraction, growth medium (no cells, n=12) was used. As negative control DMSO-treated cells or DMSO-treated cells with 10 µM PXS-4728A (n=11) were used. Final concentration of DMSO in all wells was 0.25%. After the compounds addition plate was spun down for 10 seconds at 100 rcf. Following the treatment, the cells were incubated for 72 h in a 5% $CO_2$ incubator at 37°C. Before the CTG viability assay was carried out, plate was removed from the incubator and equilibrated to room temperature (RT) for 15 min. CTG reagent was prepared and 12.5 µL was added to each well, contents mixed on an orbital shaker for 4 minutes at 460 rpm and the plate then incubated at RT in the dark w/o shaking for an additional 8 min to stabilize the signal. Luminescence signal was measured using CLARIOstar Multimode Plate Reader (BMG Labtech) using optical Fireflymode. The Focus and Gain were adjusted on a selected well of one of negative control (DMSO treated). Target value was set as 80%. Relative light unit (RLU) data were normalized and analyzed with a customized protocol. Normalization was based on the average of RLU of the negative control to calculate % viability. Dose response was assessed by non-linear regression and absolute pIC50 calculation with the average of technical duplicate of % viability.

**Figure 5c:** During passage, after centrifugation Hep 3B2.1-7 cells with trypsin and growth medium containing 10% FBS for 5 minutes 150 rcf, supernatant was discarded and cells were resuspended in the sterile DPBS. Cells were centrifuged 150 rcf for 5 minutes, the supernatant was discarded and cells were resuspended in the sterile DPBS, counted and at the appropriate density of cells were centrifuged. After centrifugation, cells were resuspended in growth medium containing 10% FBS or 10% human serum and seeded onto 384-well plate at density 1250 cells/well. Prepared plate was short centrifuged (10 seconds, 100 rcf) and placed in the 37°C, 5% $CO_2$ incubator overnight. Next day, the plate was spun down for 10 seconds at 100 rcf followed by the compounds addition performed using Echo 555 Liquid Handler. Each compound was tested in a serial dilution of 12 concentrations (0.1 nM - 30 μM) in a technical duplicate. All the compounds were dissolved in a 100% DMSO. For background subtraction, growth medium (no cells, n=6) was used and as negative control DMSO-treated cells (n=11) were used. Final concentration of DMSO in each well was 0.25%. After the compounds addition plate was spun down for 10 seconds at 100 rcf. Following treatment, the cells were incubated for 72h in a 5% $CO_2$ incubator at 37°C. CTG viability assay was performed as described in Methods section and Supplementary Figure 5a description.

**Figure 5f:** During passage, after centrifugation Hep 3B2.1-7 cells with trypsin and growth medium containing 10% FBS for 5 minutes 150 rcf, supernatant was discarded and cells were resuspended in the sterile DPBS. Cells were centrifuged 150 rcf for 5 minutes, the supernatant was discarded and cells were resuspended in the sterile DPBS, counted and at the appropriate density of cells were centrifuged. After centrifugation, cells were resuspended in growth medium containing 10% human serum and seeded onto 384-well plate at density 1250 cells/well on 384-well plate at 48 μL/well 24 h before the treatment. Prepared plate was short centrifuged (10 seconds, 100 rcf) and placed in the 37°C, 5% $CO_2$ incubator overnight. Prior to compound addition, 7.5 μM stock solution of recombinant human VAP-1 (PeproTech, cat. 150-16-250UG) was prepared in water for molecular biology (TH.GEYER, cat. 7711.1000) and added to appropriate wells at 2 μL/well. Content was mixed by gentle pipetting. Immediately after adding recombinant human VAP-1, compounds (prepared in 100% DMSO) were added directly to the plate using the Echo 555 Liquid Handler system. Final concentration of DMSO in all wells was 0.25%. Compounds were tested in technical triplicates. After 72 h incubation, assay plates were removed from the incubator and equilibrated to room temperature (RT) for 30 min. CTG reagent was prepared and 12.5 μL was added to each well, contents mixed on an orbital shaker for 4 minutes and the plates then incubated at RT in the dark for an additional 8 min. Luminescence signal was measured using CLARIOstar Multimode Plate Reader (BMG LABTECH). Normalization was based on the average of RLU of the 100% viability control (RLU 100%) and 0% viability control (RLU 0%) to calculate % viability.

Percentage of cell viability was calculated with the average of relative luminescence unit (RLU) of 100% viability control (RLU 100%) and 0% viability control (RLU 0%). 100% viability control was the average RLU of cells treated with 0.25% DMSO or treated with 0.25% DMSO with 0.3 µM VAP-1. 0% viability was the average of RLU from wells containing only medium or medium with 0.3 µM VAP-1 respectively. Dose response was assessed by non-linear regression and absolute pIC50 calculation with the average of technical triplicate of % viability. A single experiment was performed.

#### AlphaLISA-based ternary complex formation assay

***Table S11. Summary of AlphaLISA experimental conditions and reagents used.*** *Listed concentrations are those of the individual component in the final AlphaLISA reaction.*

| Target protein | CRBN/DDB1 complex | Acceptor beads | Donor beads | Buffer composition | Reference compound |
|---|---|---|---|---|---|
| GSPT1 [100 nM] | 100 nM | AlphaLISA Anti-6xHis Acceptor beads (PerkinElmer, AL178M) 20 µg/mL | Alphascreen Streptavidin Donor beads (PerkinElmer, 6760002) 20 µg/mL | 10 mM HEPES pH 7.4, 150 mM NaCl, 0.1% Tween-20, 1 mM DTT, 2% DMSO | CC-90009[2] |
| CK1α [20 nM] | 100 nM | AlphaLISA Anti-6xHis Acceptor beads (PerkinElmer, AL178M) 20 µg/mL | Alphascreen Streptavidin Donor beads (PerkinElmer, 6760002) 20 µg/mL | 20 mM phosphate buffer pH 7.4, 2.7 mM KCl, 137 mM NaCl, 0.01% Tween-20, 1 mM DTT, 2% DMSO | FPFT-2216[3] |
| NEK7 [100 nM] | 100 nM | AlphaLISA Anti-6xHis Acceptor beads (PerkinElmer, AL178M) 20 µg/mL | Alphascreen Streptavidin Donor beads (PerkinElmer, 6760002) 20 µg/mL | 10 mM HEPES pH 7.4, 150 mM NaCl, 0.1% Tween-20, 1 mM DTT, 2% DMSO | ABS-674* |
| SALL4 ZF2 [400 nM] | 100 nM | AlphaLISA Anti-6xHis Acceptor beads (PerkinElmer, AL178M) 20 µg/mL | Strep-Tactin Alpha Donor beads (PerkinElmer, AS106M) 20 µg/mL | 20 mM phosphate buffer pH 7.4, 2.7 mM KCl, 137 mM NaCl, 0.1% Tween-20, 1 mM DTT, 2% DMSO | Lenalidomide[4] |
| IKZF1 ZF2 [400 nM] | 100 nM | AlphaLISA Anti-6xHis Acceptor beads (PerkinElmer, AL178M) 20 µg/mL | Strep-Tactin Alpha Donor beads (PerkinElmer, AS106M) 20 µg/mL | 20 mM phosphate buffer pH 7.4, 2.7 mM KCl, 137 mM NaCl, 0.1% Tween-20, 1 mM DTT, 2% DMSO | Lenalidomide[5] |

* NEK7 degrader developed by Captor Therapeutics, structure is not disclosed.

#### Analysis of monoamine oxidases activity against ABS-752 using LC-MS/MS method

***Table S12. Tested compounds.*** *List of analytes with corresponding internal standards (IS). All tested compounds were dissolved in 100% DMSO at 20 mM (stock solution).*

| Compound ID | MW [Da] | Description |
|---|---|---|
| AAC-215 | 273.11 | IS for ABS-752 |
| ABR-321 | 315.12 | IS for ABT-003 |
| ABR-522 | 288.08 | IS for ABT-002 |
| ABT-394 | 272.08 | IS for ABT-971 |
| ABR-958 | 274.09 | IS for ABU-031 |
| ABS-752 | 291.1 | Analyte |
| ABT-003 | 306.06 | Analyte |
| ABT-002 | 333.11 | Analyte |
| ABT-971 | 290.07 | Analyte |
| ABU-031 | 292.09 | Analyte |

***Table S13. Materials and preparation of reagents.***

| **Reagent/solution** | **Composition/Manufacturer (Catalog No.)** |
|---|---|
| Methanol (MeOH) for LC-MS | ChemSolute (1203.2500) |
| Water LC-MS Chromasolv | Honeywell (39253-4L) |
| Formic acid (FA), 99%, for LC-MS | Serva (45640.01) |
| TCA, ACS reagent, ≥99.0% | Sigma-Aldrich (T6399-250G) |
| Liquid chromatography (LC) eluents | LC-MS grade water and MeOH with 0.1% FA (v/v) were used for the LC elution. 10% (v/v) MeOH in water was used as an autosampler needle wash |
| Extraction solvent | 1 M TCA (trichloroacetic acid) was prepared by weighing approximately 8.17 g of TCA and dissolving it in up to 50 mL of LC-MS grade water. 1 M TCA was stored at 4°C |
| Analytes and IS (internal standard) working solutions | Working solutions of each compound were prepared in water + 0.1% FA in a concentration of 100 µg/mL |
| Analytes and IS mix solutions | Mixes of analytes and IS were prepared in water + 0.1% FA in a final concentration of 10 µg/mL |
| Solvent blank | 1 mL of LC-MS grade MeOH + 0.1% FA was added to a glass vial and used as a solvent blank and column wash during LCMS analysis |
| Extraction solvent containing IS | IS mix was added to 1 M TCA to the final concentration of approx. 80 ng/mL (80 µL of IS mix for every 10 mL of 1 M TCA) |

***Table S14. LC-MS parameters.***

| | |
|---|---|
| **Instrument** | Thermo Scientific TSQ Altis Plus |
| **LC parameters** | |

| | |
|---|---|
| Column | Phenomenex Luna Omega PS C18, 1.6 µm, 50 x 2.1 mm |
| Column temp. | 30°C |
| Mobile phases | A: water + 0.1% FA<br>B: MeOH + 0.1% FA 0.4 mL/min |
| Flow | |
| Injection | 5 µL |
| Detection | MS/MS |
| Other parameters | 0.0 min 2% B, 0.5 min 5% B, 3.0 min 95% B, 3.5 min 95% B, 3.7 min 2% B, 5.5 min 2% B |
| **MS parameters** | |
| Ionization | positive/negative |
| Scan type | MRM (Multiple Reaction Monitoring) |
| Interface | ESI |
| Spray voltage | 4000 V |

***Table S15. MRM transitions.***

| Compound | Type | Polarity | Precursor (*m/z*) | Product (*m/z*) | Ion type | CE (V) |
|---|---|---|---|---|---|---|
| AAC-215 | IS (group 1) | + | 274.20 | 163.00 | Quantitative | -17 |
| | | | | 134.00 | Qualitative | -34 |
| | | | | 257.20 | Qualitative | -12 |
| ABR-321 | IS (group 2) | + | 316.05 | 205.10 | Quantitative | -17 |
| | | | | 146.05 | Qualitative | -33 |
| | | | | 243.20 | Qualitative | -24 |
| ABR-522 | IS (group 3) | - | 287.05 | 243.25 | Quantitative | 17 |
| | | | | 131.15 | Qualitative | 30 |
| | | | | 110.10 | Qualitative | 24 |
| ABT-394 | IS (group 4) | + | 273.20 | 162.00 | Quantitative | -15 |
| | | | | 199.95 | Qualitative | -25 |
| | | | | 145.20 | Qualitative | -26 |
| ABR-958 | IS (group 5) | + | 275.10 | 164.10 | Quantitative | -16 |
| | | | | 202.10 | Qualitative | -24 |
| | | | | 133.10 | Qualitative | -39 |
| ABS-752 | Analyte (group 1) | + | 292.20 | 275.15 | Quantitative | -13 |
| | | | | 164.05 | Qualitative | -25 |
| ABT-003 | Analyte (group 2) | + | 334.20 | 223.10 | Quantitative | -18 |
| | | | | 261.15 | Qualitative | -25 |
| ABT-002 | Analyte (group 3) | - | 305.10 | 261.20 | Quantitative | 17 |
| | | | | 241.10 | Qualitative | 26 |
| ABT-971 | Analyte (group 4) | + | 291.20 | 218.00 | Quantitative | -26 |
| | | | | 180.00 | Qualitative | -16 |
| ABU-031 | Analyte (group 5) | + | 293.20 | 182.05 | Quantitative | -15 |
| | | | | 220.15 | Qualitative | -25 |

#### Preparation of calibration curves:

Analytes mix (10 µg/mL) was diluted 4-times in water + 0.1% FA to give a concentration of 2500 ng/mL. Next, a dilution series in water + 0.1% FA was prepared and concentrations of 2500.00, 625.00, 156.25, 39.062, 9.76, 2.44, 0.61 ng/mL were used to prepare calibration curves. 25 µL of each calibration point were taken, added to 75 µL of 1 M TCA containing IS, and mixed. Next, samples were centrifuged (15.000 x g, 15 min, 10 °C). Each well of a 96-well microtiter plate (MTP) was prefilled with 75 µL of LC-MS grade water + 0.1% FA. Following centrifugation, 75 µL of clear supernatants were transferred to a MTP, mixed, spun down, and analyzed with LC-MS/MS. Calibration curves were prepared in duplicates.

**Sample processing from enzymatic reactions:**

Samples were thawed on ice, mixed and spun down. All samples were 3-fold diluted in LC-MS grade water prior to analysis. 25 µL of diluted samples were taken, added to 75 µL of 1 M TCA containing IS, and mixed. Next, samples were centrifuged (15.000 x g, 15 min, 10 °C). Each well of a 96-well MTP was prefilled with 75 µL of LC-MS grade water + 0.1% FA. Following centrifugation, 75 µL of clear supernatants were transferred to a MTP, mixed, spun down, and analyzed with LC-MS/MS.

**Data analysis:**

Raw LC-MS data was analyzed with the Chromeleon software (Thermo Fisher Scientific), to generate the primary data (peak areas and compounds concentrations). Next, Excel (Microsoft) was used to validate and visualize the data (Appendix_1).

#### Analysis of ABS-752 metabolites formation in whole human tissue lysates using LC-MS/MS method

***Table S16. The ABS-752 dilutions used in the experiment.***

| # | 1.1x stock of ABS-752 [µM] | ABS-752 final concentration [µM] |
|---|---|---|
| 1 | 2.17 | 1.95 |
| 2 | 4.34 | 3.91 |
| 3 | 8.68 | 7.81 |
| 4 | 17.36 | 15.63 |
| 5 | 34.73 | 31.25 |
| 6 | 69.45 | 62.5 |
| 7 | 138.9 | 125 |
| 8 | 277.8 | 250 |
| 9 | 555.6 | 500 |
| 10 | 1111.2 | 1000 |

**Methodology:**

Quantification of ABS-752 analyte and its metabolites (ABT-003, ABT-002, ABT-971, ABU-031) was performed using ACQUITY UPLC system (Waters) coupled with Xevo TQ-S mass spectrometer (Waters). Chromatographic separation was achieved using Acquity UPLC BEH Shield RP18 1.7 µm, 5 cm (Waters) column. AAC-215, ABR-321, ABR-958 and ABR-522 analytes were used as an internal standard. Supplementary Table S17 presents chromatographic gradient and mobile phases composition used to separate analytes.

Mass spectrometer was run in MRM mode (*multiple reaction monitoring*) with positive and negative ESI. The ion transmission pairs selected for the quantification were presented in Supplementary Table S18.

Sample preparation:

(1) All samples were stored at -80°C and thawed in a fridge (4°C) until defrosted prior to analysis.
(2) Calibrators, quality controls and experimental samples were centrifuged for 10 min at 1500 RPM.
(3) 30 µL of each sample was put into a previously assigned well.
(4) 100 µL of internal standard solution mix (20 ng/mL of AAC-215, ABR-321, ABR-958 and ABR-522) in 1 M TCA (Trichloroacetic acid) was added to each well using Repetman electronic repetitive pipette (Gilson).
(5) Plate was covered with a silicone mat and mixed for 5 min.
Next, 100 µL of 0.1% FA (Formic acid) in water was added to each well.
(6) Plate was covered with a silicone mat and mixed for 5 min.
(7) Finally the plate was spun down for 5 min at 4000 RPM before being placed into the autosampler.

***Table S17. Chromatographic gradient used in the separation of the analytes. Table includes composition of mobile phases, oven temperature and total method run time.***

| Mobile phase A | | 2 mM AmmFA + 0.1 FA in water | |
|---|---|---|---|
| Mobile phase B | | 2 mM AmmFA + 0.1 FA in methanol | |
| Oven temperature | | 55°C | |
| Total run time | | 2.5 min | |
| **Time [min]** | **Flow [mL/min]** | **% A** | **% B** |
| 0.00 | 0.50 | 98 | 2 |
| 0.50 | 0.50 | 98 | 2 |
| 1.00 | 0.50 | 85 | 15 |
| 2.00 | 0.50 | 0 | 100 |
| 2.50 | 0.50 | 0 | 100 |
| 2.51 | 0.50 | 98 | 2 |

***Table S18. Mass to charge ratio (m/z) for analyte and IS transmissions.***

| | | *m/z* | | |
|---|---|---|---|---|
| **Analyte** | **ESI mode** | **Parental ion** | **Quantifier** | **Identifier** |
| ABS-752 | + | 292.20 | 181.07 | 275.10 |
| ABT-003 | + | 334.20 | 164.06 | 223.14 |
| ABT-002 | - | 305.10 | 241.10 | 261.10 |
| ABT-971 | + | 291.20 | 84.00 | 218.10 |
| ABU-031 | + | 293.20 | 182.10 | 220.10 |
| **Internal standard** | | | | |
| AAC-215 (IS for ABS-752) | + | 274.20 | 134.00 | 163.10 |
| ABR-321 (IS for ABT-003 and ABT-971) | + | 316.05 | 205.10 | 146.10 |
| ABR-958 (IS for ABU-031) | + | 275.10 | 164.10 | 202.10 |
| ABR-522 (IS for ABT-002) | - | 287.05 | 243.10 | 131.10 |

**Calibration curve range and linearity:**

Calibration curves for analytes were prepared using methanol (HPLC grade). The calibration curve range was from 1 ng/mL to 2000 ng/mL. Results for calibration curve linearity are presented in Supplementary Figures S20-S24.

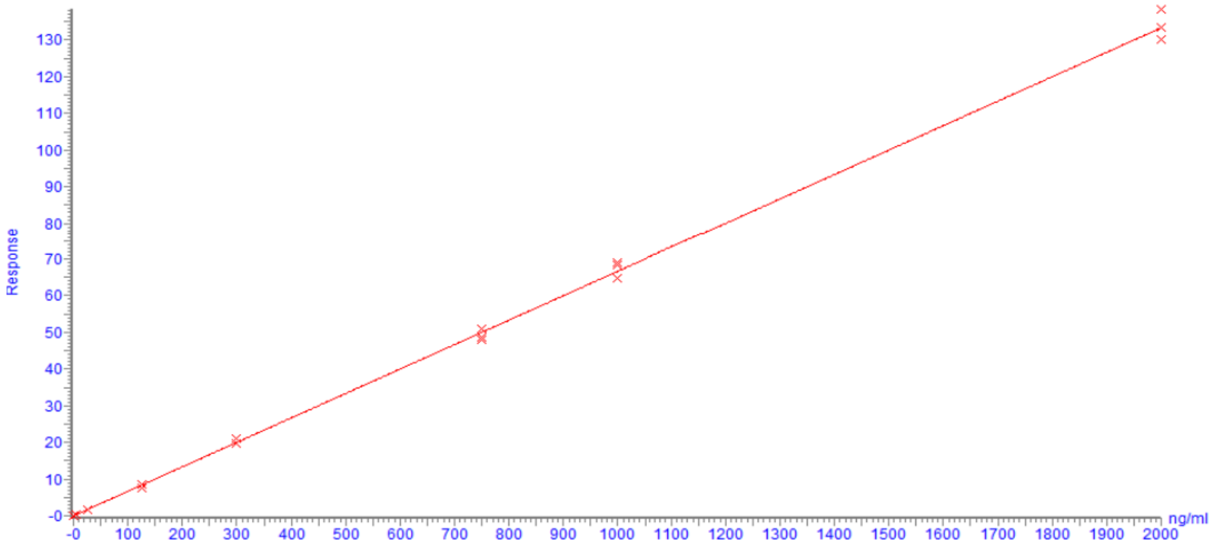

***Figure S20. Graphical representation of the linearity of 3 calibration curves for compound ABS-752.*** *Correlation coefficient factor is equal to 0.994.*

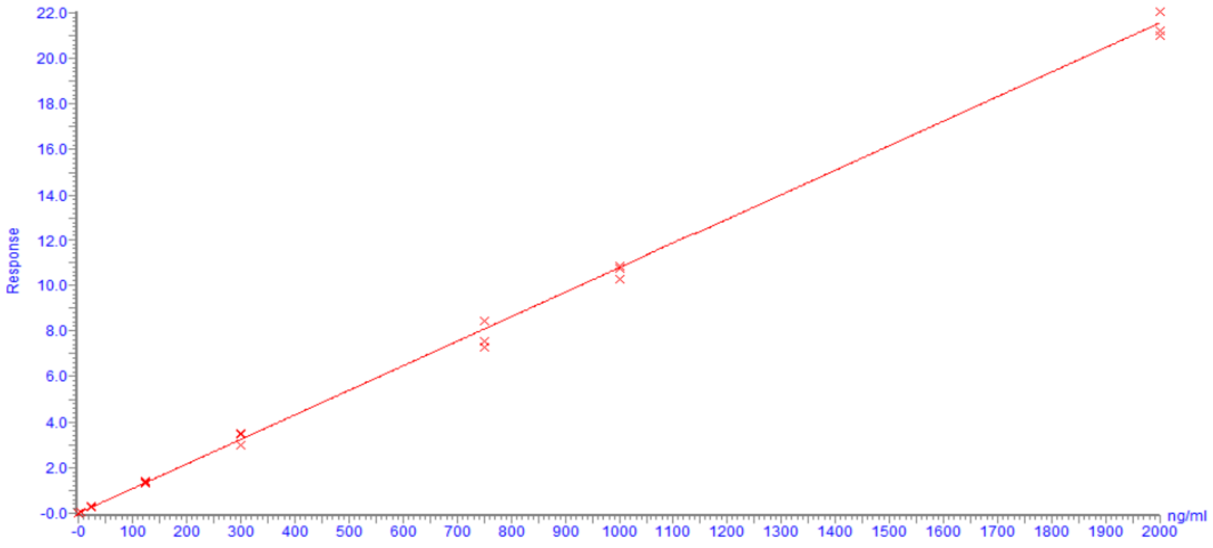

***Figure S21. Graphical representation of the linearity of 3 calibration curves for compound ABT-003.*** *Correlation coefficient factor is equal to 0.996.*

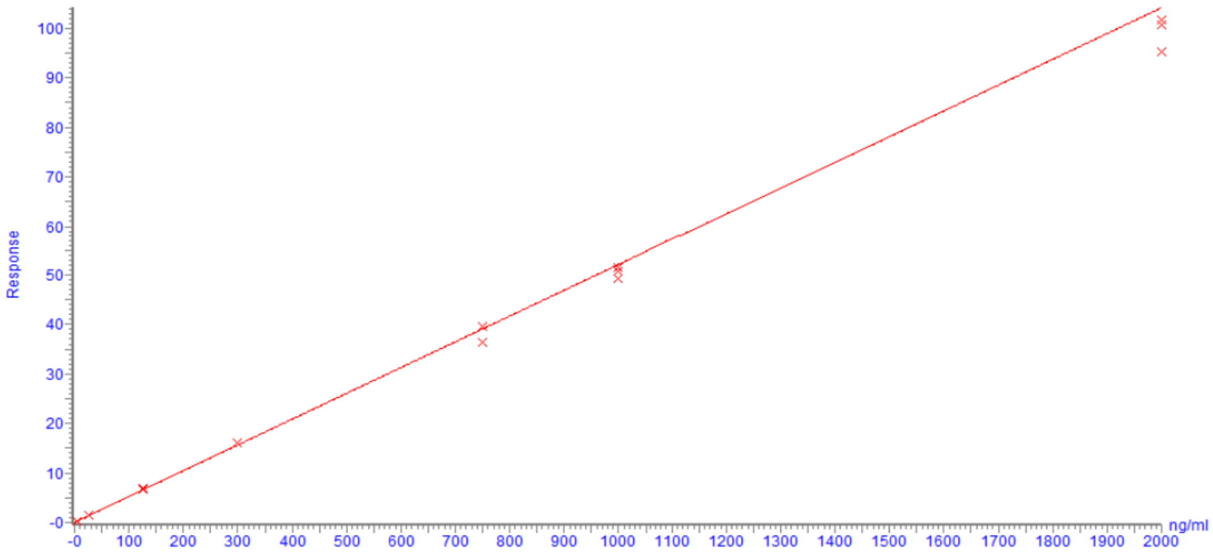

***Figure S22. Graphical representation of the linearity of 3 calibration curves for compound ABT-002.*** *Correlation coefficient factor is equal to 0.997.*

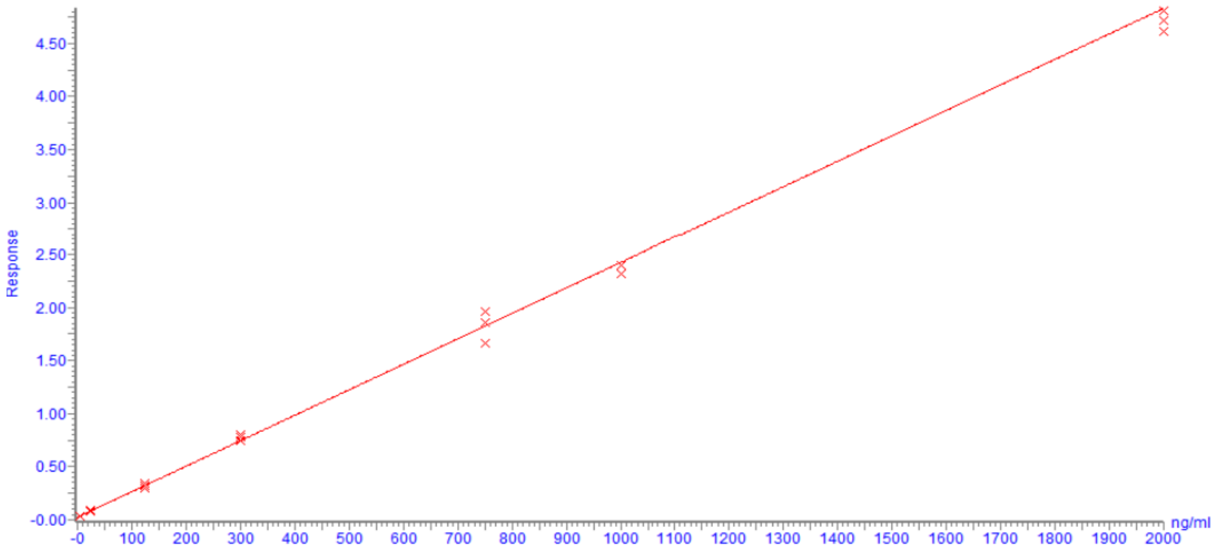

***Figure S23. Graphical representation of the linearity of 3 calibration curves for compound ABT-971.*** *Correlation coefficient factor is equal to 0.997.*

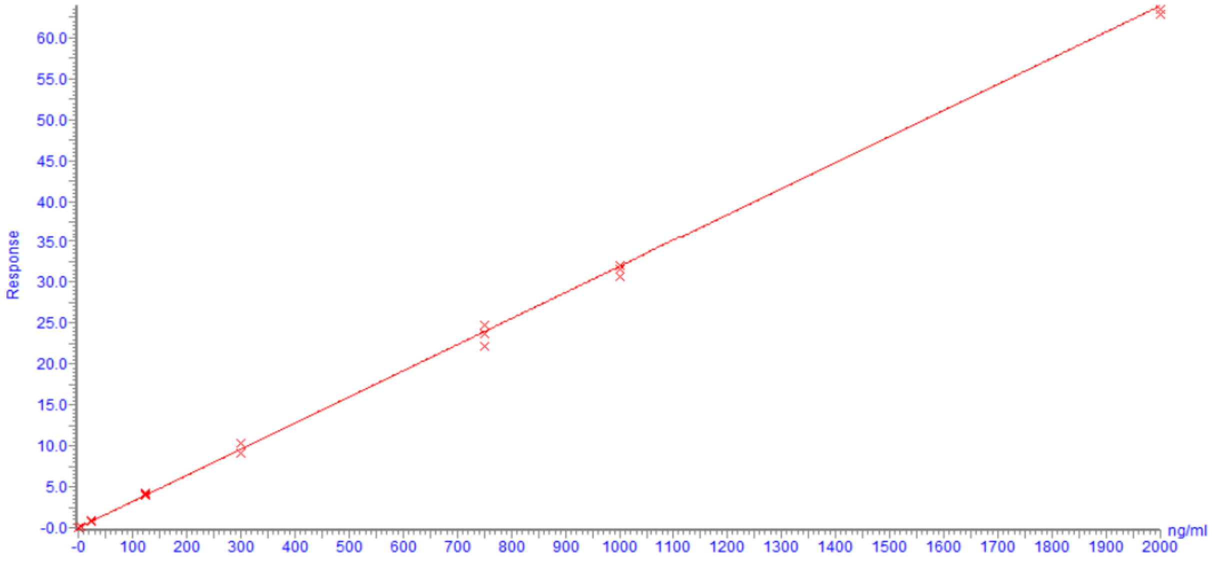

***Figure S24. Graphical representation of the linearity of 3 calibration curves for compound ABU-031.*** *Correlation coefficient factor is equal to 0.997.*

**Repeatability:**

A repeatability test was performed. Three quality controls were prepared with concentration levels reflecting different calibration curve ranges (low (QC I), medium (QC II) and high (QC III)). Repeatability results for each analyte are listed in the tables in Appendix_3.

**Chromatograms:**

Chromatograms of all the quantified analytes with their respective internal standard (IS) are given below:

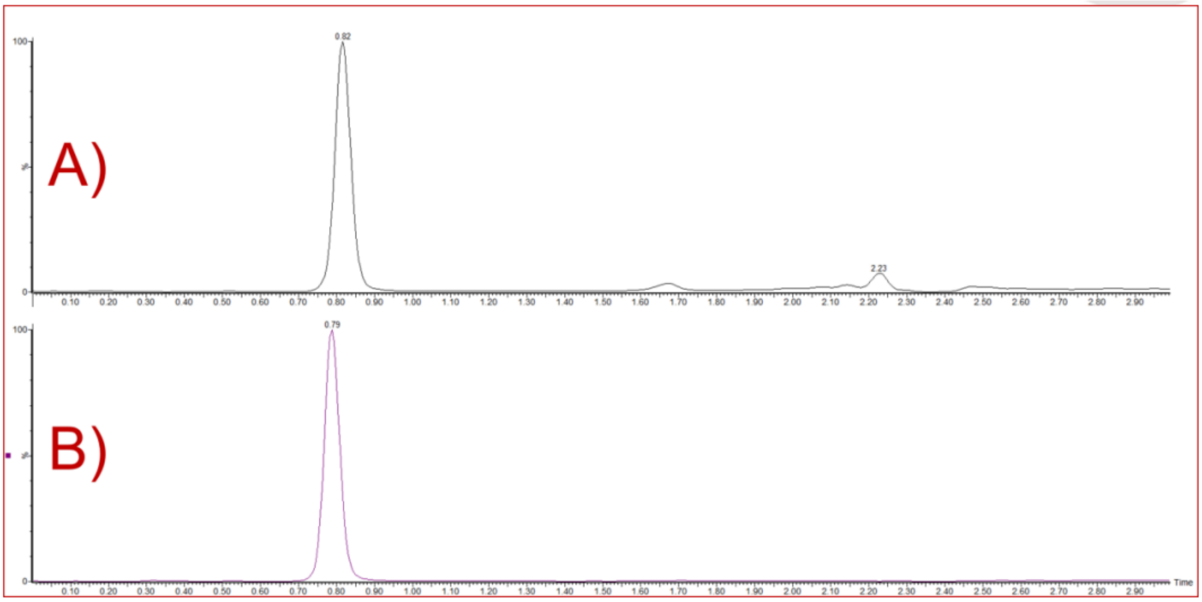

***Figure S25. Chromatogram for ABS-752 and its respective internal standard (AAC-215). A - peak for ABS-752; B – peak for AAC-215.***

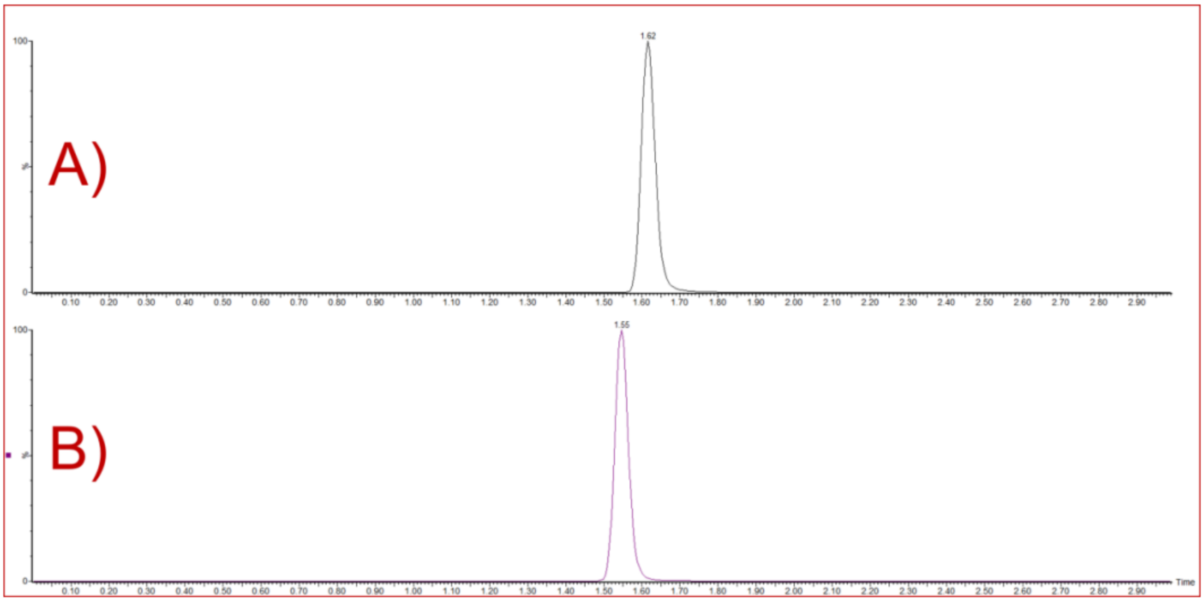

***Figure S26. Chromatogram for ABT-003 and its respective internal standard (ABR-321). A - peak for ABT-003; B – peak for ABR-321.***

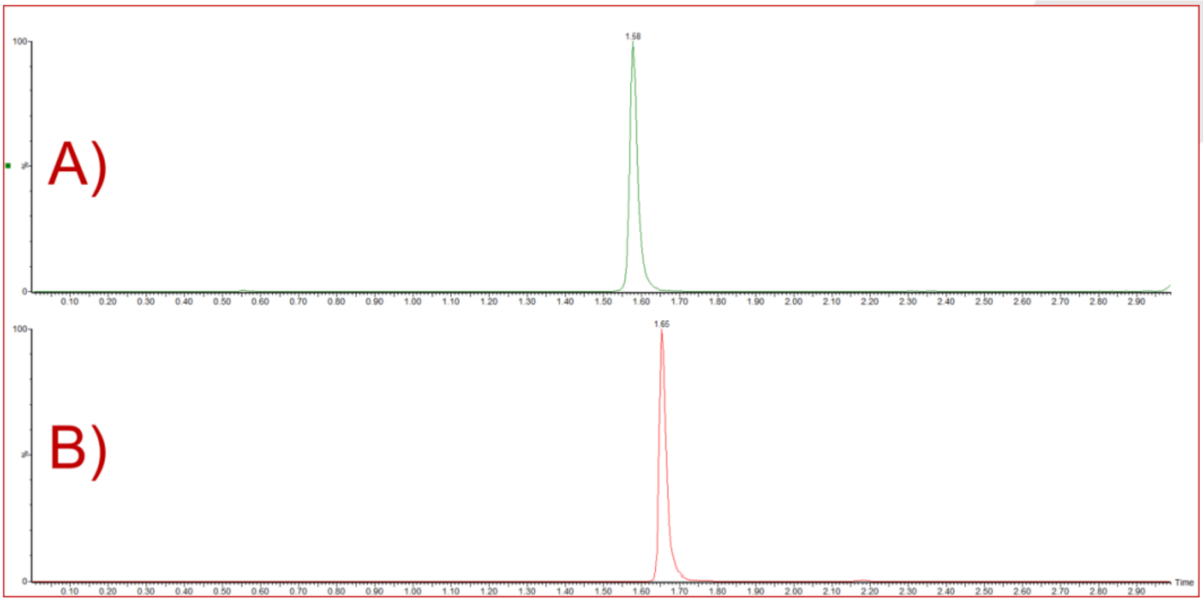

***Figure S27. Chromatogram for ABT-002 and its respective internal standard (ABR-522). A - peak for ABT-002; B – peak for ABR-522.***

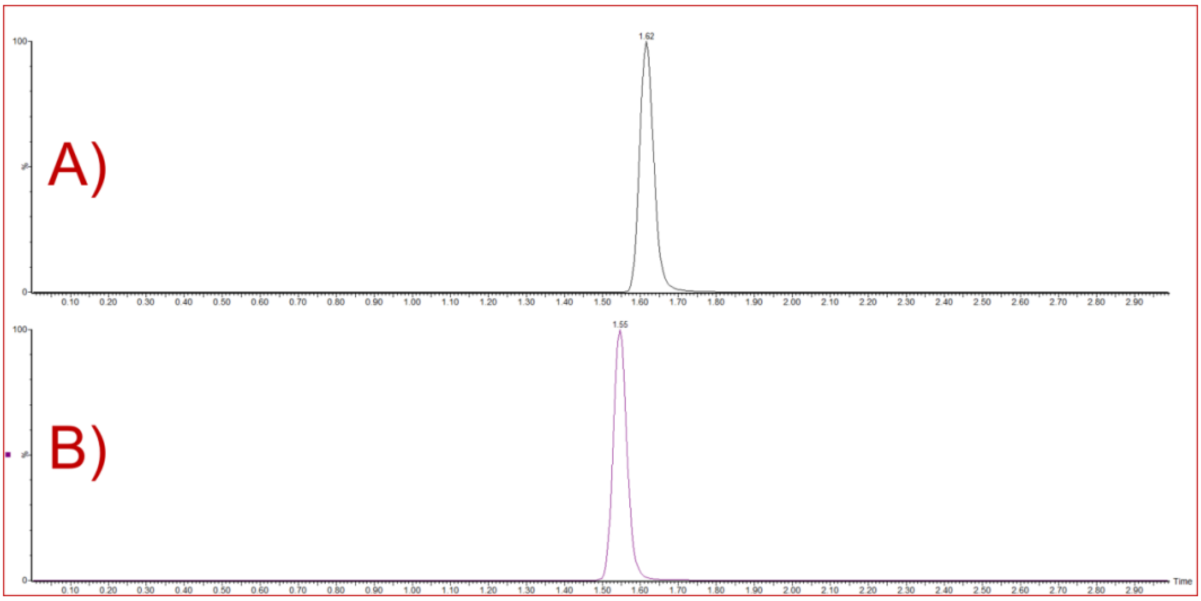

***Figure S28. Chromatogram for ABT-971 and its respective internal standard (ABR-321). A - peak for ABT-971; B – peak for ABR-321.***

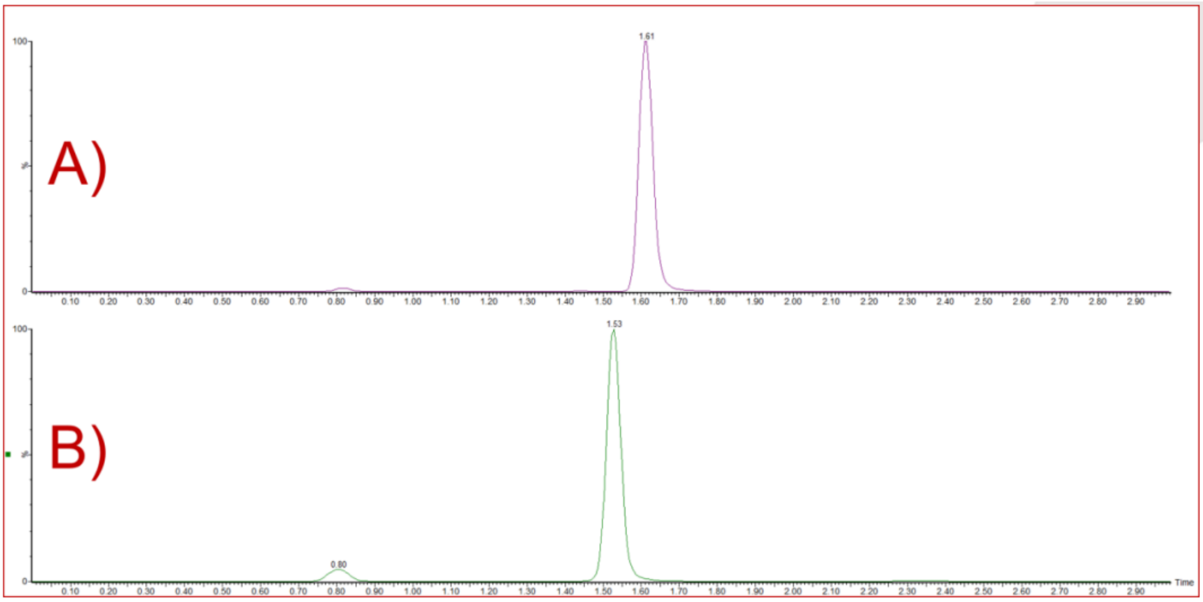

***Figure S29. Chromatogram for ABU-031 and its respective internal standard (ABR-958). A - peak for ABU-031; B – peak for ABR-958.***

#### Generation of the Hep3B G575N mutant cell line using the CRISPR/Cas9 system

Human GSPT1 gene sequence (genome version GRCh38p13) was obtained from NCBI Gene. The following gRNA sequence was identified in the vicinity of the GSPT1 G575 (5'->3'): TGCTTGGTAGACAAAAAATC with the PAM sequence: AGG. The gRNA sequence was cloned into the plasmid encoding Cas9 protein and containing Puromycin resistance gene. As a repair template, the following ssODN (single-strand oligo donor) containing: the mutation gga->aac (G575N), 50-bp flanking homology sequence and 2 phosphorothioate bonds (marked with "*") was used (5'->3'): t*t*ctcatactcttttcttggttaggccttaatctgcttggtagacaaaaaatcaaacgaaaaaagtaagacccgacccc gttttgtcaaacaagatcaagtatgc*a*t. 25x$10^4$ Hep3B cells (ATCC, clone Hep3B2.1-7) were electroporated in a 100 µL of electroporation reaction containing 2.5 µg of gRNA plasmid and 200 pmol of the repair template using the Neon Transfection System (Thermo Fisher Scientific, #MPK5000) under the following conditions: Pulse voltage -1275 V, Pulse width – 40 ms, Number of pulses – 1. 24 hours after electroporation, the cells were subject to Puromycin selection (2 µg/ml final concentration) for 48 hours, followed by clonal selection by limited dilution. GSPT1 sequence flanking the mutation site of the resulting clones was amplified by PCR with the following primers: forward (5'->3'): GTTTGACAAAACGGGGTCGG and reverse (5'->3'): TCTGCCCAGGCTATAATGCG. The PCR product was sequenced using the following primer (5' -> 3'): TCTGCCCAGGCTATAATGC.

Three clones containing the homozygous mutation gga->aac (G575N) were identified. Clone 16 was used in the experiments described in this study.

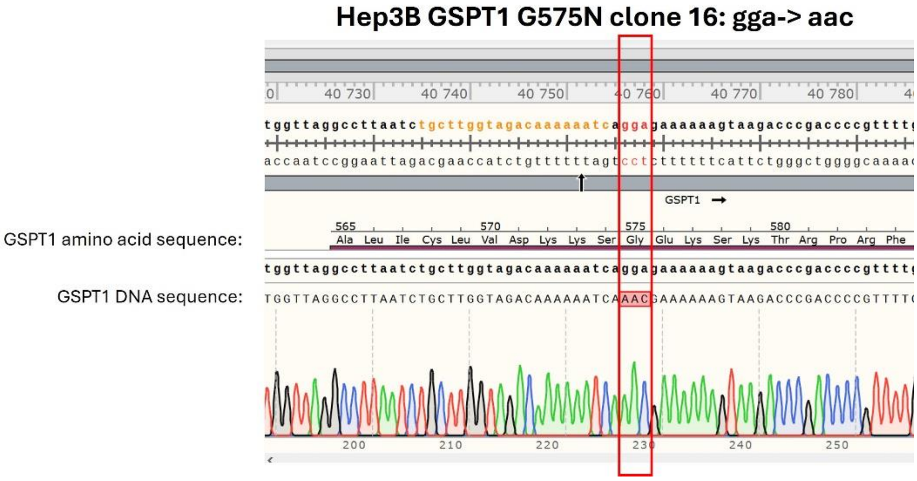

***Figure S30. Sequencing electropherogram showing the homozygous G575N mutation in GSPT1 G575N Hep3B clone 16.***
